# Dfm1 promotes Ste24-dependent translocon quality control

**DOI:** 10.64898/2026.07.01.735875

**Authors:** James A. Avaala, Sophia L. Owutey, Connor G. Bailey, Kathryn L. Alexander, Karen E. Alexander, Melissa Palackel, Travis S. Beckett, Katrina A. Procuniar, Kyle A. Richards, Kelsey A. Woodruff, Katie M. Lehman, Kaikeyi M. Paxton, Sudheeksha Bommineni, Emmanuel Akoto, Samantha M. Turk, Jacob M. Miller, Hailey J. Barton, Eric S. Fults, Robert J. Tomko, Eric M. Rubenstein

## Abstract

Proteins stalled during endoplasmic reticulum translocation clog Sec61 channels and require translocon quality control (TQC) mechanisms for clearance. Genetic screens implicated the derlin Dfm1 in TQC; however, its mechanistic role remained unclear. Using engineered and disease-relevant translocon-clogging substrates in *Saccharomyces cerevisiae*, we show Dfm1 functions in the Ste24-dependent branch of TQC, but not Hrd1- or Ltn1-dependent pathways. Loss of *DFM1* increased accumulation and toxicity of translocon-clogging proteins. Increased dosage of wild type or catalytically inactive Hrd1 or Ste24 rescued toxicity, supporting a non-enzymatic buffering function for these factors in TQC. These findings identify Dfm1 as a key TQC factor whose critical role in clearing clogged translocons can be suppressed in multiple ways. Our results suggest enhancing TQC may protect cells from proteotoxic stress caused by disease-relevant translocon-clogging substrates, with potential implications for pancreatic β-cell dysfunction in diabetes.

**Significance Statement:**

- Cells rely on translocon quality control (TQC) to clear proteins that clog the Sec61 translocon, but how the derlin Dfm1 contributes to this process has remained unclear.
- We demonstrate that Dfm1 functions specifically in the Ste24-dependent TQC pathway and show that increased levels of the TQC factors Hrd1 or Ste24 rescues Dfm1 deficiency even without catalytic activity, revealing an unexpected non-enzymatic buffering function.
- This work defines Dfm1’s mechanistic role in TQC, provides new insight into how cells preserve endoplasmic reticulum function during translocation stress, and establishes a framework for investigating conserved pathways that protect against diseases associated with translocon clogging.

## Introduction

Approximately one third of eukaryotic proteins enter the endoplasmic reticulum (ER) en route to their final destination (Chen *et al*., 2005; Choi *et al*., 2010). Many of these proteins cross the ER membrane by passing through the Sec61 translocon (Itskanov and Park, 2023). Occasionally, proteins stall during ER translocation, producing clogged, functionally compromised Sec61 channels (Rubenstein *et al*., 2012; Ast *et al*., 2016; Wang and Ye, 2020). Cells possess multiple conserved translocon quality control (TQC) mechanisms to resolve these clogged channels (Wang and Ye, 2020). Several of these mechanisms were discovered and initially characterized in *Saccharomyces cerevisiae*. The zinc metalloprotease Ste24 (human homolog, ZMPSTE24) and ubiquitin ligase Hrd1 (human homologs, gp78 and HRD1) exhibit partially redundant function in resolving translocons clogged by proteins that engage translocons post-translationally (Rubenstein *et al*., 2012; Ast *et al*., 2016; Runnebohm *et al*., 2020). By contrast, the ubiquitin ligase Ltn1 (also known as Rkr1; human homolog, Listerin) degrades proteins that clog translocons due to arrested translation, as an extension of its role in ribosome quality control (Crowder *et al*., 2015; von der Malsburg *et al*., 2015; Arakawa *et al*., 2016; Ennis *et al*., 2025). Additional layers of TQC exist in mammals, including marking of translationally stalled, ER-bound ribosomes by the ubiquitin-like modifier UFM1 (Wang *et al*., 2020; Scavone *et al*., 2023).

Two genetic screens implicated the derlin Dfm1 in TQC (Ast *et al*., 2016; Kayatekin *et al*., 2018). Derlins are ER-resident rhomboid pseudoproteases with diverse roles in ER protein quality control (PQC), lipid biosynthesis, and organellar homeostasis (Lemberg, 2013; Kandel and Neal, 2020; Bhaduri *et al*., 2023a; Bhaduri *et al*., 2023b; Scott *et al*., 2026). Yeast encode two derlins, Der1 and Dfm1, while mammals encode Derlin-1, -2, and -3 (Bhaduri *et al*., 2023b). Dfm1 and mammalian Derlin-1 and Derlin-2 (but not yeast Der1 or mammalian Derlin-3) possess C-terminal SHP motifs that recruit the AAA ATPase Cdc48/p97. Cdc48 powers retrotranslocation of transmembrane ER proteins to the cytosol (Sato and Hampton, 2006; Goder *et al*., 2008; Stolz *et al*., 2010; Lim *et al*., 2016).

Whereas Der1 promotes retrotranslocation of aberrant ER luminal proteins for degradation (Knop *et al*., 1996; Carvalho *et al*., 2006; Horn *et al*., 2009; Wu *et al*., 2020), Dfm1’s role in ER PQC had been controversial. Early studies reported limited or no contribution of Dfm1 to ER PQC (Sato and Hampton, 2006; Goder *et al*., 2008), but later work showed Dfm1 promotes retrotranslocation and reduces toxicity of transmembrane PQC substrates of the ER ubiquitin ligases Hrd1 and Doa10 (Stolz *et al*., 2010; Neal *et al*., 2018; Vitali *et al*., 2024). Earlier negative findings have been explained by rapid genetic suppression in *dfm1*Δ cells, often via *HRD1* gene amplification and remodeling of the Hrd1 complex to create an alternative retrotranslocation route (Neal *et al*., 2018; Neal *et al*., 2020).

Genome-wide studies showed loss of *DFM1* impairs growth of yeast expressing the engineered translocon-clogging substrate Clogger (Figure 1A) (Ast *et al*., 2016) and sensitizes cells to a yeast model for the translocon-obstructing oligomeric islet amyloid polypeptide (IAPP) (Figure 3A) (Kayatekin *et al*., 2018). However, Dfm1’s mechanistic role in TQC remained undefined. Here, using defined translocon-clogging proteins targeted by Ste24, Hrd1, or Ltn1, we show that Dfm1 clears translocons and mitigates toxicity of proteins handled primarily by the Ste24 pathway, but not by Hrd1- or Ltn1-dependent mechanisms. We further show that increased dosage of wild type or catalytically inactive Hrd1 or Ste24 rescues toxicity of translocon-clogging proteins in *dfm1*Δ yeast. Rescue is accompanied by increased frequency of successful translocation, but not removal of ER translocon-clogging species, suggesting non-enzymatic functions for Ste24 and Hrd1 in relieving clogged translocons. Our findings identify a critical, but suppressible, role for Dfm1 in TQC and provide new mechanistic insight into how cells relieve translocation stress.

**Figure 1.**
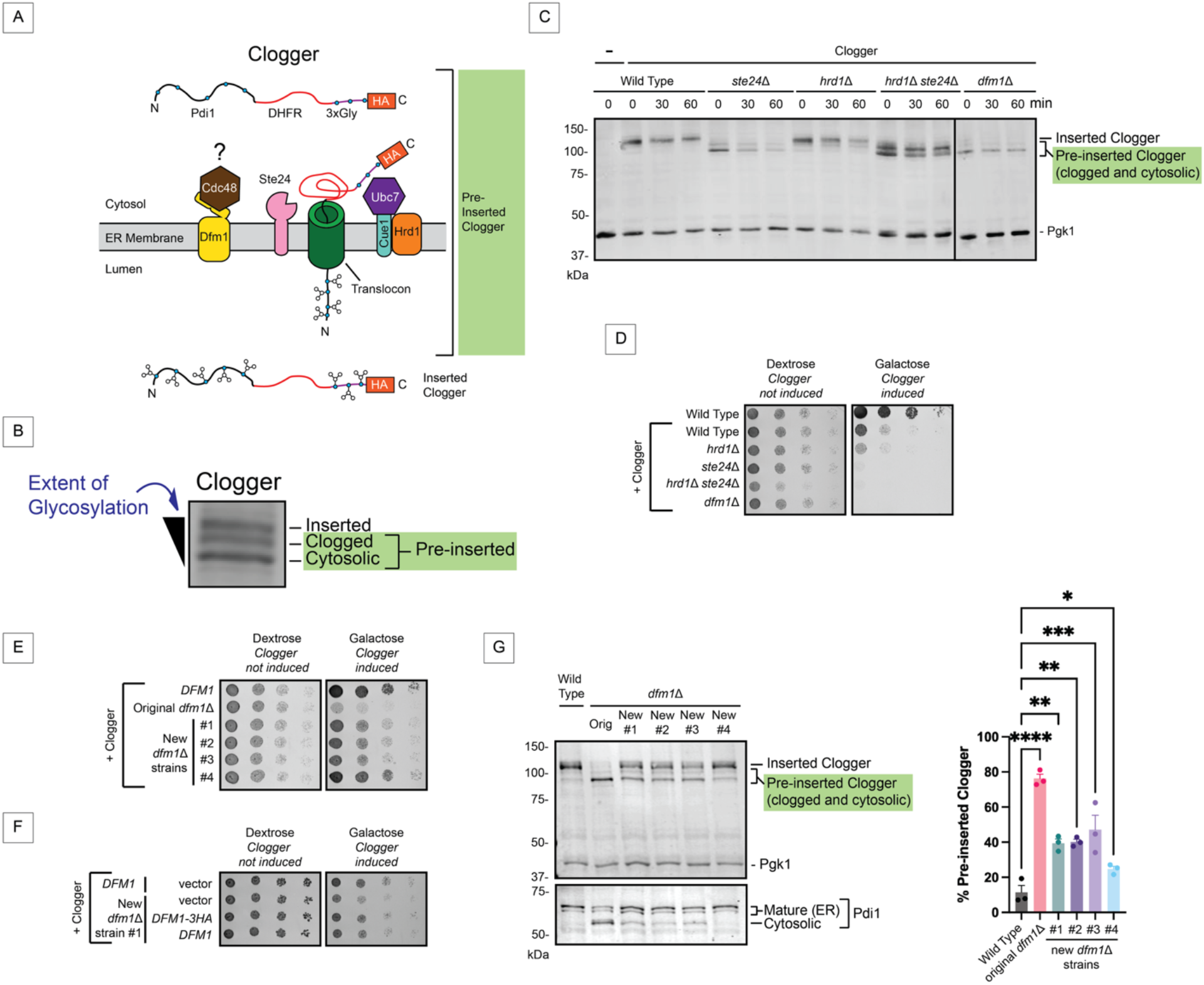
Dfm1 is required for TQC. (A) Clogger is an engineered translocon-clogging protein targeted by Ste24 and the Hrd1 complex (Hrd1, Ubc7, Cue1). Clogger consists of the post-translationally translocated ER glycoprotein Pdi1; the rapidly folding, translocon-clogging *E. coli* dihydrofolate reductase (DHFR); additional glycosylation sites; and an HA tag. Blue circles, glycosylated amino acids. (B) Western blot of Clogger. (C) Cycloheximide chase of Clogger. (D-F) 6-fold dilutions of yeast spotted onto medium containing dextrose or galactose. Yeast in (F) were transformed with empty vector or plasmid encoding Dfm1(−3HA). (G) Left, western blot of Clogger and Pdi1 in steady state extracts. Right, means of % Clogger that is pre-inserted were compared by one-way ANOVA with Tukey’s multiple comparison test. *, *p* < 0.05. **, *p* < 0.01. ***, *p* < 0.001. ****, *p* < 0.0001. (C, G) Pgk1, loading control. All experiments were conducted in triplicate.

## Results and Discussion

### Dfm1 is required for Translocon Quality Control

We obtained the *dfm1*Δ strain generated in the original Clogger degradation screen (Ast *et al*., 2016). Clogger glycosylation status and electrophoretic mobility reflect its position relative to the ER membrane (Figure 1B). The slowest migrating, fully glycosylated species have been fully translocated into the ER lumen. Intermediate-mobility, hemi-glycosylated species are translocon-clogged Clogger. The fastest migrating, unglycosylated species are cytosolic, although their efficient interaction with Ste24 (Ast *et al*., 2016) suggests they may represent early ER-associated or translocation intermediates. The clogged and cytosolic species (collectively, “pre-inserted” Clogger) accumulate in TQC-deficient cells, and their abundance correlates with reduction in fitness.

To assess the role of Dfm1 in TQC, we analyzed Clogger dynamics by cycloheximide chase in wild type yeast, strains lacking established TQC factors Hrd1 and Ste24, and *dfm1*Δ cells (Figure 1C). In wild type cells, Clogger was predominantly observed in its fully translocated form. As previously reported, *ste24*Δ yeast accumulated pre-inserted Clogger, while *HRD1* deletion modestly stabilized the clogged form. Combined loss of *STE24* and *HRD1* further increased pre-inserted Clogger, consistent with overlapping roles in resolving clogged translocons (Runnebohm *et al*., 2020). Notably, *dfm1*Δ cells also showed pronounced stabilization of pre-inserted Clogger. The distribution of Clogger species in *dfm1*Δ cells closely resembled that of *ste24*Δ cells, suggesting Dfm1 and Ste24 function in the same pathway.

We next compared the sensitivity of wild type and TQC mutant strains to translocon-clogging stress (Figure 1D). Clogger expression is driven by the *GAL1* promoter. Loss of *DFM1* did not impair fitness in the absence of Clogger expression (dextrose). As previously observed (Runnebohm *et al*., 2020), *HRD1* and *STE24* exhibited a synthetic negative interaction. Clogger induction reduced fitness in wild type cells. *HRD1* deletion modestly enhanced Clogger sensitivity. Loss of *STE24* or *DFM1* substantially exacerbated this defect, consistent with impaired clearance of toxic pre-inserted Clogger species.

Because the *dfm1*Δ strain analyzed above was recovered directly from the original genetic screen and harbors additional screen-specific alterations (Ast *et al*., 2016), we generated independent Clogger-expressing *dfm1*Δ strains for validation. Given frequent genetic suppression reported in *dfm1*Δ yeast (Neal *et al*., 2018; Neal *et al*., 2020), we deleted *DFM1* in cells harboring a plasmid-borne copy of *DFM1*, followed by plasmid eviction.

Three independent *dfm1*Δ isolates showed pronounced sensitivity to clogging stress (Figure 1E), although less severe than the screen-derived strain. A fourth isolate showed only a modest fitness defect, suggesting a potential suppressor mutation. Plasmid-based expression of *DFM1* restored resistance to clogging stress, and a C-terminal 3HA tag did not impair complementation (Figure 1F). Importantly, accumulation of pre-inserted Clogger (Figure 1G) correlated with growth impairment. Consistent with translocon blockage, loss of *DFM1* also impaired ER import of endogenous Pdi1 in proportion to Clogger sensitivity and pre-inserted species accumulation (Figure 1G, bottom panel). Together, these results establish *DFM1* as a bona fide TQC factor.

### Dfm1 functions in the Ste24 TQC pathway

Hrd1 and Ste24 mediate partially redundant TQC mechanisms. Clogger species distribution and sensitivity in *dfm1*Δ cells more closely resembled *ste24*Δ than *hrd1*Δ cells (Figures 1C and 1D). Moreover, *ste24*Δ *dfm1*Δ double mutants did not show increased Clogger sensitivity (Figure 2A) or accumulation of pre-inserted Clogger relative to either single mutant (Figure 2B), unlike the additive defects observed in *hrd1*Δ *ste24*Δ yeast (Figure 1C, (Runnebohm *et al*., 2020)). These results suggest Dfm1 functions in the Ste24-dependent pathway.

**Figure 2.**
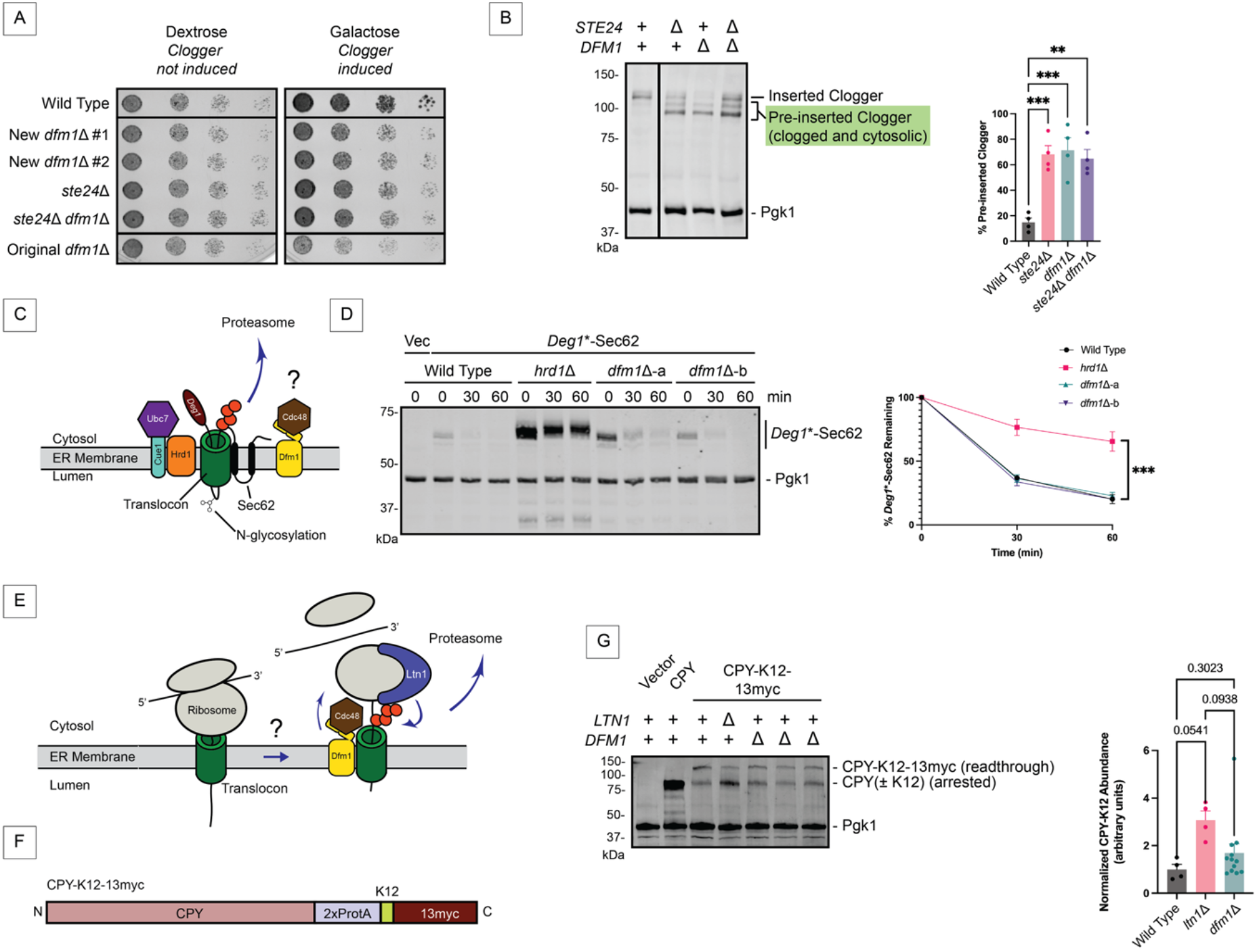
Dfm1 functions in Ste24-dependent TQC. (A) 6-fold dilutions of yeast spotted onto medium containing dextrose or galactose. (B) Left, western blot of Clogger in steady state extracts. Right, means of % Clogger that is pre-inserted in 4 biological replicates were compared by one-way ANOVA with Tukey’s multiple comparison test. **, *p* < 0.01. ***, *p* < 0.001. (C) Following insertion of 2 typical transmembrane segments, a portion of the N-terminal tail of *Deg1\**-Sec62 loops into and persistently engages the ER translocon. *Deg1*\*-Sec62 is targeted primarily by Hrd1. (D) Left, cycloheximide chase of *Deg1*\*-Sec62. Right, means of % *Deg1*\*-Sec62 remaining over time plotted for 3 biological replicates. Means of % *Deg1*\*-Sec62 remaining at 60 min were compared by one-way ANOVA with Tukey’s multiple comparison test. ***, *p* < 0.001 and applies to comparison between *hrd1*Δ yeast and all 3 other strains. (E) Translationally arrested ER-targeted proteins are targeted by Ltn1-mediated ribosome quality control. (F) CPY-K12-13myc is a model ER-targeted Ltn1 substrate. Left, western blot of CPY-K12 (translationally stalled) and CPY-K12-13myc (readthrough) in steady state extracts. 3-4 biological replicates of *dfm1*Δ were included in each of 3 experiments. Right, for each replicate, CPY-K12 signal intensity was normalized to Pgk1. The mean wild type value across 4 biological replicates was set to 1. Means were compared by one-way ANOVA with Tukey’s multiple comparison tests. *p* values are indicated above the graph. (C, E) Red circles, ubiquitin. (B, D, G) Pgk1, loading control.

Our prior work indicated Dfm1 is not required for Hrd1-mediated TQC, as degradation of the Hrd1-dependent substrate *Deg1*\*-Sec62 (Figure 2C) proceeds independently of Dfm1 (Rubenstein *et al*., 2012). Because *dfm1*Δ phenotypes are prone to genetic suppression, we repeated this analysis using three independently generated *dfm1*Δ strains (Figures 2D and S1A). Although genetic suppression remains a formal possibility, the failure of multiple *dfm1*Δ isolates to stabilize *Deg1*\*-Sec62 supports the conclusion that Dfm1 does not contribute to the Hrd1 TQC pathway.

We next tested whether Dfm1 contributes to Ltn1-mediated co-translational TQC (Figure 2E) using the model substrate CPY-K12 (Crowder *et al*., 2015). CPY-K12 consists of ER-targeted CPY followed by two Protein A tags, a translation-arresting polylysine (K12) tract, and 13 myc tags, which are translated only upon K12 readthrough (Figure 2F). Translationally arrested (CPY-K12) and readthrough (CPY-K12-13myc) products are distinguished by molecular weight. TQC defects are reflected by increased abundance of CPY-K12. Loss of *DFM1* did not significantly alter abundance of stalled CPY-K12 (Figure 2G). Together, these results suggest Dfm1 function in TQC is primarily limited to the Ste24 pathway.

### Dfm1 mitigates IAPP-induced translocon stress

To further investigate Dfm1 function in TQC, we assessed sensitivity to 6xIAPP, a yeast expression model of oligomeric islet amyloid polypeptide (IAPP) (Kayatekin *et al*., 2018) (Figure 3A). β cell failure in diabetes correlates with accumulation of oligomeric IAPP, which impedes translocon function (Lorenzo *et al*., 1994; Jurgens *et al*., 2011; Mukherjee *et al*., 2017; Kayatekin *et al*., 2018). Consistent with our Clogger results, loss of *DFM1* sensitized cells to galactose-induced 6xIAPP expression to a similar extent as *STE24* deletion, whereas *hrd1*Δ yeast showed minimal sensitivity (Figure 3B).

**Figure 3.**
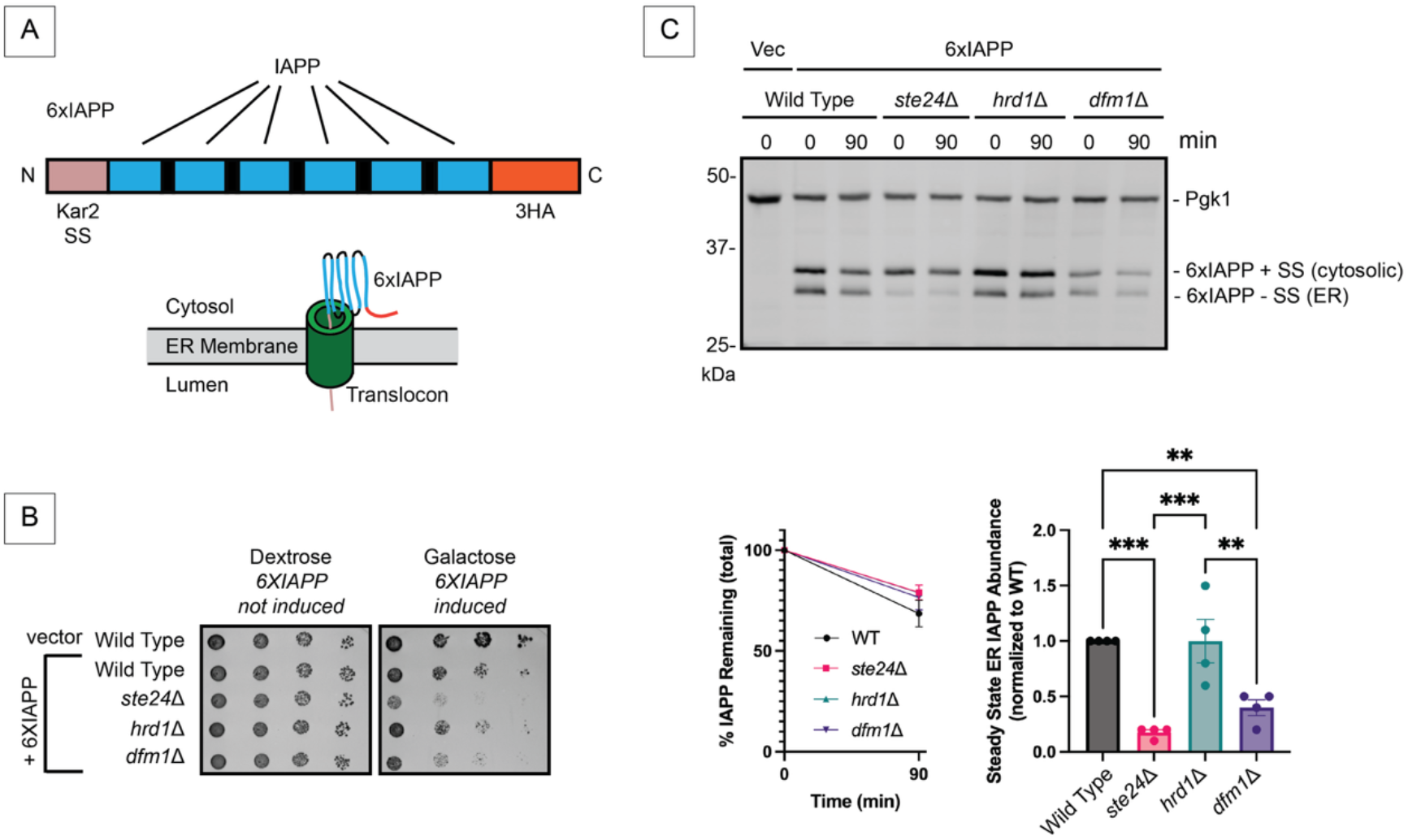
Dfm1 and Ste24 mitigate IAPP toxicity. (A) The yeast expression variant of 6xIAPP contains the Kar2 signal sequence (SS) to ensure efficient ER targeting, 6 copies of mature IAPP separated by short linkers, and 3 HA tags. (B) 6-fold dilutions of yeast transformed with a plasmid encoding 6xIAPP-3HA or an empty vector were spotted onto medium containing dextrose or galactose. (C) Top, cycloheximide chase of 6xIAPP-3HA. Bottom left, means of % total (cytosolic + ER) 6xIAPP-3HA remaining over time plotted from 3 biological replicates. Means of % 6xIAPP-3HA remaining at 90 min were compared by one-way ANOVA with Tukey’s multiple comparison test. No significant differences were observed. Bottom right, means of steady state abundance of ER (SS-cleaved) 6xIAPP (normalized to wild type) were compared by one-way ANOVA with Tukey’s multiple comparison test. **, *p* < 0.01. ***, *p* < 0.001. Pgk1, loading control.

We next examined 6xIAPP processing by cycloheximide chase in wild type, *hrd1*Δ, *ste24*Δ, and *dfm1*Δ cells (Figure 3C). Although 6xIAPP was highly stable in all cases, differences in maturation were consistent with impaired translocon clearance in *ste24*Δ and *dfm1*Δ cells. Both mutants showed reduced abundance of ER-resident 6xIAPP (i.e. signal sequence-cleaved protein), consistent with decreased ER translocation efficiency and unresolved translocon clogging. These results further support a model in which Dfm1 collaborates with Ste24 in response to translocon clogging.

### Increased gene dosage of wild type or inactive *HRD1* or *STE24* rescues *dfm1*Δ TQC phenotypes

The requirement for Dfm1 in retrotranslocation of misfolded transmembrane Hrd1 substrates can be bypassed by Hrd1 overproduction, which provides an alternate route for membrane extraction (Neal *et al*., 2018). We tested whether increased production of TQC factors Hrd1 or Ste24 rescues Clogger sensitivity of *dfm1*Δ yeast recovered from the original screen. Low-copy (CEN) plasmids encoding *HRD1* or *STE24* driven by their native promoters markedly improved growth of *dfm1*Δ cells under clogging stress (Figure 4A). Remarkably, catalytically inactive Hrd1 or Ste24 variants also restored Clogger resistance (Figure 4B). Rescue was observed in both the original screen-derived (Figures 4A and 4B) and newly generated *dfm1*Δ strains (Figure 4C).

**Figure 4.**
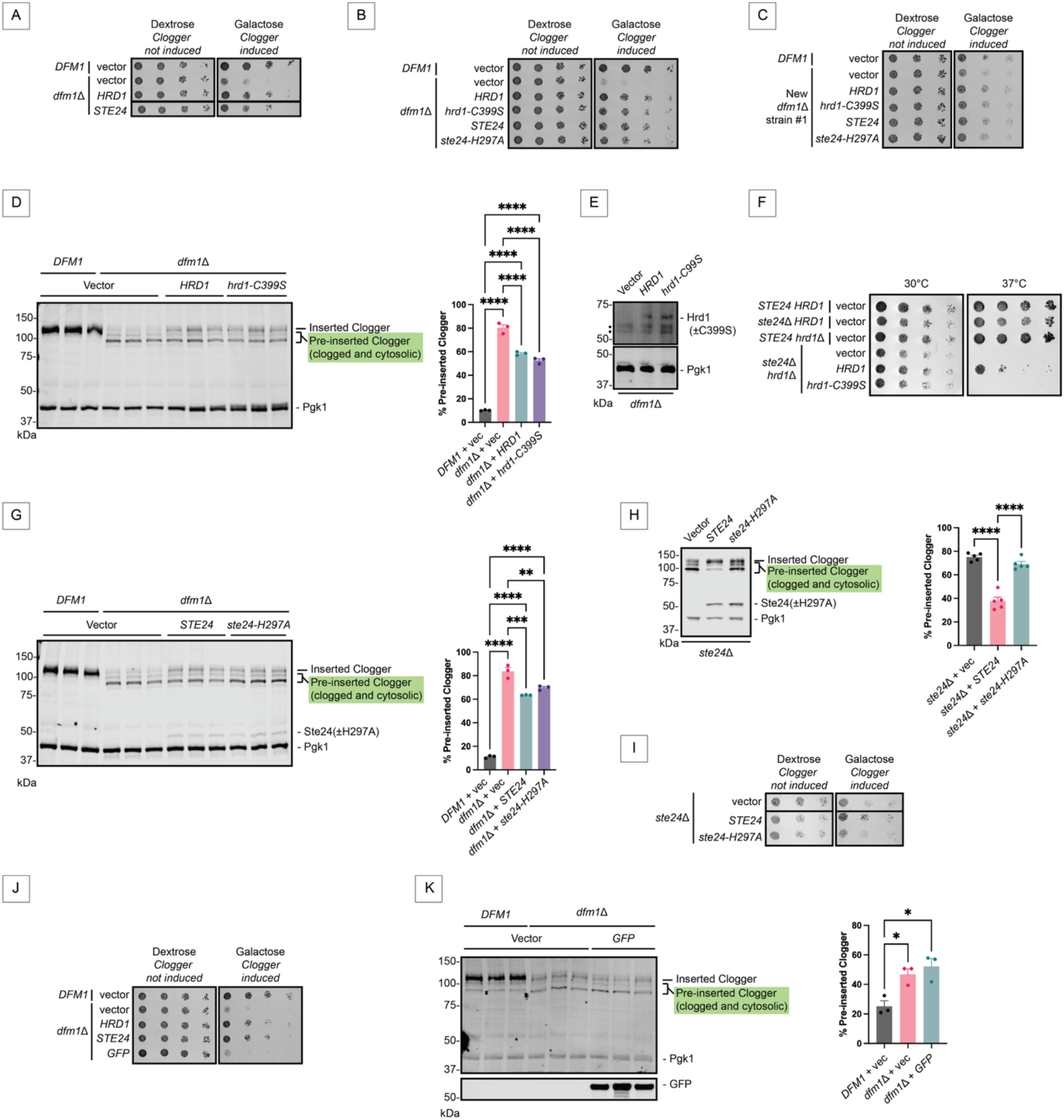
Increased *HRD1* or *STE24* dosage rescues translocon clogging sensitivity of *dfm1*Δ yeast. (A-C, F, I, J) 6-fold dilutions of yeast harboring indicated plasmids were spotted onto medium containing dextrose or galactose. (D, G, H, K) Left, western blot of Clogger in steady state extracts of yeast harboring indicated plasmid from 3 biological replicates. Ste24(-H297A), which possesses an N-terminal 3HA tag, is detected in (G) and (H), and GFP is detected in (K). Right, means of % Clogger that was pre-inserted from 3 biological replicates were compared by one-way ANOVA with Tukey’s multiple comparison test. *, *p* < 0.05. **, *p* < 0.01. ***, *p* < 0.001. ****, *p* < 0.0001. (E) Western blot of Hrd1(-C399S), which possesses a C-terminal 3HA tag, from steady state lysates prepared from representative parallel dextrose-grown cultures in (D). Pgk1, loading control.•, non-specific band.

To determine whether rescue reflected improved Clogger clearance, we examined Clogger dynamics in *dfm1*Δ cells expressing plasmid-borne wild type or inactive Hrd1 or Ste24. Surprisingly, increased *HRD1* or *STE24* dosage did not reduce the absolute abundance of pre-inserted Clogger. Instead, it consistently increased the proportion of fully inserted (i.e. luminal) Clogger (Figures 4D and 4G). Expression of plasmid-encoded Hrd1 and Ste24 was confirmed by immunoblotting (Figures 4E and 4G). This effect was specific, as plasmid-driven expression of GFP neither rescued the growth defect nor increased fully inserted Clogger (Figures 4J and 4K).

Catalytic inactivation was confirmed by plasmid sequencing (File S1) and phenotypic analysis. Wild type Hrd1, but not hrd1-C399S, partially rescued the synthetic growth defect of *ste24*Δ *hrd1*Δ cells (Figure 4F). Similarly, wild type Ste24, but not ste24-H297A, complemented TQC defects in *ste24*Δ cells (Figures 4H and 4I), indicating Ste24 catalytic activity is required under normal conditions.

These findings suggest that increased levels of Hrd1 or Ste24 promote resolution of clogged translocons independent of catalytic activity and without directly degrading the obstructing substrate, revealing a non-enzymatic function in TQC. We speculate that excess TQC factors act as sinks that sequester clogging proteins laterally released from translocons. Both Hrd1 and Ste24 are multipass membrane proteins capable of binding substrates within membrane-embedded cavities (Pryor *et al*., 2013; Neal *et al*., 2020; Wu *et al*., 2020; Spear *et al*., 2026), raising the possibility that increased abundance buffers toxic cloggers.

### Spontaneous phenotypic suppression of *dfm1*Δ yeast

While culturing the screen-derived *dfm1*Δ strain under Clogger-inducing conditions, we isolated fast-growing suppressor colonies (Figure 5A). Three independent suppressor isolates exhibited wild type resistance to clogging stress (Figure 5B). Restored fitness coincided with clearance of pre-inserted Clogger species (Figure 5C), indicating that suppression restores TQC through a mechanism distinct from Hrd1 or Ste24 overproduction. The emergence of suppressors under conditions of clogging stress echoes prior studies showing spontaneous suppression of *dfm1*Δ sensitivity to misfolded membrane proteins (Neal *et al*., 2018). To determine whether large-scale genomic changes accompanied suppression, we assessed the ploidy of the suppressor strains. Flow cytometry-based ploidy analysis of these suppressors and the parental *dfm1*Δ strain indicated that one suppressor had undergone diploidization (Figure S1C). The molecular basis of suppression and its relationship to TQC regulation will be examined in future studies.

**Figure 5.**
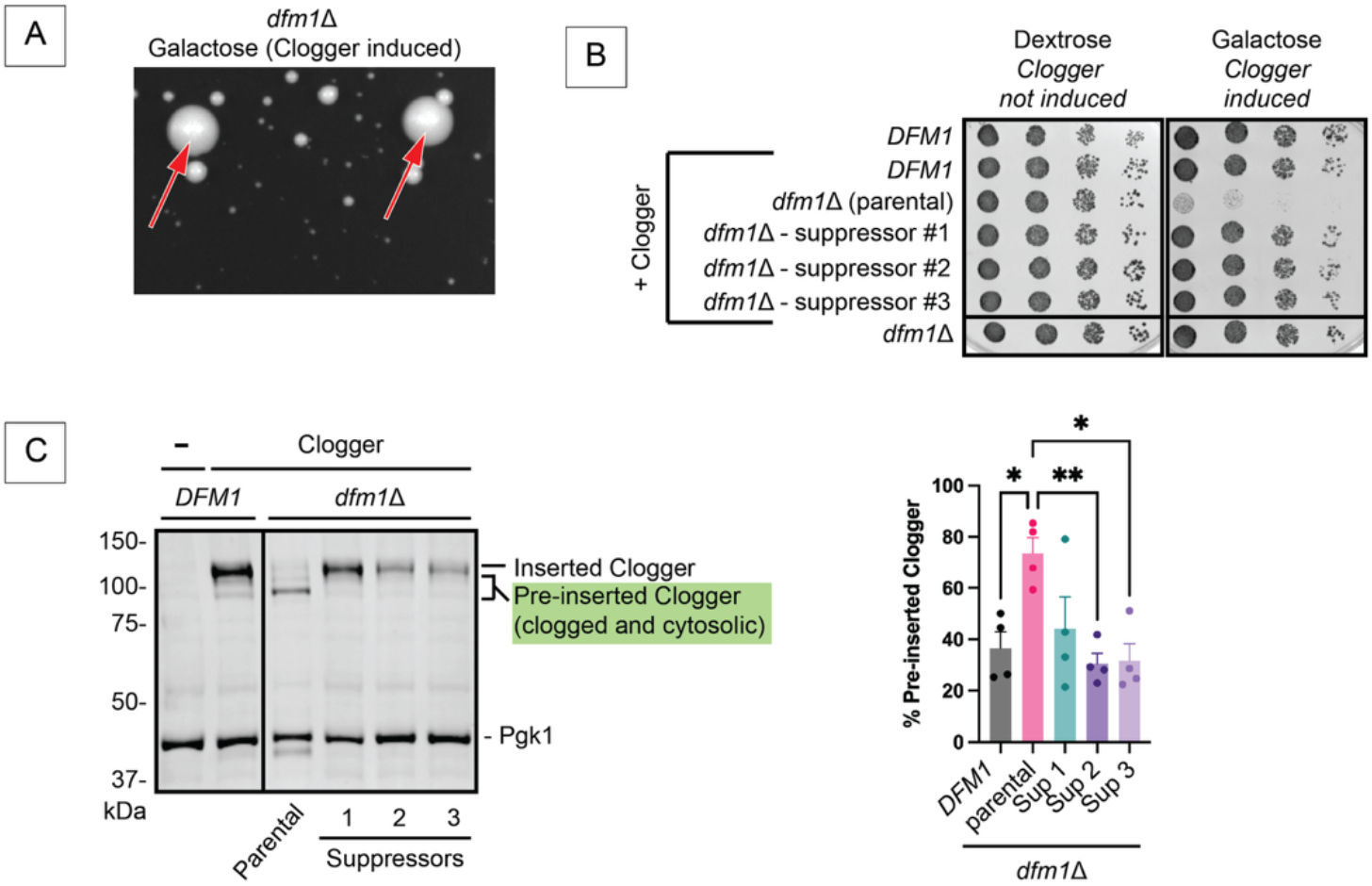
Spontaneous suppression of Clogger sensitivity of *dfm1*Δ yeast. (A) *dfm1*Δ yeast expressing Clogger were streaked onto medium containing galactose. Arrows indicate suppressors. (B) 6-fold dilutions of yeast were spotted onto medium containing dextrose or galactose. Experiment was repeated 3 times. (C) Left, western blot of Clogger in steady state extracts. Right, means of % Clogger that was pre-inserted from 4 biological replicates were compared by one-way ANOVA with Tukey’s multiple comparison test. *, *p* < 0.05. **, *p* < 0.01. Pgk1, loading control.

Overall, these findings define Dfm1 as a key component of the Ste24-dependent branch of TQC and support a model in which Dfm1 promotes extraction of translocon-clogging polypeptides from the Sec61 channel. We speculate that this process occurs via Cdc48-dependent retrotranslocation. This model is consistent with increased IAPP toxicity in *cdc48* mutants (Kayatekin *et al*., 2018) and recruitment of Cdc48 to clogged translocons (Ast *et al*., 2016). Increased Hrd1 or Ste24 levels mitigated toxicity associated with translocon clogging despite retaining clogged substrates, suggesting that increasing the buffering capacity of TQC factors can preserve ER function independently of substrate degradation. Increasing the abundance or buffering capacity of their homologs may represent a strategy to preserve ER function and pancreatic β-cell viability during IAPP-induced translocation stress associated with diabetes.

## Materials and Methods

### Yeast and Plasmid Methods

Unless otherwise indicated, yeast were maintained at 30°C and cultured on rich yeast extract-peptone-dextrose (YPD) or minimal synthetic defined (SD) media (Guthrie and Fink, 2004), which have 2% dextrose as the carbon source. For induction of genes under the control of the *GAL1* promoter (Clogger and 6xIAPP), dextrose was replaced with 2% galactose. Lithium acetate transformation was used to introduce plasmids into yeast. Yeast strains and plasmids are presented in Tables S1 and S2.

To delete *DFM1*, a cassette containing *HIS3MX6* flanked by 49 bp of DNA homologous to sequence upstream and downstream of *DFM1* was PCR-amplified from pFA6a-His3MX6 (Wach *et al*., 1997) using primers VJR515 (AGTGAATTCCGCATTTCCAAGTCAAATCAAAAACTATTTTCGAGGAAATACGGATCCCCGGGTTAATTAA) and VJR516 (TCTTGATGTATATAATGAATGGCAAAGTACATAGAAATAGATAAAAGTTGGAATTCGAGCTCGTTTAAAC). This cassette was directed to the yeast *DFM1* locus by homologous recombination. Integration at the *DFM1* locus was confirmed by PCR at the 5’ and 3’ integration junctions. pRS316-Hrd1-3HA (pVJ476) and pRS316-hrd1-C399S-3HA (pVJ498) were generated by inserting EcoRI/SacI fragments containing the *HRD1* promoter and *HRD1-3HA* or *hrd1-C399S-3HA* from pRH642 (Sato *et al*., 2009) and pRH1245 (Gardner *et al*., 2000), respectively, into EcoRI and SacI sites of pRS316. p416-*P*_*GAL1*_-Kar2SS-6xIAPP-3HA (pVJ636) was generated by custom plasmid synthesis (GenScript). p316-DFM1-3HA (pVJ691) was generated by inserting a SacI/XhoI fragment containing the *DFM1* promoter, *DFM1*, and 3HA tag from pSN113 (equivalent to pSN90 in (Nejatfard *et al*., 2021)) into the SacI and XhoI sites of pRS316. To generate p316-DFM1 (pVJ704), a SpeI/XhoI fragment containing a portion of *DFM1* without the 3HA tag from pSN164 (Bhaduri *et al*., 2023a) was inserted into SpeI and XhoI sites of pVJ691. All plasmids used or created for this study have been confirmed by whole plasmid sequencing by Plasmidsaurus using Oxford Nanopore Technology (File S1).

### Cycloheximide Chase, Cell Lysis, Western Blotting, and Data Analysis

Cycloheximide chase experiments were performed as described (Buchanan *et al*., 2016). In brief, yeast were cultured to exponential growth prior to treatment with 250 µg/ml cycloheximide and collection of cells at the indicated times. Proteins were extracted by alkaline lysis (Kushnirov, 2000; Watts *et al*., 2015). Following SDS-PAGE, proteins were transferred to polyvinylidene difluoride membranes by wet transfer at 20 V for 1 h at 4°C. Membranes were blocked in 5% (w/v) skim milk in tris-buffered saline (TBS) for 1 h at room temperature or overnight at 4°C. Clogger, Hrd1-3HA variants, 3HA-Ste24 variants, and 6xIAPP-3HA were detected with mouse anti-HA.11 antibodies (clone 16B12; BioLegend) at a dilution of 1:1,000 to 1:10,000. GFP was detected with mouse anti-GFP antibodies (clone JL-8; Clontech) at a dilution of 1:1,000. Pdi1 was detected with mouse anti-PDI (clone 38H8; Invitrogen). Pgk1 was detected with mouse anti-Pgk1 (clone 22C5D8; Life Technologies) at a dilution of 1:20,000 to 1:40,000. Mouse primary antibodies were followed by AlexaFluor-680-conjugated rabbit anti-mouse secondary antibodies at a dilution of 1:20,000 to 1:40,000. *Deg1*\*-Sec62 and CPY variants possess tandem *Staphylococcus aureus* Protein A tags (Hjelm *et al*., 1972), which were detected directly by AlexaFluor-680-conjugated rabbit anti-mouse secondary antibodies at a dilution of 1:20,000 to 1:40,000. Membranes were imaged using Odyssey CLx and DLx IR imaging systems (LI-COR). Relative protein abundance was determined using ImageStudio software (LI-COR). Band intensity of proteins of interest were subjected to background correction followed by normalization to background-adjusted Pgk1 (loading control). Data were analyzed by one-way ANOVA with Tukey’s multiple comparison test using GraphPad Prism 10. Error bars represent standard error of mean. All observations were independent of each other.

### DNA Content Profiling

For DNA content profiling, yeast were grown to saturation overnight. In the morning, cultures were diluted to OD_600_ = 0.1 and incubated with shaking at 30°C until OD_600_ = 0.6. Then, four OD_600_ equivalents were harvested by centrifugation at 10,000 x *g*, room temperature, washed once with 1 mL TE (10 mM Tris-HCl, 1 mM EDTA, pH 8.0), and centrifuged again, and the supernatant was aspirated. The cell pellet was then resuspended in 3 mL of TE, and added to 7 mL of 70% ethanol prepared in TE dropwise while vortexing to minimize aggregation. The sample was nutated overnight at 4°C. The following morning, the cells were centrifuged at 5000 x *g*, room temperature, and the supernatant was removed. The pellet was then washed three times with 1 mL of TE buffer to remove residual ethanol and then resuspended in 1 mL TE with 0.25 µg/mL RNase (Qiagen) for 2 hours at 37°C to remove RNA. The sample was then centrifuged and washed with 1 mL of TE as above and resuspended in 1 mL TE with 0.25 µg/mL proteinase K before incubation at 50°C for one hour. Following this incubation, the cells were pelleted as above, washed with 1 mL TE, and then stained overnight at 4°C with 1 mL of 10 µg/mL propidium iodide prepared in TE. The samples were then analyzed on a Cytek Aurora flow cytometer with forward and side scatter gating to exclude debris and cell aggregates. Known haploid and diploid control strains were used to calibrate the 1n, 2n, and 4n DNA peaks. For each sample, at least 20,000 PI-positive cells were included in analysis.

## Abbreviations

ER: endoplasmic reticulum
IAPP: islet amyloid polypeptide
PQC: protein quality control
TQC: translocon quality control

## Acknowledgments

We thank Randy Hampton, Toshifumi Inada, Can Kayatekin, Stefan Kreft, Susan Michaelis, Sonya Neal, and Maya Schuldiner for generously sharing plasmids, strains, and advice. We thank Lindy Caffo and Kieran Claypool for assistance with generating or validating reagents. We thank Douglas Bernstein, Philip Smaldino, Douglas Roossien, Ashley Kalinski, Paul Venturelli, and members of the Rubenstein lab for helpful discussions. We thank the *Saccharomyces* Genome Database for curation of yeast genetic information (Wong *et al*., 2023).

## References

Arakawa, S., Yunoki, K., Izawa, T., Tamura, Y., Nishikawa, S., and Endo, T. (2016). Quality control of nonstop membrane proteins at the ER membrane and in the cytosol. Sci Rep 6, 30795.

Ast, T., Michaelis, S., and Schuldiner, M. (2016). The Protease Ste24 Clears Clogged Translocons. Cell 164, 103–114.

Bhaduri, S., Aguayo, A., Ohno, Y., Proietto, M., Jung, J., Wang, I., Kandel, R., Singh, N., Ibrahim, I., Fulzele, A., Bennett, E.J., Kihara, A., and Neal, S.E. (2023a). An ERAD-independent role for rhomboid pseudoprotease Dfm1 in mediating sphingolipid homeostasis. EMBO J 42, e112275.

Bhaduri, S., Scott, N.A., and Neal, S.E. (2023b). The Role of the Rhomboid Superfamily in ER Protein Quality Control: From Mechanisms and Functions to Diseases. Cold Spring Harb Perspect Biol 15, a041248.

Buchanan, B.W., Lloyd, M.E., Engle, S.M., and Rubenstein, E.M. (2016). Cycloheximide Chase Analysis of Protein Degradation in Saccharomyces cerevisiae. Journal of visualized experiments : JoVE 110, 53975.

Carvalho, P., Goder, V., and Rapoport, T.A. (2006). Distinct ubiquitin-ligase complexes define convergent pathways for the degradation of ER proteins. Cell 126, 361–373.

Chen, Y., Zhang, Y., Yin, Y., Gao, G., Li, S., Jiang, Y., Gu, X., and Luo, J. (2005). SPD--a web-based secreted protein database. Nucleic Acids Res 33, D169–173.

Choi, J., Park, J., Kim, D., Jung, K., Kang, S., and Lee, Y.H. (2010). Fungal secretome database: integrated platform for annotation of fungal secretomes. BMC Genomics 11, 105.

Crowder, J.J., Geigges, M., Gibson, R.T., Fults, E.S., Buchanan, B.W., Sachs, N., Schink, A., Kreft, S.G., and Rubenstein, E.M. (2015). Rkr1/Ltn1 Ubiquitin Ligase-Mediated Degradation of Translationally Stalled Endoplasmic Reticulum Proteins. J Biol Chem 290, 18454–18466.

Ennis, A., Wang, L., Xu, Y., Saidi, L., Wang, X., Yu, C., Yun, S., Huang, L., and Ye, Y. (2025). NEMF-mediated CAT tailing facilitates translocation-associated quality control. J Cell Biol 224.

Gardner, R.G., Swarbrick, G.M., Bays, N.W., Cronin, S.R., Wilhovsky, S., Seelig, L., Kim, C., and Hampton, R.Y. (2000). Endoplasmic reticulum degradation requires lumen to cytosol signaling. Transmembrane control of Hrd1p by Hrd3p. J Cell Biol 151, 69–82.

Goder, V., Carvalho, P., and Rapoport, T.A. (2008). The ER-associated degradation component Der1p and its homolog Dfm1p are contained in complexes with distinct cofactors of the ATPase Cdc48p. FEBS Lett 582, 1575–1580.

Guthrie, C., and Fink, G.R. (2004). Guide to Yeast Genetics and Molecular and Cell Biology. Elsevier: San Diego.

Hjelm, H., Hjelm, K., and Sjoquist, J. (1972). Protein A from Staphylococcus aureus. Its isolation by affinity chromatography and its use as an immunosorbent for isolation of immunoglobulins. FEBS Lett 28, 73–76.

Horn, S.C., Hanna, J., Hirsch, C., Volkwein, C., Schutz, A., Heinemann, U., Sommer, T., and Jarosch, E. (2009). Usa1 functions as a scaffold of the HRD-ubiquitin ligase. Mol Cell 36, 782–793.

Itskanov, S., and Park, E. (2023). Mechanism of Protein Translocation by the Sec61 Translocon Complex. Cold Spring Harb Perspect Biol 15, a041250.

Jurgens, C.A., Toukatly, M.N., Fligner, C.L., Udayasankar, J., Subramanian, S.L., Zraika, S., Aston-Mourney, K., Carr, D.B., Westermark, P., Westermark, G.T., Kahn, S.E., and Hull, R.L. (2011). beta-cell loss and beta-cell apoptosis in human type 2 diabetes are related to islet amyloid deposition. Am J Pathol 178, 2632–2640.

Kandel, R.R., and Neal, S.E. (2020). The role of rhomboid superfamily members in protein homeostasis: Mechanistic insight and physiological implications. Biochim Biophys Acta Mol Cell Res 1867, 118793.

Kayatekin, C., Amasino, A., Gaglia, G., Flannick, J., Bonner, J.M., Fanning, S., Narayan, P., Barrasa, M.I., Pincus, D., Landgraf, D., Nelson, J., Hesse, W.R., Costanzo, M., Consortium, A.T.D.G., Myers, C.L., Boone, C., Florez, J.C., and Lindquist, S. (2018). Translocon Declogger Ste24 Protects against IAPP Oligomer-Induced Proteotoxicity. Cell 173, 62–73 e69.

Knop, M., Finger, A., Braun, T., Hellmuth, K., and Wolf, D.H. (1996). Der1, a novel protein specifically required for endoplasmic reticulum degradation in yeast. EMBO J 15, 753–763.

Kushnirov, V.V. (2000). Rapid and reliable protein extraction from yeast. Yeast 16, 857–860.

Lemberg, M.K. (2013). Sampling the membrane: function of rhomboid-family proteins. Trends Cell Biol 23, 210–217.

Lim, J.J., Lee, Y., Yoon, S.Y., Ly, T.T., Kang, J.Y., Youn, H.S., An, J.Y., Lee, J.G., Park, K.R., Kim, T.G., Yang, J.K., Jun, Y., and Eom, S.H. (2016). Structural insights into the interaction of human p97 N-terminal domain and SHP motif in Derlin-1 rhomboid pseudoprotease. FEBS Lett 590, 4402–4413.

Lorenzo, A., Razzaboni, B., Weir, G.C., and Yankner, B.A. (1994). Pancreatic islet cell toxicity of amylin associated with type-2 diabetes mellitus. Nature 368, 756–760.

Mukherjee, A., Morales-Scheihing, D., Salvadores, N., Moreno-Gonzalez, I., Gonzalez, C., Taylor-Presse, K., Mendez, N., Shahnawaz, M., Gaber, A.O., Sabek, O.M., Fraga, D.W., and Soto, C. (2017). Induction of IAPP amyloid deposition and associated diabetic abnormalities by a prion-like mechanism. J Exp Med 214, 2591–2610.

Neal, S., Jaeger, P.A., Duttke, S.H., Benner, C., c, K.G., Ideker, T., and Hampton, R.Y. (2018). The Dfm1 Derlin Is Required for ERAD Retrotranslocation of Integral Membrane Proteins. Mol Cell 69, 306–320 e304.

Neal, S., Syau, D., Nejatfard, A., Nadeau, S., and Hampton, R.Y. (2020). HRD Complex Self-Remodeling Enables a Novel Route of Membrane Protein Retrotranslocation. iScience 23, 101493.

Nejatfard, A., Wauer, N., Bhaduri, S., Conn, A., Gourkanti, S., Singh, N., Kuo, T., Kandel, R., Amaro, R.E., and Neal, S.E. (2021). Derlin rhomboid pseudoproteases employ substrate engagement and lipid distortion to enable the retrotranslocation of ERAD membrane substrates. Cell Rep 37, 109840.

Pryor, E.E., Jr., Horanyi, P.S., Clark, K.M., Fedoriw, N., Connelly, S.M., Koszelak-Rosenblum, M., Zhu, G., Malkowski, M.G., Wiener, M.C., and Dumont, M.E. (2013). Structure of the integral membrane protein CAAX protease Ste24p. Science 339, 1600–1604.

Rubenstein, E.M., Kreft, S.G., Greenblatt, W., Swanson, R., and Hochstrasser, M. (2012). Aberrant substrate engagement of the ER translocon triggers degradation by the Hrd1 ubiquitin ligase. J Cell Biol 197, 761–773.

Runnebohm, A.M., Richards, K.A., Irelan, C.B., Turk, S.M., Vitali, H.E., Indovina, C.J., and Rubenstein, E.M. (2020). Overlapping function of Hrd1 and Ste24 in translocon quality control provides robust channel surveillance. J Biol Chem 295, 16113–16120.

Sato, B.K., and Hampton, R.Y. (2006). Yeast Derlin Dfm1 interacts with Cdc48 and functions in ER homeostasis. Yeast 23, 1053–1064.

Sato, B.K., Schulz, D., Do, P.H., and Hampton, R.Y. (2009). Misfolded membrane proteins are specifically recognized by the transmembrane domain of the Hrd1p ubiquitin ligase. Mol Cell 34, 212–222.

Scavone, F., Gumbin, S.C., Da Rosa, P.A., and Kopito, R.R. (2023). RPL26/uL24 UFMylation is essential for ribosome-associated quality control at the endoplasmic reticulum. Proc Natl Acad Sci U S A 120, e2220340120.

Scott, N.A., Afolabi, J.M., Marshall, A.G., Schafer, J.C., Baskerville, V.R., Prasad, P., Kadam, A.A., Som de Cerff, C., Whisenant, T., Phillips, M.A., Tomar, D., McReynolds, M.R., Hinton Jr., A., and Neal, S.E. (2026). Derlin-mediated ERAD of lipid regulator ORMDL3 safeguards mitochondrial function. bioRxiv, 10.64898/62026.64802.64827.708653

Spear, E.D., Shilagardi, K., Sarju, S., and Michaelis, S. (2026). The zinc metalloprotease ZMPSTE24 binds a distinct topological isoform of the tail-anchored protein IFITM3. bioRxiv, 10.64898/62026.64802.64827.708584.

Stolz, A., Schweizer, R.S., Schafer, A., and Wolf, D.H. (2010). Dfm1 forms distinct complexes with Cdc48 and the ER ubiquitin ligases and is required for ERAD. Traffic 11, 1363–1369.

Vitali, D.G., Fonseca, D., and Carvalho, P. (2024). The derlin Dfm1 couples retrotranslocation of a folded protein domain to its proteasomal degradation. J Cell Biol 223, e202308074.

von der Malsburg, K., Shao, S., and Hegde, R.S. (2015). The Ribosome Quality Control Pathway can Access Nascent Polypeptides Stalled at the Sec61 Translocon. Mol Biol Cell 26, 2168–2180.

Wach, A., Brachat, A., Alberti-Segui, C., Rebischung, C., and Philippsen, P. (1997). Heterologous HIS3 marker and GFP reporter modules for PCR-targeting in Saccharomyces cerevisiae. Yeast 13, 1065–1075.

Wang, L., Xu, Y., Rogers, H., Saidi, L., Noguchi, C.T., Li, H., Yewdell, J.W., Guydosh, N.R., and Ye, Y. (2020). UFMylation of RPL26 links translocation-associated quality control to endoplasmic reticulum protein homeostasis. Cell Res 30, 5–20.

Wang, L., and Ye, Y. (2020). Clearing Traffic Jams During Protein Translocation Across Membranes. Front Cell Dev Biol 8, 610689.

Watts, S.G., Crowder, J.J., Coffey, S.Z., and Rubenstein, E.M. (2015). Growth-based Determination and Biochemical Confirmation of Genetic Requirements for Protein Degradation in Saccharomyces cerevisiae. Journal of visualized experiments : JoVE 96, e52428.

Wong, E.D., Miyasato, S.R., Aleksander, S., Karra, K., Nash, R.S., Skrzypek, M.S., Weng, S., Engel, S.R., and Cherry, J.M. (2023). Saccharomyces genome database update: server architecture, pan-genome nomenclature, and external resources. Genetics 224.

Wu, X., Siggel, M., Ovchinnikov, S., Mi, W., Svetlov, V., Nudler, E., Liao, M., Hummer, G., and Rapoport, T.A. (2020). Structural basis of ER-associated protein degradation mediated by the Hrd1 ubiquitin ligase complex. Science 368, eaaz2449.

