## Supplemental Materials for "Dfm1 promotes Ste24-dependent translocon quality control"

**Running Head: Dfm1 promotes translocon quality control**

### Supplemental Tables

**Table S1. Yeast strains used in this study.**

| Name | Genotype | Source |
|---|---|---|
| VJY246 | *MATa his3Δ1 leu2Δ0 met15Δ0 ura3Δ0 ltn1Δ::kanMX4* | (Tong *et al.*, 2001) |
| VJY378 | *MATa his3Δ1 leu2Δ0 met15Δ0 ura3Δ0 ste24Δ::kanMX4* | (Tong *et al.*, 2001) |
| VJY405 (2379/29) | *MATa his3Δ1 leu2Δ0 ura3Δ0 met15Δ0 HO::NatR-Galp-PDIClogger ste24Δ::pcgURA* | (Ast *et al.*, 2016) |
| VJY457 | *MATa his3Δ1 leu2Δ0 met15Δ0 ura3Δ0 hrd1Δ::kanMX4 ste24Δ::natMX4* | (Runnebohm *et al.*, 2020) |
| VJY476 (BY4741) | *MATa his3Δ1 leu2Δ0 met15Δ0 ura3Δ0* | (Tong *et al.*, 2001) |
| VJY511 | *MATa his3Δ1 leu2Δ0 met15Δ0 ura3Δ0 hrd1Δ::kanMX4* | (Tong *et al.*, 2001) |
| VJY512 | *MATa his3Δ1 leu2Δ0 lys2Δ0 ura3Δ0 met15Δ0 HO::NatR-Galp-PDIClogger* | (Runnebohm *et al.*, 2020) |
| VJY513 | *MATa his3Δ1 leu2Δ0 lys2Δ0 ura3Δ0 met15Δ0 HO::NatR-Galp-PDIClogger ste24Δ::pcgURA hrd1Δ::kanMX4* | (Runnebohm *et al.*, 2020) |
| VJY521 | *MATa his3Δ1 leu2Δ0 ura3Δ0 met15Δ0 HO::NatR-Galp-PDIClogger hrd1Δ::kanMX4* | (Runnebohm *et al.*, 2020) |
| VJY931 (SKY527) | *MATa his3Δ1 leu2Δ0 met15Δ0 ura3Δ0 dfm1Δ::hphMX4* | Gift of Stefan Kreft |
| VJY935 (YMS2384; "Original *dfm1Δ*") | *MATα his3Δ1 leu2Δ0 met15Δ0 ura3Δ0 lyp1Δ::STE3pr-LEU2 can1Δ::STE2pr-spHIS5 HO::NatR-Galp-PDIClogger dfm1Δ::KanR* | (Ast *et al.*, 2016) |
| VJY952 ("*dfm1Δ* Supp 1") | *MATα his3Δ1 leu2Δ0 met15Δ0 ura3Δ0 lyp1Δ::STE3pr-LEU2 can1Δ::STE2pr-spHIS5 HO::NatR-Galp-PDIClogger dfm1Δ::KanR* | This study |
| VJY953 ("*dfm1Δ* Supp 2") | *MATα his3Δ1 leu2Δ0 met15Δ0 ura3Δ0 lyp1Δ::STE3pr-LEU2 can1Δ::STE2pr-spHIS5 HO::NatR-Galp-PDIClogger dfm1Δ::KanR* | This study |
| VJY954 ("*dfm1Δ* Supp 3") | *MATα his3Δ1 leu2Δ0 met15Δ0 ura3Δ0 lyp1Δ::STE3pr-LEU2 can1Δ::STE2pr-spHIS5 HO::NatR-Galp-PDIClogger dfm1Δ::KanR* | This study |
| VJY1077 | *MATa his3Δ1 leu2Δ0 met15Δ0 ura3Δ0 dfm1Δ::kanMX4* | (Tong *et al.*, 2001) |
| VJY1129 | *MATa his3Δ1 leu2Δ0 met15Δ0 ura3Δ0 HO::NatR-Galp-PDIClogger ste24Δ::pcgURA dfm1Δ::HIS3MX6* | This study |
| VJY1206 ("new *dfm1Δ* #1") | *MATa his3Δ1leu2Δ0 ura3Δ0 lys2Δ0 HO::NatR-Galp-PDIClogger met15Δ0 dfm1Δ::HIS3MX6* | This study |
| VJY1207 ("new *dfm1Δ* #2") | *MATa his3Δ1leu2Δ0 ura3Δ0 lys2Δ0 HO::NatR-Galp-PDIClogger met15Δ0 dfm1Δ::HIS3MX6* | This study |
| VJY1208 ("new *dfm1Δ* #3") | *MATa his3Δ1leu2Δ0 ura3Δ0 lys2Δ0 HO::NatR-Galp-PDIClogger met15Δ0 dfm1Δ::HIS3MX6* | This study |
| VJY1209 ("new *dfm1Δ* #4") | *MATa his3Δ1leu2Δ0 ura3Δ0 lys2Δ0 HO::NatR-Galp-PDIClogger met15Δ0 dfm1Δ::HIS3MX6* | This study |

**Table S2. Plasmids used in this study.**

| Name | Alias | Yeast selection | Description | Source |
|---|---|---|---|---|
| pRS313 | pVJ26 | *HIS3* | Empty vector | (Sikorski and Hieter, 1989) |
| pRS316 | pVJ27 | *URA3* | Empty vector | (Sikorski and Hieter, 1989) |
| p416-$P_{MET25}$-*Deg1**-Sec62-2xProtA | pVJ317 | *URA3* | *Deg1**-Flag-Sec62-2xProtA ("*Deg1**-Sec62") driven by *MET25* promoter; *Deg1** = F18S, I22T | (Rubenstein *et al.*, 2012) |
| YCPlac33-$P_{GPD}$-GFP-2A-Flag-HIS3 | pVJ382; pSA158 | *URA3* | GFP-2A-Flag-HIS3 in YCPlac33 backbone driven by *TDH3* (*GPD*) promoter | (Ito-Harashima *et al.*, 2007) |
| p413-$P_{GPD}$-CPY-ProtA | pVJ464; STK07.4.9 | *HIS3* | CPY-2xProtA driven by *TDH3* (*GPD*) promoter | (Crowder *et al.*, 2015) |
| pRS316-Hrd1-3HA | pVJ476 | *URA3* | Hrd1-3HA driven by native (*HRD1*) promoter | This study |
| p413-$P_{GPD}$-CPY-ProtA-K12-13myc | pVJ484; STK07.6.9 | *HIS3* | CPY-2xProtA-K12-13myc driven by *TDH3* (*GPD*) promoter; K12 = 12 lysine residues | (Crowder *et al.*, 2015) |
| pRS316-hrd1-C399S-3HA | pVJ498 | *URA3* | hrd1-C399S-3HA driven by native (*HRD1*) promoter; C399S inactivates catalytic RING domain | This study |
| p3HA-Ste24 | pVJ556; pSM1107 | *URA3* | 3HA-Ste24 driven by native (*STE24*) promoter | (Fujimura-Kamada *et al.*, 1997) |
| p3HA-ste24-H297A | pVJ557; pSM1103 | *URA3* | 3HA-ste24 -H297A driven by native (*STE24*) promoter; H297A is protease inactivating mutant | (Fujimura-Kamada *et al.*, 1997) |
| p416-$P_{GAL1}$-Kar2SS-6xIAPP-3HA | pVJ636 | *URA3* | Kar2SS-6xIAPP-3HA driven by *GAL1* promoter (Kar2SS = signal sequence for yeast Kar2) | This study |
| p316-Dfm1-3HA | pVJ691 | *URA3* | Dfm1-3HA driven by native (*DFM1*) promoter | This study |
| p316-Dfm1 | pVJ704 | *URA3* | Dfm1 driven by native (*DFM1*) promoter | This study |

All plasmids harbor the *AmpR* (ampicillin resistance gene) for bacterial selection. All constructs are yeast centromeric (CEN) plasmids.

**Supplemental Figure**

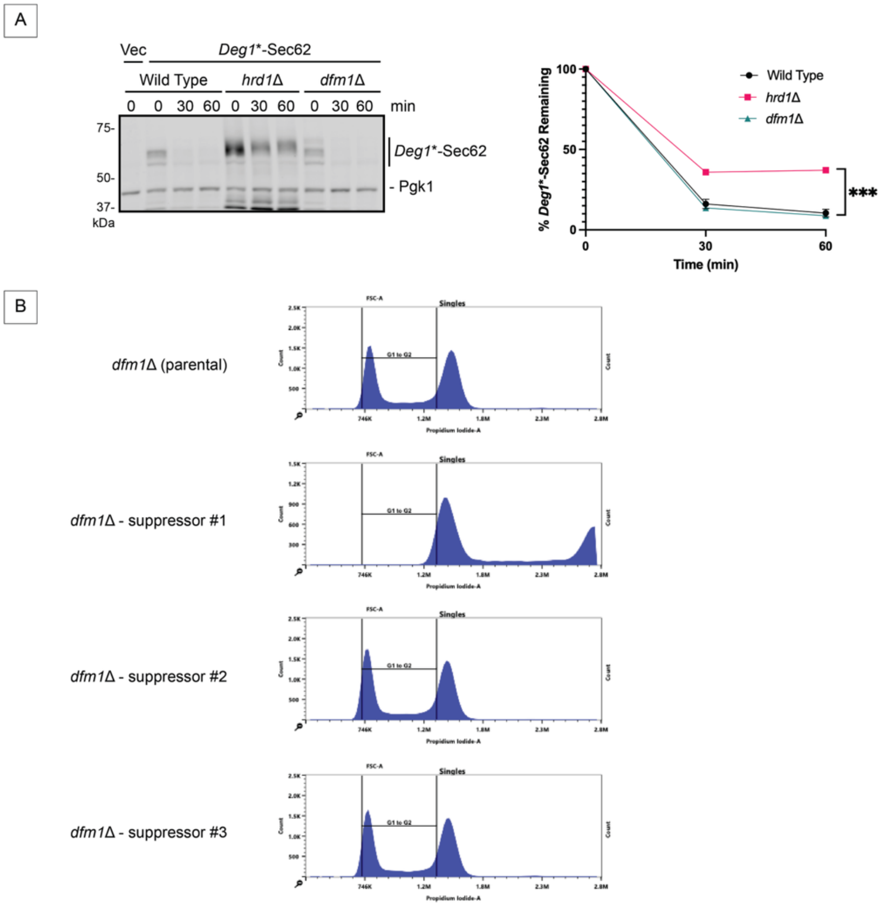


**Figure S1. Supporting Data.** (A) **Dfm1 is not required for Hrd1-dependent TQC.** Left, cycloheximide chase of *Deg1**-Sec62. The *dfm1*Δ strain used for these experiments is the newly generated *dfm1*Δ strain #1 with impaired Clogger clearance (see Figure 1E-1G), and distinct from the two *dfm1*Δ strains in Figure 2D. Right, means of % *Deg1**-Sec62 remaining over time plotted for 3 biological replicates. Means of % *Deg1**-Sec62 remaining at 60 min were compared by one-way ANOVA with Tukey's multiple comparison test. ***, $p < 0.001$ and applies to comparison between *hrd1*Δ yeast and both other strains. (B) **Flow cytometry-based ploidy analysis of parental and suppressor *dfm1*Δ yeast strains.** Propidium iodide staining was used to assess DNA content of the original screen-derived *dfm1*Δ parental strain and three independent suppressor strains. Histograms of single cells show DNA content distributions corresponding to G1 and G2 populations. Suppressor #1 displays a right-shifted DNA content profile compared with the parental strain and suppressors #2 and #3, consistent with altered ploidy.

**Supplemental References**


Ast, T., Michaelis, S., and Schuldiner, M. (2016). The Protease Ste24 Clears Clogged Translocons. Cell *164*, 103–114.

Crowder, J.J., Geigges, M., Gibson, R.T., Fults, E.S., Buchanan, B.W., Sachs, N., Schink, A., Kreft, S.G., and Rubenstein, E.M. (2015). Rkr1/Ltn1 Ubiquitin Ligase-Mediated Degradation of Translationally Stalled Endoplasmic Reticulum Proteins. J Biol Chem *290*, 18454–18466.

Fujimura-Kamada, K., Nouvet, F.J., and Michaelis, S. (1997). A novel membrane-associated metalloprotease, Ste24p, is required for the first step of NH2-terminal processing of the yeast a-factor precursor. J Cell Biol *136*, 271–285.

Ito-Harashima, S., Kuroha, K., Tatematsu, T., and Inada, T. (2007). Translation of the poly(A) tail plays crucial roles in nonstop mRNA surveillance via translation repression and protein destabilization by proteasome in yeast. Genes Dev *21*, 519–524.

Rubenstein, E.M., Kreft, S.G., Greenblatt, W., Swanson, R., and Hochstrasser, M. (2012). Aberrant substrate engagement of the ER translocon triggers degradation by the Hrd1 ubiquitin ligase. J Cell Biol *197*, 761–773.

Runnebohm, A.M., Richards, K.A., Irelan, C.B., Turk, S.M., Vitali, H.E., Indovina, C.J., and Rubenstein, E.M. (2020). Overlapping function of Hrd1 and Ste24 in translocon quality control provides robust channel surveillance. J Biol Chem *295*, 16113–16120.

Sikorski, R.S., and Hieter, P. (1989). A system of shuttle vectors and yeast host strains designed for efficient manipulation of DNA in Saccharomyces cerevisiae. Genetics *122*, 19–27.

Tong, A.H., Evangelista, M., Parsons, A.B., Xu, H., Bader, G.D., Page, N., Robinson, M., Raghibizadeh, S., Hogue, C.W., Bussey, H., Andrews, B., Tyers, M., and Boone, C. (2001). Systematic genetic analysis with ordered arrays of yeast deletion mutants. Science *294*, 2364–2368.

**Supplemental File S1**

**Plasmid sequences used in this study**

**pVJ26 (pRS313)**

ttgagatcctttttttctgcgcgtaatctgctgcttgcaaacaaaaaaaccaccgctaccagcg
gtggtttgtttgccggatcaagagctaccaactctttttccgaaggtaactggcttcagcagag
cgcagataccaaatactgttcttctagtgtagccgtagttaggccaccacttcaagaactctgt
agcaccgcctacatacctcgctctgctaatcctgttaccagtggctgctgccagtggcgataag
tcgtgtcttaccgggttggactcaagacgatagttaccggataaggcgcagcggtcgggctgaa
cggggggttcgtgcacacagcccagcttggagcgaacgacctacaccgaactgagatacctaca
gcgtgagctatgagaaagcgccacgcttcccgaagggagaaaggcggacaggtatccggtaagc
ggcagggtcggaacaggagagcgcacgagggagcttccagggggaaacgcctggtatctttata
gtcctgtcgggtttcgccacctctgacttgagcgtcgatttttgtgatgctcgtcaggggggcg
gagcctatggaaaaacgccagcaacgcggcctttttacggttcctggccttttgctggcctttt
gctcacatgttctttcctgcgttatcccctgattctgtggataaccgtattaccgcctttgagt
gagctgataccgctcgccgcagccgaacgaccgagcgcagcgagtcagtgagcgaggaagcgga
agagcgcccaatacgcaaaccgcctctccccgcgcgttggccgattcattaatgcagctggcac
gacaggtttcccgactggaaagcgggcagtgagcgcaacgcaattaatgtgagttagctcactc
attaggcaccccaggctttacactttatgcttccggctcgtatgttgtgtggaattgtgagcgg
ataacaatttcacacaggaaacagctatgaccatgattacgccaagctcgaaattaaccctcac
taaagggaacaaaagctggtaccgggccccccctcgaggtcgacggtatcgataagcttgatat
cgaattcctgcagcccgggggatccactagttctagagcggccgccaccgcggtggagctccaa
ttcgccctatagtgagtcgtattacaattcactggccgtcgttttacaacgtcgtgactgggaa
aaccctggcgttacccaacttaatcgccttgcagcacatccccctttcgccagctggcgtaata
gcgaagaggcccgcaccgatcgcccttcccaacagttgcgcagcctgaatggcgaatggacgcg
ccctgtagcggcgcattaagcgcggcgggtgtggtggttacgcgcagcgtgaccgctacacttg
ccagcgccctagcgcccgctcctttcgctttcttcccttcctttctcgccacgttcgccggctt
tccccgtcaagctctaaatcgggggctccctttagggttccgatttagtgctttacggcacctc
gaccccaaaaaacttgattagggtgatggttcacgtagtgggccatcgccctgatagacggttt
ttcgccctttgacgttggagtccacgttctttaatagtggactcttgttccaaactggaacaac
actcaaccctatctcggtctattcttttgatttataagggattttgccgatttcggcctattgg
ttaaaaaatgagctgatttaacaaaaatttaacgcgaattttaacaaaatattaacgcttacaa
tttcctgatgcggtattttctccttacgcatctgtgcggtatttcacaccgcatagatccgtcg
agttcaagagaaaaaaaaagaaaaagcaaaaagaaaaaaggaaagcgcgcctcgttcagaatga
cacgtatagaatgatgcattaccttgtcatcttcagtatcatactgttcgtatacatacttact
gacattcataggtatacatatatacacatgtatatatatcgtatgctgcagctttaaataatcg
gtgtcactacataagaacacctttggtggagggaacatcgttggtaccattgggcgaggtggct
tctcttatggcaaccgcaagagccttgaacgcactctcactacggtgatgatcattcttgcctc
gcagacaatcaacgtggagggtaattctgctagcctctgcaaagctttcaagaaaatgcgggat
catctcgcaagagagatctcctactttctccctttgcaaaccaagttcgacaactgcgtacggc

ctgttcgaaagatctaccaccgctctggaaagtgcctcatccaaaggcgcaaatcctgatccaa
acctttttactccacgcacggcccctagggcctctttaaaagcttgaccgagagcaatcccgca
gtcttcagtggtgtgatggtcgtctatgtgtaagtcaccaatgcactcaacgattagcgaccag
ccggaatgcttggccagagcatgtatcatatggtccagaaaccctatacctgtgtggacgttaa
tcacttgcgattgtgtggcctgttctgctactgcttctgcctctttttctgggaagatcgagtg
ctctatcgctaggggaccaccctttaaagagatcgcaatctgaatcttggtttcatttgtaata
cgctttactagggctttctgctctgtcatctttgccttcgtttatcttgcctgctcatttttta
gtatattcttcgaagaaatcacattactttatataatgtataattcattatgtgataatgccaa
tcgctaagaaaaaaaagagtcatccgctaggtggaaaaaaaaaaatgaaaatcattaccgaggca
taaaaaaatatagagtgtactagaggaggccaagagtaatagaaaaagaaaattgcgggaaagg
actgtgttatgacttccctgactaatgccgtgttcaaacgatacctggcagtgactcctagcgc
tcaccaagctcttaaaacgggaatttatggtgcactctcagtacaatctgctctgatgccgcat
agttaagccagccccgacacccgccaacacccgctgacgcgccctgacgggcttgtctgctccc
ggcatccgcttacagacaagctgtgaccgtctccgggagctgcatgtgtcagaggttttcaccg
tcatcaccgaaacgcgcgagacgaaagggcctcgtgatacgcctatttttataggttaatgtca
tgataataatggtttcttagggtcctttcatcacgtgctataaaaataattataatttaaatt
ttttaatataaatatataaattaaaaatagaaagtaaaaaaagaaattaaagaaaaaatagttt
ttgttttccgaagatgtaaaagactctagggggatcgccaacaaatactacctttatcttgct
cttcctgctctcaggtattaatgccgaattgtttcatcttgtctgtgtagaagaccacacacga
aaatcctgtgattttacattttacttatcgttaatcgaatgtatatctatttaatctgctttc
ttgtctaataaatatatatgtaaagtacgctttttgttgaaattttttaaacctttgtttattt
ttttttcttcattccgtaactcttctaccttctttatttactttctaaaatccaaatacaaaac
ataaaaataaataaacacagagtaaattcccaaattattccatcattaaaagatacgaggcgcg
tgtaagttacaggcaagcgatccgtcaggtggcacttttcggggaaatgtgcgcggaacccta
tttgtttatttttctaaatacattcaaatatgtatccgctcatgagacaataaccctgataaat
gcttcaataatattgaaaaaggaagagtatgagtattcaacatttccgtgtcgcccttattccc
ttttttgcggcattttgccttcctgtttttgctcacccagaaacgctggtgaaagtaaaagatg
ctgaagatcagttgggtgcacgagtgggttacatcgaactggatctcaacagcggtaagatcct
tgagagttttcgccccgaagaacgttttccaatgatgagcacttttaaagttctgctatgtggc
gcggtattatcccgtattgacgccgggcaagagcaactcggtcgccgcatacactattctcaga
atgacttggttgagtactcaccagtcacagaaaagcatcttacggatggcatgacagtaagaga
attatgcagtgctgccataaccatgagtgataacactgcggccaacttacttctgacaacgatc
ggaggaccgaaggagctaaccgcttttttgcacaacatgggggatcatgtaactcgccttgatc
gttgggaaccggagctgaatgaagccataccaaacgacgagcgtgacaccacgatgcctgtagc
aatggcaacaacgttgcgcaaactattaactggcgaactacttactctagcttcccggcaacaa
ttaatagactggatggaggcggataaagttgcaggaccacttctgcgctcggcccttccggctg
gctggtttattgctgataaatctggagccggtgagcgtgggtctcgcggtatcattgcagcact
ggggccagatggtaagccctcccgtatcgtagttatctacacgacggggagtcaggcaactatg
gatgaacgaaatagacagatcgctgagataggtgcctcactgattaagcattggtaactgtcag
accaagtttactcatatatactttagattgatttaaaacttcatttttaatttaaaaggatcta

ggtgaagatcctttttgataatctcatgaccaaaatcccttaacgtgagttttcgttccactga
gcgtcagaccccgtagaaaagatcaaaggatcttc

**pVJ27 (pRS316)**

ttgagatcctttttttctgcgcgtaatctgctgcttgcaaacaaaaaaaccaccgctaccagcg
gtggtttgtttgccggatcaagagctaccaactctttttccgaaggtaactggcttcagcagag
cgcagataccaaatactgttcttctagtgtagccgtagttaggccaccacttcaagaactctgt
agcaccgcctacatacctcgctctgctaatcctgttaccagtggctgctgccagtggcgataag
tcgtgtcttaccgggttggactcaagacgatagttaccggataaggcgcagcggtcgggctgaa
cggggggttcgtgcacacagcccagcttggagcgaacgacctacaccgaactgagatacctaca
gcgtgagctatgagaaagcgccacgcttcccgaagggagaaaggcggacaggtatccggtaagc
ggcagggtcggaacaggagagcgcacgagggagcttccagggggaaacgcctggtatctttata
gtcctgtcgggtttcgccacctctgacttgagcgtcgatttttgtgatgctcgtcaggggggcg
gagcctatggaaaaacgccagcaacgcggcctttttacggttcctggccttttgctggcctttt
gctcacatgttctttcctgcgttatcccctgattctgtggataaccgtattaccgcctttgagt
gagctgataccgctcgccgcagccgaacgaccgagcgcagcgagtcagtgagcgaggaagcgga
agagcgcccaatacgcaaaccgcctctccccgcgcgttggccgattcattaatgcagctggcac
gacaggtttcccgactggaaagcgggcagtgagcgcaacgcaattaatgtgagttagctcactc
attaggcaccccaggctttacactttatgcttccggctcgtatgttgtgtggaattgtgagcgg
ataacaatttcacacaggaaacagctatgaccatgattacgccaagctcgaaattaaccctcac
taaagggaacaaaagctggtaccgggccccccctcgaggtcgacggtatcgataagcttgatat
cgaattcctgcagcccgggggatccactagttctagagcggccgccaccgcggtggagctccaa
ttcgccctatagtgagtcgtattacaattcactggccgtcgttttacaacgtcgtgactgggaa
aaccctggcgttacccaacttaatcgccttgcagcacatccccctttcgccagctggcgtaata
gcgaagaggcccgcaccgatcgcccttcccaacagttgcgcagcctgaatggcgaatggacgcg
ccctgtagcggcgcattaagcgcggcgggtgtggtggttacgcgcagcgtgaccgctacacttg
ccagcgccctagcgcccgctcctttcgctttcttcccttcctttctcgccacgttcgccggctt
tccccgtcaagctctaaatcgggggctccctttagggttccgatttagtgctttacggcacctc
gaccccaaaaaacttgattagggtgatggttcacgtagtgggccatcgccctgatagacggttt
ttcgccctttgacgttggagtccacgttctttaatagtggactcttgttccaaactggaacaac
actcaaccctatctcggtctattcttttgatttataagggattttgccgatttcggcctattgg
ttaaaaaatgagctgatttaacaaaaatttaacgcgaattttaacaaaatattaacgcttacaa
tttcctgatgcggtattttctccttacgcatctgtgcggtatttcacaccgcataggtaataa
ctgatataattaaattgaagctctaatttgtgagtttagtatacatgcatttacttataataca
gtttttagttttgctggccgcatcttctcaaatatgcttcccagcctgcttttctgtaacgtt
caccctctaccttagcatcccttccctttgcaaatagtcctcttccaacaataataatgtcaga
tcctgtagagaccacatcatccacggttctatactgttgacccaatgcgtctcccttgtcatct
aaacccacaccgggtgtcataatcaaccaatcgtaaccttcatctcttccacccatgtctcttt
gagcaataaagccgataacaaaatctttgtcgctcttcgcaatgtcaacagtacccttagtata

ttctccagtagatagggagcccttgcatgacaattctgctaacatcaaaaggcctctaggttcc
tttgttacttcttctgccgcctgcttcaaaccgctaacaatacctgggcccaccacaccgtgtg
cattcgtaatgtctgcccattctgctattctgtatacacccgcagagtactgcaatttgactgt
attaccaatgtcagcaaattttctgtcttcgaagagtaaaaaattgtacttggcggataatgcc
tttagcggcttaactgtgccctccatggaaaaatcagtcaagatatccacatgtgtttttagta
aacaaattttgggacctaatgcttcaactaactccagtaattccttggtggtacgaacatccaa
tgaagcacacaagtttgtttgcttttcgtgcatgatattaaatagcttggcagcaacaggacta
ggatgagtagcagcacgttccttatatgtagctttcgacatgatttatcttcgtttcctgcagg
tttttgttctgtgcagttgggttaagaatactgggcaatttcatgtttcttcaacactacatat
gcgtatatataccaatctaagtctgtgctccttccttcgttcttccttctgttcggagattacc
gaatcaaaaaaatttcaaggaaaccgaaatcaaaaaaaagaataaaaaaaaaatgatgaattga
aaaggtggtatggtgcactctcagtacaatctgctctgatgccgcatagttaagccagccccga
cacccgccaacacccgctgacgcgccctgacgggcttgtctgctcccggcatccgcttacagac
aagctgtgaccgtctccgggagctgcatgtgtcagaggttttcaccgtcatcaccgaaacgcgc
gagacgaaagggcctcgtgatacgcctatttttataggttaatgtcatgataataatggtttct
tagacggatcgcttgcctgtaacttacacgcgcctcgtatcttttaatgatggaataatttggg
aatttactctgtgtttatttatttttatgttttgtatttggattttagaaagtaaataaagaag
gtagaagagttacggaatgaagaaaaaaaaataaacaaaggtttaaaaaatttcaacaaaaagc
gtactttacatatatatttattagacaagaaaagcagattaaatagatatacattcgattaacg
ataagtaaaatgtaaaatcacaggattttcgtgtgtggtcttctacacagacaagatgaaacaa
ttcggcattaatacctgagagcaggaagagcaagataaaaggtagtatttgttggcgatccccc
tagagtcttttacatcttcggaaaacaaaaactattttttctttaatttctttttttactttct
atttttaatttatatatttatattaaaaaatttaaattataattatttttatagcacgtgatga
aaaggacccaggtggcacttttcggggaaatgtgcgcggaacccctatttgtttatttttctaa
atacattcaaatatgtatccgctcatgagacaataaccctgataaatgcttcaataatattgaa
aaaggaagagtatgagtattcaacatttccgtgtcgcccttattcccttttttgcggcattttg
ccttcctgtttttgctcacccagaaacgctggtgaaagtaaaagatgctgaagatcagttgggt
gcacgagtgggttacatcgaactggatctcaacagcggtaagatccttgagagttttcgccccg
aagaacgttttccaatgatgagcacttttaaagttctgctatgtggcgcggtattatcccgtat
tgacgccgggcaagagcaactcggtcgccgcatacactattctcagaatgacttggttgagtac
tcaccagtcacagaaaagcatcttacggatggcatgacagtaagagaattatgcagtgctgcca
taaccatgagtgataacactgcggccaacttacttctgacaacgatcggaggaccgaaggagct
aaccgcttttttgcacaacatgggggatcatgtaactcgccttgatcgttgggaaccggagctg
aatgaagccataccaaacgacgagcgtgacaccacgatgcctgtagcaatggcaacaacgttgc
gcaaactattaactggcgaactacttactctagcttcccggcaacaattaatagactggatgga
ggcggataaagttgcaggaccacttctgcgctcggcccttccggctggctggtttattgctgat
aaatctggagccggtgagcgtgggtctcgcggtatcattgcagcactggggccagatggtaagc
cctcccgtatcgtagttatctacacgacggggagtcaggcaactatggatgaacgaaatagaca
gatcgctgagataggtgcctcactgattaagcattggtaactgtcagaccaagtttactcatat
atactttagattgatttaaaacttcatttttaatttaaaaggatctaggtgaagatcctttttg

ataatctcatgaccaaaatcccttaacgtgagttttcgttccactgagcgtcagaccccgtaga
aaagatcaaaggatcttc

**pVJ317 (p416MET25-Deg1*-Sec62-ProtA)**

ttgagatcctttttttctgcgcgtaatctgctgcttgcaaacaaaaaaaccaccgctaccagcg
gtggtttgtttgccggatcaagagctaccaactctttttccgaaggtaactggcttcagcagag
cgcagataccaaatactgttcttctagtgtagccgtagttaggccaccacttcaagaactctgt
agcaccgcctacatacctcgctctgctaatcctgttaccagtggctgctgccagtggcgataag
tcgtgtcttaccgggttggactcaagacgatagttaccggataaggcgcagcggtcgggctgaa
cggggggttcgtgcacacagcccagcttggagcgaacgacctacaccgaactgagatacctaca
gcgtgagctatgagaaagcgccacgcttcccgaagggagaaaggcggacaggtatccggtaagc
ggcagggtcggaacaggagagcgcacgagggagcttccagggggaaacgcctggtatctttata
gtcctgtcgggtttcgccacctctgacttgagcgtcgatttttgtgatgctcgtcaggggggcg
gagcctatggaaaaacgccagcaacgcggcctttttacggttcctggccttttgctggcctttt
gctcacatgttctttcctgcgttatcccctgattctgtggataaccgtattaccgcctttgagt
gagctgataccgctcgccgcagccgaacgaccgagcgcagcgagtcagtgagcgaggaagcgga
agagcgcccaatacgcaaaccgcctctccccgcgcgttggccgattcattaatgcagctggcac
gacaggtttcccgactggaaagcgggcagtgagcgcaacgcaattaatgtgagttagctcactc
attaggcaccccaggctttacactttatgcttccggctcgtatgttgtgtggaattgtgagcgg
ataacaatttcacacaggaaacagctatgaccatgattacgccaagcgcgcaattaaccctcac
taaagggaacaaaagctggagctccggatgcaagggttcgaatcccttagctctcattatttt
tgctttttctcttgaggtcacatgatcgcaaaatggcaaatggcacgtgaagctgtcgatattg
gggaactgtggtggttggcaaatgactaattaagttagtcaaggcgccatcctcatgaaaactg
tgtaacataataaccgaagtgtcgaaaaggtggcaccttgtccaattgaacacgctcgatgaaa
aaaataagatatatataaggttaagtaaagcgtctgttagaaaggaagtttttcctttttcttg
ctctcttgtcttttcatctactatttccttcgtgtaatacagggtcgtcagatacatagataca
attctattacccccatccatactctagaactagtggatccatgaataaaatacccattaaagac
cttttaaatccacaaatcacagatgagtctaaatccagcacactagacataaataaaaagctct
tttctatttgctgtaatttacctaagttaccagagagtgtaacaacagaagaagaagttgaatt
aagggatatattaggattcttatctagggccaacaaaaaccgtaagattgagctcgactacaag
gacgacgatgacaaggagctcggtaccatggtagccgagcaaacacaggagaacatgtcagccg
taggtccaggtagcaatgcaggtgctagtgtcaatggcggatctgctacagccattgctactct
tttacgcaaccataaggagttgaagcaaaggcagggtttattccaggccaagcagacagacttc
ttccgttacaagagatttgttagggcactgcattctgaggagtatgccaacaaatctgcaagac
agccagagatatatccaactataccttctaacaagatcgaagaccagcttaagtcacgtgaaat
ctttattcaacttataaaggcacaaatggtgatcccagtgaaaaaattgcatagtcaagaatgc
aaagagcatggattgaagccaagcaaggactttccgcacttgattgtttcgaataaagcacaat
tggaagccgatgaatactttgtttggaactataaccctagaacctacatggattacttgatcgt
cattggtgtcgtgtccatcatattggcgctcgtatgctacccattgtggccgcgctccatgaga
cgcggctcctattatgtgtctctgggtgccttcggcatcttagcaggcttctttgctgtcgcta
tcctgagattgatattatatgttttgtcattaattgtctacaaagatgttggtgggttctggat
ctttcccaacctgttcgaagattgcggtgtactagagagttttaagccactttacggttttggt
gagaaggatacttatagttacaaaaagaaactgaaaagaatgaagaagaaacaagccaagagag

aaagcaataagaagaaagccatcaatgaaaaagccgaacaaaacggtaccggttctggtcttgc
gcaacacgatgaagccgtggacaacaaattcaacaaagaacaacaaaacgcgttctatgagatc
ttacatttacctaacttaaacgaagaacaacgaaacgccttcatccaaagtttaaaagatgacc
caagccaaagcgctaacctttagcagaagctaaaaagctaaatgatgctcaggcgccgaaagt
agacaacaaattcaacaaagaacaacaaaacgcgttctatgagatcttacatttacctaactta
aacgaagaacaacgaaacgccttcatccaaagtttaaaagatgacccaagccaaagcgctaacc
ttttagcagaagctaaaaagctaaatgatgctcaggcgccgaaagtagacgcgctgcagccaag
ctaattccgggcgaatttcttatgatttatgatttttattattaaataagttataaaaaaaata
agtgtatacaaattttaaagtgactcttaggttttaaaacgaaaattcttattcttgagtaact
ctttcctgtaggtcaggttgctttctcaggtatagcatgaggtcgctcttattgaccacacctc
taccggcatgcaagcttatcgataccgtcgacctcgagtcatgtaattagttatgtcacgctta
cattcacgccctccccccacatccgctctaaccgaaaaggaaggagttagacaacctgaagtct
aggtccctatttatttttttatagttatgttagtattaagaacgttatttatatttcaaatttt
tctttttttctgtacagacgcgtgtacgcatgtaacattatactgaaaaccttgcttgagaag
gttttgggacgctcgaaggctttaatttgcggccggtacccaattcgccctatagtgagtcgta
ttacgcgcgctcactggccgtcgttttacaacgtcgtgactgggaaaaccctggcgttaccaa
cttaatcgccttgcagcacatccccctttcgccagctggcgtaatagcgaagaggcccgcaccg
atcgcccttcccaacagttgcgcagcctgaatggcgaatggacgcgccctgtagcggcgcatta
agcgcggcgggtgtggtggttacgcgcagcgtgaccgctacacttgccagcgccctagcgcccg
ctcctttcgctttcttcccttcctttctcgccacgttcgccggctttccccgtcaagctctaaa
tcgggggctccctttagggttccgatttagtgctttacggcacctcgaccccaaaaaacttgat
tagggtgatggttcacgtagtgggccatcgccctgatagacggtttttcgccctttgacgttgg
agtccacgttctttaatagtggactcttgttccaaactggaacaacactcaaccctatctcggt
ctattcttttgatttataagggattttgccgatttcggcctattggttaaaaaatgagctgatt
taacaaaaatttaacgcgaattttaacaaaatattaacgcttacaatttcctgatgcggtattt
tctccttacgcatctgtgcggtatttcacaccgcatagggtaataactgatataattaaattga
agctctaatttgtgagtttagtatacatgcatttacttataatacagtttttttagttttgctgg
ccgcatcttctcaaatatgcttcccagcctgcttttctgtaacgttcaccctctaccttagcat
cccttccctttgcaaatagtcctcttccaacaataataatgtcagatcctgtagagaccacatc
atccacggttctatactgttgacccaatgcgtctcccttgtcatctaaacccacaccgggtgtc
ataatcaaccaatcgtaaccttcatctcttccacccatgtctctttgagcaataaagccgataa
caaaatctttgtcgctcttcgcaatgtcaacagtacccttagtatattctccagtagatagga
gcccttgcatgacaattctgctaacatcaaaaggcctctaggttcctttgttacttcttctgcc
gcctgcttcaaaccgctaacaatacctgggcccaccacaccgtgtgcattcgtaatgtctgccc
attctgctattctgtatacacccgcagagtactgcaatttgactgtattaccaatgtcagcaaa
ttttctgtcttcgaagagtaaaaaattgtacttggcggataatgcctttagcggcttaactgtg
ccctccatggaaaaatcagtcaagatatccacatgtgtttttagtaaacaaattttgggaccta
atgcttcaactaactccagtaattccttggtggtacgaacatccaatgaagcacacaagtttgt
ttgcttttcgtgcatgatattaaatagcttggcagcaacaggactaggatgagtagcagcacgt
tccttatatgtagctttcgacatgatttatcttcgtttcctgcaggtttttgttctgtgcagtt
gggttaagaatactgggcaatttcatgtttcttcaacactacatatgcgtatatataccaatct

aagtctgtgctccttccttcgttcttccttctgttcggagattaccgaatcaaaaaaatttcaa
ggaaaccgaaatcaaaaaaaagaataaaaaaaaaatgatgaattgaaaaggtggtatggtgcac
tctcagtacaatctgctctgatgccgcatagttaagccagccccgacacccgccaacacccgct
gacgcgccctgacgggcttgtctgctcccggcatccgcttacagacaagctgtgaccgtctccg
ggagctgcatgtgtcagaggttttcaccgtcatcaccgaaacgcgcgagacgaaagggcctcgt
gatacgcctatttttataggttaatgtcatgataataatggtttcttagacggatcgcttgcct
gtaacttacacgcgcctcgtatcttttaatgatggaataatttgggaatttactctgtgtttat
ttatttttatgttttgtatttggattttagaaagtaaataaagaaggtagaagagttacggaat
gaagaaaaaaaaataaacaaaggtttaaaaaatttcaacaaaaagcgtactttacatatatatt
tattagacaagaaaagcagattaaatagatatacattcgattaacgataagtaaaatgtaaaat
cacaggattttcgtgtgtggtcttctacacagacaagatgaaacaattcggcattaatacctga
gagcaggaagagcaagataaaaggtagtatttgttggcgatccccctagagtcttttacatctt
cggaaaacaaaaactatttttctttaattgctttttttactttctatttttaatttatatatt
tatattaaaaaatttaaattataattatttttatagcacgtgatgaaaaggacccaggtggcac
ttttcggggaaatgtgcgcggaacccctatttgtttatttttctaaatacattcaaatatgtat
ccgctcatgagacaataaccctgataaatgcttcaataatattgaaaaaggaagagtatgagta
ttcaacatttccgtgtcgcccttattcccttttttgcggcattttgccttcctgtttttgctca
cccagaaacgctggtgaaagtaaaagatgctgaagatcagttgggtgcacgagtgggttacatc
gaactggatctcaacagcggtaagatccttgagagttttcgccccgaagaacgttttccaatga
tgagcacttttaaagttctgctatgtggcgcggtattatcccgtattgacgccgggcaagagca
actcggtcgccgcatacactattctcagaatgacttggttgagtactcaccagtcacagaaaag
catcttacggatggcatgacagtaagagaattatgcagtgctgccataaccatgagtgataaca
ctgcggccaacttacttctgacaacgatcggaggaccgaaggagctaaccgcttttttgcacaa
catgggggatcatgtaactcgccttgatcgttgggaaccggagctgaatgaagccataccaaac
gacgagcgtgacaccacgatgcctgtagcaatggcaacaacgttgcgcaaactattaactggcg
aactacttactctagcttcccggcaacaattaatagactggatggaggcggataaagttgcagg
accacttctgcgctcggcccttccggctggctggtttattgctgataaatctggagccggtgag
cgtgggtctcgcggtatcattgcagcactggggccagatggtaagccctcccgtatcgtagtta
tctacacgacggggagtcaggcaactatggatgaacgaaatagacagatcgctgagataggtgc
ctcactgattaagcattggtaactgtcagaccaagtttactcatatatactttagattgattta
aaacttcatttttaatttaaaaggatctaggtgaagatcctttttgataatctcatgaccaaaa
tcccttaacgtgagttttcgttccactgagcgtcagaccccgtagaaaagatcaaaggatcttc

**pVJ382 (YCPlac33-$P_{GPD}$-GFP-2A-Flag-HIS3)**

ttgagatcctttttttctgcgcgtaatctgctgcttgcaaacaaaaaaaccaccgctaccagcg
gtggtttgtttgccggatcaagagctaccaactctttttccgaaggtaactggcttcagcagag
cgcagataccaaatactgttcttctagtgtagccgtagttaggccaccacttcaagaactctgt
agcaccgcctacatacctcgctctgctaatcctgttaccagtggctgctgccagtggcgataag
tcgtgtcttaccgggttggactcaagacgatagttaccggataaggcgcagcggtcgggctgaa
cggggggttcgtgcacacagcccagcttggagcgaacgacctacaccgaactgagatacctaca
gcgtgagctatgagaaagcgccacgcttcccgaagggagaaaggcggacaggtatccggtaagc
ggcagggtcggaacaggagagcgcacgagggagcttccagggggaaacgcctggtatctttata
gtcctgtcgggtttcgccacctctgacttgagcgtcgatttttgtgatgctcgtcaggggggcg
gagcctatggaaaaacgccagcaacgcggcctttttacggttcctggccttttgctggcctttt
gctcacatgttctttcctgcgttatcccctgattctgtggataaccgtattaccgcctttgagt
gagctgataccgctcgccgcagccgaacgaccgagcgcagcgagtcagtgagcgaggaagcgga
agagcgcccaatacgcaaaccgcctctccccgcgcgttggccgattcattaatgcagctggcac
gacaggtttcccgactggaaagcgggcagtgagcgcaacgcaattaatgtgagttagctcactc
attaggcaccccaggctttacactttatgcttccggctcgtatgttgtgtggaattgtgagcgg
ataacaatttcacacaggaaacagctatgaccatgattacgccaagcttgcatgcctgcaggtc
gactctagaggatccccgggtaccgagctcagtttatcattatcaatactcgccatttcaaaga
atacgtaaataattaatagtagtgattttcctaactttatttagtcaaaaaattagccttttaa
ttctgctgtaacccgtacatgcccaaaatagggggcgggttacacagaatatataacatcgtag
gtgtctgggtgaacagtttattcctggcatccactaaatataatggagcccgctttttaagctg
gcatccagaaaaaaaaagaatcccagcaccaaaatattgttttcttcaccaaccatcagttcat
aggtccattctcttagcgcaactacagagaacaggggcacaaacaggcaaaaaacgggcacaac
ctcaatggagtgatgcaacctgcctggagtaaatgatgacacaaggcaattgacccacgcatgt
atctatctcattttcttacaccttctattaccttctgctctctctgatttggaaaaagctgaaa
aaaaaggttgaaaccagttccctgaaattattcccctacttgactaataagtatataaagacgg
taggtattgattgtaattctgtaaatctatttcttaaacttcttaaattctacttttatagtta
gtcttttttttagttttaaaacaccagaacttagtttcgacggattctagaATGAGTAAAGGAG
AAGAACTTTTCACTGGAGTTGTCCCAATTCTTGTTGAATTAGATGGTGATGTTAATGGGCACAA
ATTTTCTGTCAGTGGAGAGGGTGAAGGTGATGCAACATACGGAAAACTTACCCTTAAATTTATT
TGCACTACTGGAAAACTACCTGTTCCATGGCCAACACTTGTCACTACTCTGACGTATGGTGTTC
AATGCTTTTCCCGTTATCCGGATCATATGAAACGGTATGACTTTTTCAAGAGTGCCATGCCCGA
AGGTTATGTACAGGAACGCACTATATCTTTCAAAGATGACGGGAACTACAAGACGCGTGCTGAA
GTCAAGTTTGAAGGTGATACCCTTGTTAATCGTATCGAGTTAAAAGGTATTGATTTTAAAGAAG
ATGGAAACATTCTCGGACACAAACTCGAGTACAACTATAACTCACACAATGTATACATCACGGC
AGACAAACAAAAGAATGGAATCAAAGCTAACTTCAAAATTCGCCACAACATTGAAGATGGATCC
GTTCAACTAGCAGACCATTATCAACAAAATACTCCAATTGGCGATGGCCCTGTCCTTTTACCAG
ACAACCATTACCTGTCGACACAATCTGCCCTTTTGAAAGATCCCAACGAAAAGCGTGACCACAT
GGTCCTTCTTGAGTTTGTAACTGCTGCTGGGATTACACATGGCATGGATGAACTATACAAAACT

AGCCAGCTGTTGAATTTTGACCTTCTTAAGCTTGCGGGAGACGTCGAGTCCAACCCTGGGCCCA
CTAGTGGACCTGGTGACTACAAGGACGACGATGACAAGGGTCCTGGTATGACAGAGCAGAAAGC
CCTAGTAAAGCGTATTACAAATGAAACCAAGATTCAGATTGCGATCTCTTTAAAGGGTGGTCCC
CTAGCGATAGAGCACTCGATCTTCCCAGAAAAAGAGGCAGAAGCAGTAGCAGAACAGGCCACAC
AATCGCAAGTGATTAACGTCCACACAGGTATAGGGTTTCTGGACCATATGATACATGCTCTGGC
CAAGCATTCCGGCTGGTCGCTAATCGTTGAGTGCATTGGTGACTTACACATAGACGACCATCAC
ACCACTGAAGACTGCGGGATTGCTCTCGGTCAAGCTTTTAAAGAGGCCCTAGGGGCCGTGCGTG
GAGTAAAAAGGTTTGGATCAGGATTTGCGCCTTTGGATGAGGCACTTTCCAGAGCGGTGGTAGA
TCTTTCGAACAGGCCGTACGCAGTTGTCGAACTTGGTTTGCAAAGGGAGAAAGTAGGAGATCTC
TCTTGCGAGATGATCCCGCATTTTCTTGAAAGCTTTGCAGAGGCTAGCAGAATTACCCTCCACG
TTGATTGTCTGCGAGGCAAGAATGATCATCACCGTAGTGAGAGTGCGTTCAAGGCTCTTGCGGT
TGCCATAAGAGAAGCCACCTCGCCCAATGGTACCAACGATGTTCCCTCCACCAAAGGTGTTCTT
ATGggatcctagtgacaccgattatttaaagctgcagcatacgatatatatacatgtgtatata
tgtatacctatgaatgtcagtaagtatgtatacgaacagtatgatactgaagatgacaaggtaa
tgcatcattctatacgtgtcattctgaacgaggaattcactggccgtcgttttacaacgtcgtg
actgggaaaaccctggcgttacccaacttaatcgccttgcagcacatccccctttcgccagctg
gcgtaatagcgaagaggcccgcaccgatcgcccttcccaacagttgcgcagcctgaatggcgaa
tggcgcctgatgcggtattttctccttacgcatctgtgcggtatttcacaccgcataatcgctg
ggccattctcatgaagaatatcttgaatttattgtcatattactagttggtgtggaagtcctaa
tatcggtgatcaatatagtggttgacatgctggctagtcaacattgagccttttgatcatgcaa
atatattacggtattttacaatcaaatatcaaacttaactattgactttataacttatttaggt
ggtaacattcttataaaaaagaaaaaaattactgcaaaacagtactagcttttaacttgtatcc
taggttatctatgctgtctcaccatagagaatattacctatttcagaatgtatgtccatgattc
gccgggtaaatacatataatacacaaatctggcttaataaagtctataatatatctcataaaga
agtgctaaattggctagtgctatatatttttaagaaaatttcttttgactaagtccatatcgac
tttgtaaaagttcacattagcatacatatattacacgagccagaaatagtaacttttgcctaaa
tcacaaattgcaaaatttaattgcttgcaaaaggtcacatgcttataatcaactttttaaaaa
tttaaaatactttttatttttttattttttaaacataaatgaaataatttatttattgtttatga
ttaccgaaacataaaacctgctcaagaaaaagaaactgttttgtccttggaaaaaaagcactac
ctaggagcggccaaaatgccgaggctttcatagcttaaactctttacagaaaataggcattata
gatcagttcgagttttcttattcttccttccggttttatcgtcacagttttacagtaaataagt
atcacctcttagagttcgatgataagctgtcaaacatgagaattaattccacatgttaaaatag
tgaaggagcatgttcggcacacagtggaccgaacgtgggtaagtgcactagggtccggttaaa
cggatctcgcattgatgaggcaacgctaattatcaacatatagattgttatctatctgcatgaa
cacgaaatctttacttgacgacttgaggctgatggtgtttatgcaaagaaaccactgtgtttaa
tatgtgtcactgtttgatattactgtcagcgtagaagataatagtaaaagcggttaataagtgt
atttgagataagtgtgataaagtttttacagcgaaaagacgataaatacaagaaaatgattacg
aggatacggagagaggtatgtacatgtgtatttatatactaagctgccggcggttgtttgcaag
accgagaaaaggctagcaagaatcgggtcattgtagcgtatgcgcctgtgaacattctcttcaa
caagtttgattccattgcggtgaaatggtaaaagtcaacccccctgcgatgtatattttcctgta
caatcaatcaaaaagccaaatgatttagcattatctttacatcttgttattttacagattttat

gtttagatcttttatgcttgcttttcaaaaggcttgcaggcaagtgcacaaacaatacttaaat
aaatactactcagtaataacctatttcttagcattttttgacgaaatttgctattttgttagagt
cttttacaccatttgtctccacacctccgcttacatcaacaccaataacgccatttaatctaag
cgcatcaccaacattttctggcgtcagtccaccagctaacataaaatgtaagctctcggggctc
tcttgccttccaacccagtcagaaatcgagttccaatccaaaagttcacctgtcccacctgctt
ctgaatcaaacaagggaataaacgaatgaggtttctgtgaagctgcactgagtagtatgttgca
gtcttttggaaatacgagtcttttaataactggcaaaccgaggaactcttggtattcttgccac
gactcatctccatgcagttggacgatcgatgataagctgtcaaacatgagaattgggtaataac
tgatataattaaattgaagctctaatttgtgagtttagtatacatgcatttacttataatacag
tttttttagttttgctggccgcatcttctcaaatatgcttcccagcctgcttttctgtaacgttc
accctctaccttagcatcccttccctttgcaaatagtcctcttccaacaataataatgtcagat
cctgtagagaccacatcatccacggttctatactgttgacccaatgcgtctcccttgtcatcta
aacccacaccgggtgtcataatcaaccaatcgtaaccttcatctcttccacccatgtctctttg
agcaataaagccgataacaaaatctttgtcgctcttcgcaatgtcaacagtacccttagtatat
tctccagtagatagggagcccttgcatgacaattctgctaacatcaaaaggcctctaggttcct
ttgttacttcttctgccgcctgcttcaaaccgctaacaatacctgggcccaccacaccgtgtgc
attcgtaatgtctgcccattctgctattctgtatacacccgcagagtactgcaatttgactgta
ttaccaatgtcagcaaattttctgtcttcgaagagtaaaaaattgtacttggcggataatgcct
ttagcggcttaactgtgccctccatggaaaaatcagtcaagatatccacatgtgtttttagtaa
acaaattttgggacctaatgcttcaactaactccagtaattccttggtggtacgaacatccaat
gaagcacacaagtttgtttgcttttcgtgcatgatattaaatagcttggcagcaacaggactag
gatgagtagcagcacgttccttatatgtagctttcgacatgatttatcttcgtttcctgcatgt
tttgttctgtgcagttgggttaagaatactgggcaatttcatgtttcttcaacactacatatg
cgtatatataccaatctaagtctgtgctccttccttcgttcttccttctgttcggagattaccg
aatcaaaaaaatttcaaggaaaccgaaatcaaaaaaaagaataaaaaaaaaaatgatgaattgaa
aagctaattcttgaagacgaaagggcctcgtgatacgcctatttttataggttaatgtcatgat
aataatggtttcttagacgtcaggtggcacttttcggggaaatgtgcgcggaacccctatttgt
ttatttttctaaatacattcaaatatgtatccgctcatgagacaataaccctgataaatgcttc
aataatattgaaaaaggaagagtatgagtattcaacatttccgtgtcgcccttattcccttttt
tgcggcattttgccttcctgtttttgctcacccagaaacgctggtgaaagtaaaagatgctgaa
gatcagttgggtgcacgagtgggttacatcgaactggatctcaacagcggtaagatccttgaga
gttttcgccccgaagaacgttttccaatgatgagcacttttaaagttctgctatgtggcgcggt
attatcccgtattgacgccgggcaagagcaactcggtcgccgcatacactattctcagaatgac
ttggttgagtactcaccagtcacagaaaagcatcttacggatggcatgacagtaagagaattat
gcagtgctgccataaccatgagtgataacactgcggccaacttacttctgacaacgatcggagg
accgaaggagctaaccgcttttttgcacaacatgggggatcatgtaactcgccttgatcgttgg
gaaccggagctgaatgaagccataccaaacgacgagcgtgacaccacgatgcctgtagcaatgg
caacaacgttgcgcaaactattaactggcgaactacttactctagcttcccggcaacaattaat
agactggatggaggcggataaagttgcaggaccacttctgcgctcggcccttccggctggctgg
tttattgctgataaatctggagccggtgagcgtgggtctcgcggtatcattgcagcactggggc
cagatggtaagccctcccgtatcgtagttatctacacgacggggagtcaggcaactatggatga

acgaaatagacagatcgctgagataggtgcctcactgattaagcattggtaactgtcagaccaa
gtttactcatatatactttagattgatttaaaacttcatttttaatttaaaaggatctaggtga
agatcctttttgataatctcatgaccaaaatcccttaacgtgagttttcgttccactgagcgtc
agaccccgtagaaaagatcaaaggatcttc

**pVJ464 (p413-PGPD-CPY-ProtA)**

ttgagatcctttttttctgcgcgtaatctgctgcttgcaaacaaaaaaaccaccgctaccagcg
gtggtttgtttgccggatcaagagctaccaactctttttccgaaggtaactggcttcagcagag
cgcagataccaaatactgttcttctagtgtagccgtagttaggccaccacttcaagaactctgt
agcaccgcctacatacctcgctctgctaatcctgttaccagtggctgctgccagtggcgataag
tcgtgtcttaccgggttggactcaagacgatagttaccggataaggcgcagcggtcgggctgaa
cggggggttcgtgcacacagcccagcttggagcgaacgacctacaccgaactgagatacctaca
gcgtgagctatgagaaagcgccacgcttcccgaagggagaaaggcggacaggtatccggtaagc
ggcagggtcggaacaggagagcgcacgagggagcttccagggggaaacgcctggtatctttata
gtcctgtcgggtttcgccacctctgacttgagcgtcgatttttgtgatgctcgtcaggggggcg
gagcctatggaaaaacgccagcaacgcggcctttttacggttcctggccttttgctggcctttt
gctcacatgttctttcctgcgttatcccctgattctgtggataaccgtattaccgcctttgagt
gagctgataccgctcgccgcagccgaacgaccgagcgcagcgagtcagtgagcgaggaagcgga
agagcgcccaatacgcaaaccgcctctccccgcgcgttggccgattcattaatgcagctggcac
gacaggtttcccgactggaaagcgggcagtgagcgcaacgcaattaatgtgagttagctcactc
attaggcaccccaggctttacactttatgcttccggctcgtatgttgtgtggaattgtgagcgg
ataacaatttcacacaggaaacagctatgaccatgattacgccaagcgcgcaattaaccctcac
taaagggaacaaaagctggagctcagtttatcattatcaatactcgccatttcaaagaatacgt
aaataattaatagtagtgattttcctaactttatttagtcaaaaaattagccttttaattctgc
tgtaacccgtacatgcccaaaatagggggcgggttacacagaatatataacatcgtaggtgtct
gggtgaacagtttattcctggcatccactaaatataatggagcccgctttttaagctggcatcc
agaaaaaaaaagaatcccagcaccaaaatattgttttcttcaccaaccatcagttcataggtcc
attctcttagcgcaactacagagaacaggggcacaaacaggcaaaaaacgggcacaacctcaat
ggagtgatgcaacctgcctggagtaaatgatgacacaaggcaattgacccacgcatgtatctat
ctcattttcttacaccttctattaccttctgctctctctgatttggaaaaagctgaaaaaaaag
gttgaaaccagttccctgaaattattcccctacttgactaataagtatataaagacggtaggta
ttgattgtaattctgtaaatctatttcttaaacttcttaaattctacttttatagttagtcttt
tttttagttttaaaacaccagaacttagtttcgacggattctagaactagtatgaaagcattca
ccagtttactatgtggactaggcctgtccactacactcgctaaggccatctcattgcaaagacc
gttgggtctagataaggacgttttgctgcaagctgcggaaaaatttggtttggacctcgacctg
gatcatctcttgaaggagttggactccaatgtattggacgcttgggcccaaatagagcatttgt
acccaaaccaggttatgagccttgaaacttccactaagccaaaattccctgaagcaatcaaaac
gaagaaagactgggactttgtggtcaagaatgacgcaattgaaaactatcagcttcgtgtcaac
aagattaaggaccctaaaatcctgggcattgacccaaatgtcacacagtacacgggttacttgg
atgtggaagacgaggacaagcatttcttcttttggacttttgaaagtagaaacgatcctgcaaa
ggatccggtcatcctttggttgaacggggggtccaggttgttcttcactaaccgggctgttcttt
gaattaggaccctcatccattggacctgatttgaaacccatcgggaacccttactcttggaaca
gcaatgccaccgtgatcttccttgaccagcctgtcaacgttgggttctcgtattccgggtcctc
aggtgtttccaacactgtcgccgctggtaaggatgtctataacttcttggagttgttcttcgat
cagttccctgaatacgtcaacaagggccaagatttccacatcgctggggaatcctacgccggcc

attacatccctgtttttgcctctgaaattttgtctcacaaggacagaaacttcaacttaacctc
cgtcttgatcggaaatggcctcactgacccattgactcagtataactattacgaaccaatggcc
tgtggtgaaggtggcgaaccatctgttttgccctcggaggaatgctctgctatggaagactctt
tggaacgttgtttgggcttgatcgagtcgtgctatgactcgcaatccgtctggtcctgtgttcc
agctaccatttattgtaataacgcccaattggctccttaccaacgtaccggcagaaacgtttac
gatatcaggaaggattgtgaaggtggcaatttgtgctacccaacgttacaagatatcgacgact
acttaaaccaggactacgtcaaagaagctgtcggtgcggaggttgaccactacgaatcctgtaa
cttcgatatcaacagaaatttcctgtttgcgggtgattggatgaagccttaccacaccgccgta
acagatcttttgaatcaagacctacccattctggtatatgcaggcgataaagatttcatctgta
actggttgggtaataaggcgtggacggatgtcttgccatggaagtacgacgaagaatttgcaag
ccaaaaagtacgtaactggactgcttctatcaccgacgaggtcgctggtgaagtcaaatcctac
aagcacttcacctatttgagagtcttcaatggtggccacatggttccatttgacgtccctgaaa
acgccttaagtatggttaacgaatggatccacggtggtttctccttaaccggttctggtcttgc
gcaacacgatgaagccgtggacaacaaattcaacaaagaacaacaaaacgcgttctatgagatc
ttacatttacctaacttaaacgaagaacaacgaaacgccttcatccaaagtttaaaagatgacc
caagccaaagcgctaaccttttagcagaagctaaaaagctaaatgatgctcaggcgccgaaagt
agacaacaaattcaacaaagaacaacaaaacgcgttctatgagatcttacatttacctaactta
aacgaagaacaacgaaacgccttcatccaaagtttaaaagatgacccaagccaaagcgctaacc
ttttagcagaagctaaaaagctaaatgatgctcaggcgccgaaagtagacgcgctgcccgggta
gtgacaccgattatttaaagctgcagcatacgatatatatacatgtgtatatatgtatacctat
gaatgtcagtaagtatgtatacgaacagtatgatactgaagatgacaaggtaatgcatcattct
atacgtgtcattctgaacgaggctcgagtcatgtaattagttatgtcacgcttacattcacgcc
ctccccccacatccgctctaaccgaaaaggaaggagttagacaacctgaagtctaggtccctat
ttatttttttatagttatgttagtattaagaacgttatttatatttcaaatttttctttttttt
ctgtacagacgcgtgtacgcatgtaacattatactgaaaaccttgcttgagaaggttttgggac
gctcgaaggctttaatttgcggccggtacccaattcgccctatagtgagtcgtattacgcgcgc
tcactggccgtcgttttacaacgtcgtgactgggaaaaccctggcgttacccaacttaatcgcc
ttgcagcacatccccctttcgccagctggcgtaatagcgaagaggcccgcaccgatcgcccttc
ccaacagttgcgcagcctgaatggcgaatggacgcgccctgtagcggcgcattaagcgcggcgg
gtgtggtggttacgcgcagcgtgaccgctacacttgccagcgccctagcgcccgctcctttcgc
tttcttcccttcctttctcgccacgttcgccggctttccccgtcaagctctaaatcgggggctc
cctttagggttccgatttagtgctttacggcacctcgaccccaaaaaacttgattagggtgatg
gttcacgtagtgggccatcgccctgatagacggtttttcgccctttgacgttggagtccacgtt
ctttaatagtggactcttgttccaaactggaacaacactcaaccctatctcggtctattctttt
gatttataagggattttgccgatttcggcctattggttaaaaaatgagctgatttaacaaaaat
ttaacgcgaattttaacaaaatattaacgcttacaatttcctgatgcggtattttctccttacg
catctgtgcggtatttcacaccgcatagatccgtcgagttcaagagaaaaaaaaagaaaaagca
aaaagaaaaaaggaaagcgcgcctcgttcagaatgacacgtatagaatgatgcattaccttgtc
atcttcagtatcatactgttcgtatacatacttactgacattcataggtatacatatatacaca
tgtatatatatcgtatgcgctttaaataatcggtgtcactacataagaacacctttggtggagg
gaacatcgttggtaccattgggcgaggtggcttctcttatggcaaccgcaagagccttgaacgc

actctcactacggtgatgatcattcttgcctcgcagacaatcaacgtggagggtaattctgcta
gcctctgcaaagctttcaagaaaatgcgggatcatctcgcaagagagatctcctactttctccc
tttgcaaaccaagttcgacaactgcgtacggcctgttcgaaagatctaccaccgctctggaaag
tgcctcatccaaaggcgcaaatcctgatccaaacctttttactccacgcacggcccctagggcc
tctttaaaagcttgaccgagagcaatcccgcagtcttcagtggtgtgatggtcgtctatgtgta
agtcaccaatgcactcaacgattagcgaccagccggaatgcttggccagagcatgtatcatatg
gtccagaaaccctatacctgtgtggacgttaatcacttgcgattgtgtggcctgttctgctact
gcttctgcctctttttctgggaagatcgagtgctctatcgctaggggaccaccctttaaagaga
tcgcaatctgaatcttggtttcatttgtaatacgctttactagggctttctgctctgtcatctt
tgccttcgtttatcttgcctgctcatttttttagtatattcttcgaagaaatcacattactttat
ataatgtataattcattatgtgataatgccaatcgctaagaaaaaaaaagagtcatccgctagg
tggaaaaaaaaaaaatgaaaatcattaccgaggcataaaaaaatatagagtgtactagaggaggc
caagagtaatagaaaaagaaaattgcgggaaaggactgtgttatgacttccctgactaatgccg
tgttcaaacgatacctggcagtgactcctagcgctcaccaagctcttaaaacgggaatttatgg
tgcactctcagtacaatctgctctgatgccgcatagttaagccagccccgacacccgccaacac
ccgctgacgcgccctgacgggcttgtctgctcccggcatccgcttacagacaagctgtgaccgt
ctccgggagctgcatgtgtcagaggttttcaccgtcatcaccgaaacgcgcgagacgaaagggc
ctcgtgatacgcctatttttataggttaatgtcatgataataatggtttcttagggtccttttc
atcacgtgctataaaaataattataatttaaatttttttaatataaatatataaattaaaaatag
aaagtaaaaaaagaaattaaagaaaaaatagtttttgttttccgaagatgtaaaagactctagg
gggatcgccaacaaatactacctttttatcttgctcttcctgctctcaggtattaatgccgaatt
gtttcatcttgtctgtgtagaagaccacacacgaaaatcctgtgattttacattttacttatcg
ttaatcgaatgtatatctatttaatctgcttttcttgtctaataaatatatatgtaaagtacgc
tttttgttgaaattttttaaacctttgtttattttttttttcttcattccgtaactcttctacct
tctttatttactttctaaaatccaaatacaaaacataaaaaataaataaacacagagtaaattcc
caaattattccatcattaaaagatacgaggcgcgtgtaagttacaggcaagcgatccgtcaggt
ggcacttttcggggaaatgtgcgcggaacccctatttgtttatttttctaaatacattcaaata
tgtatccgctcatgagacaataaccctgataaatgcttcaataatattgaaaaaggaagagtat
gagtattcaacatttccgtgtcgcccttattcccttttttgcggcattttgccttcctgttttt
gctcacccagaaacgctggtgaaagtaaaagatgctgaagatcagttgggtgcacgagtgggtt
acatcgaactggatctcaacagcggtaagatccttgagagttttcgccccgaagaacgttttcc
aatgatgagcacttttaaagttctgctatgtggcgcggtattatcccgtattgacgccgggcaa
gagcaactcggtcgccgcatacactattctcagaatgacttggttgagtactcaccagtcacag
aaaagcatcttacggatggcatgacagtaagagaattatgcagtgctgccataaccatgagtga
taacactgcggccaacttacttctgacaacgatcggaggaccgaaggagctaaccgcttttttg
cacaacatgggggatcatgtaactcgccttgatcgttgggaaccggagctgaatgaagccatac
caaacgacgagcgtgacaccacgatgcctgtagcaatggcaacaacgttgcgcaaactattaac
tggcgaactacttactctagcttcccggcaacaattaatagactggatggaggcggataaagtt
gcaggaccacttctgcgctcggcccttccggctggctggtttattgctgataaatctggagccg
gtgagcgtgggtctcgcggtatcattgcagcactggggccagatggtaagccctcccgtatcgt
agttatctacacgacggggagtcaggcaactatggatgaacgaaatagacagatcgctgagata

ggtgcctcactgattaagcattggtaactgtcagaccaagtttactcatatatactttagattg
atttaaaacttcatttttaatttaaaaggatctaggtgaagatcctttttgataatctcatgac
caaaatcccttaacgtgagttttcgttccactgagcgtcagaccccgtagaaaagatcaaagga
tcttc

**pVJ476 (pRS316-Hrd1-3HA)**

gaagatcctttgatcttttctacggggtctgacgctcagtggaacgaaaactcacgttaaggga
ttttggtcatgagattatcaaaaaggatcttcacctagatccttttaaattaaaaatgaagttt
taaatcaatctaaagtatatatgagtaaacttggtctgacagttaccaatgcttaatcagtgag
gcacctatctcagcgatctgtctatttcgttcatccatagttgcctgactccccgtcgtgtaga
taactacgatacgggagggcttaccatctggccccagtgctgcaatgataccgcgagacccacg
ctcaccggctccagatttatcagcaataaaccagccagccggaagggccgagcgcagaagtggt
cctgcaactttatccgcctccatccagtctattaattgttgccgggaagctagagtaagtagtt
cgccagttaatagtttgcgcaacgttgttgccattgctacaggcatcgtggtgtcacgctcgtc
gtttggtatggcttcattcagctccggttcccaacgatcaaggcgagttacatgatcccccatg
ttgtgcaaaaaagcggttagctccttcggtcctccgatcgttgtcagaagtaagttggccgcag
tgttatcactcatggttatggcagcactgcataattctcttactgtcatgccatccgtaagatg
cttttctgtgactggtgagtactcaaccaagtcattctgagaatagtgtatgcggcgaccgagt
tgctcttgcccggcgtcaatacgggataataccgcgccacatagcagaactttaaaagtgctca
tcattggaaaacgttcttcggggcgaaaactctcaaggatcttaccgctgttgagatccagttc
gatgtaacccactcgtgcacccaactgatcttcagcatcttttactttcaccagcgtttctggg
tgagcaaaaacaggaaggcaaaatgccgcaaaaaagggaataagggcgacacggaaatgttgaa
tactcatactcttcctttttcaatattattgaagcatttatcagggttattgtctcatgagcgg
atacatatttgaatgtatttagaaaaataaacaaataggggttccgcgcacatttccccgaaaa
gtgccacctgggtccttttcatcacgtgctataaaaataattataatttaaattttttaatata
aatatataaattaaaaatagaaagtaaaaaaagaaattaaagaaaaaatagtttttgttttccg
aagatgtaaaagactctagggggatcgccaacaaatactaccttttatcttgctcttcctgctc
tcaggtattaatgccgaattgtttcatcttgtctgtgtagaagaccacacacgaaaatcctgtg
attttacattttacttatcgttaatcgaatgtatatctatttaatctgcttttcttgtctaata
aatatatatgtaaagtacgcttttgttgaaattttttaaacctttgtttatttttttttcttc
attccgtaactcttctaccttctttatttactttctaaaatccaaatacaaaacataaaaataa
ataaacacagagtaaattcccaaattattccatcattaaaagatacgaggcgcgtgtaagttac
aggcaagcgatccgtctaagaaaccattattatcatgacattaacctataaaaataggcgtatc
acgaggccctttcgtctcgcgcgtttcggtgatgacggtgaaaacctctgacacatgcagctcc
cggagacggtcacagcttgtctgtaagcggatgccgggagcagacaagcccgtcagggcgcgtc
agcgggtgttggcgggtgtcggggctggcttaactatgcggcatcagagcagattgtactgaga
gtgcaccataccaccttttcaattcatcatttttttttattcttttttttgatttcggtttcc
ttgaaatttttttgattcggtaatctccgaacagaaggaagaacgaaggaaggagcacagactt
agattggtatatatacgcatatgtagtgttgaagaaacatgaaattgcccagtattcttaaccc
aactgcacagaacaaaaacctgcaggaaacgaagataaatcatgtcgaaagctacatataagga
acgtgctgctactcatcctagtcctgttgctgccaagctatttaatatcatgcacgaaaagcaa
acaaacttgtgtgcttcattggatgttcgtaccaccaaggaattactggagttagttgaagcat
taggtcccaaaatttgtttactaaaaacacatgtggatatcttgactgatttttccatggaggg
cacagttaagccgctaaaggcattatccgccaagtacaattttttactcttcgaagacagaaaa
tttgctgacattggtaatacagtcaaattgcagtactctgcgggtgtatacagaatagcagaat

gggcagacattacgaatgcacacggtgtggtgggcccaggtattgttagcggtttgaagcaggc
ggcagaagaagtaacaaaggaacctagaggccttttgatgttagcagaattgtcatgcaagggc
tccctatctactggagaatatactaagggtactgttgacattgcgaagagcgacaaagatttg
ttatcggctttattgctcaaagagacatgggtggaagagatgaaggttacgattggttgattat
gacacccggtgtgggtttagatgacaagggagacgcattgggtcaacagtatagaaccgtggat
gatgtggtctctacaggatctgacattattattgttggaagaggactatttgcaaagggaaggg
atgctaaggtagagggtgaacgttacagaaaagcaggctgggaagcatatttgagaagatgcgg
ccagcaaaactaaaaaactgtattataagtaaatgcatgtatactaaactcacaaattagagct
tcaatttaattatatcagttattaccctatgcggtgtgaaataccgcacagatgcgtaaggaga
aaataccgcatcaggaaattgtaagcgttaatattttgttaaaattcgcgttaaatttttgtta
aatcagctcattttttaaccaataggccgaaatcggcaaaatcccttataaatcaaaagaatag
accgagatagggttgagtgttgttccagtttggaacaagagtccactattaaagaacgtggact
ccaacgtcaaagggcgaaaaaccgtctatcagggcgatggcccactacgtgaaccatcaccta
atcaagttttttggggtcgaggtgccgtaaagcactaaatcggaaccctaaagggagcccccga
tttagagcttgacggggaaagccggcgaacgtggcgagaaaggaagggaagaaagcgaaaggag
cgggcgctagggcgctggcaagtgtagcggtcacgctgcgcgtaaccaccacacccgccgcgct
taatgcgccgctacagggcgcgtccattcgccattcaggctgcgcaactgttgggaagggcgat
cggtgcgggcctcttcgctattacgccagctggcgaaagggggatgtgctgcaaggcgattaag
ttgggtaacgccagggttttcccagtcacgacgttgtaaaacgacggccagtgaattgtaatac
gactcactatagggcgaattggagctccaccgcggtggcggccgctctagaactagtggatcca
aagctgccatttatttgactgctgtgttggaatatttgactgctgaagtgctggagttggctgg
taatgcagccaaggatcttaaagtcaagaggattaccccaagacatctgcaattggccattaga
ggtgacgatgagttagattctttgatcagggctacaattgcttcaggtggtgttttgcctcata
taaataaagcattattattgaaagtggaaaaaaagggaagtaagaaataagaagaatagttttt
ctttcccatttcccgtaattctccctgctcctgtatcctcattctatccatattaacgtctgta
tcattattataagttgaaatttgtctctccgtacccgagttatatacatacatatatgtaatct
atctctcctttcacgtctaatctcttatgtttttactataatatatagcatgaaaatatactct
cacacaatattcccaacgatagcgcagtagccgttatgtacgcagaaatttggtcagggctttc
tttttgttgaactttgcatacccggaaaaatggaaattaaggacacctacaaaagtacactgta
gaaatcaaatcaaagaaaaggtttaatagacgataaatttccatacgtgccgacaggacaaaaa
aaaaaccccccctaccatttctaataacagcttcaccactagttatactgtcgactttctaccga
tttgttggttccttttttataaccagaaaatcatctatcaattgcaatttgtaagagaaggggag
aaagacaaaataataatatggtgccagaaaatagaaggaaacagttggcaattttttgtagttgt
cacatatttgctcacattttattgcgtgtattcagccaccaagacaagcgtttccttttttgcaa
gtaacactgaagctaaatgaaggcttcaatctaatggttttgtcgatattcatcttattaaatt
ctaccttactatggcaactcctaacgaaactattatttggtgaactgaggcttattgagcatga
gcacatttttgaaaggttaccatttaccattataaacaccttgtttatgtcctcactgttccac
gaacggtatttttttcacagtggcatttttttggactattactactctatctgaaagttttccatt
ggattttaaaggataggctggaggccttattacagtcaataaatgattccaccacaatgaaaac
ccttatctttagtagattctcatttaacctcgtactattggcggttgtagactaccagataata
acacgatgcatctcctccatatatacaaaccaaaagagtgatattgaatccacatccctttacc

tgatacaagtaatggagtttaccatgcttttgattgatttgctaaatttattcctacagacttg
tttgaatttctgggaattttatcgctcacaacaaagtctgtctaatgagaacaaccatattgtc
catggcgatcctacagatgaaaacacggttgagtctgatcaatctcagccagtgctgaatgacg
acgacgatgacgacgatgatgatagacaatttaccggcctggagggtaaattcatgtatgaaaa
agcaattgacgtattcacaagattcttaaaaacggcacttcatttgtctatgctaataccattt
aggatgcctatgatgcttttgaaagatgtggtgtgggatatcttggcactatatcaaagtggca
caagtttgtggaaaatctggagaaataacaaacagctcgacgacactcttgtcactgtcaccgt
agaacagctacaaaattctgcaaatgatgacaatatttgtatcatttgtatggatgagttaata
cattctccaaaccagcagacgtggaagaataaaaacaagaaacccaaaaggttaccttgtggcc
acatacttcatttgtcgtgtttaaagaattggatggaacgttctcagacttgtcctatttgtag
attgcctgtctttgatgaaaaaggtaatgttgtgcaaacgactttcacttccaatagtgatatc
acgacacagaccaccgtaacagatagcactgggatagcgacagatcaacaaggtttcgcaaacg
aagtagatctacttcccacaagaacaacttcccctgatataaggatagtgcctactcaaaatat
agacacattagcaatgagaacaaggtcaacctctacaccatctcctacgtggtatacgttccca
ttacataaaactggtgataattctgttgggtcaagccgatcagcctacgaattttgatcacaa
attcagatgagaaagaaaatggtattcctgtcaaattaacaatagaaaatcacgaagtaaattc
tctgcatggagacgggggcgagcaaattgccaagaaaattgtcataccagataaatttatccag
catatcggccgcatcttttacccatacgatgttcctgactatgcgggctatccctatgacgtcc
cggactatgcaggatcctatccatatgacgttccagattacgctgctcagtgcggctagtaatg
acgcatgctttgtccccctgttaatcaggaagtcgcccaaagcgagaatcataccactagacca
cacgcccgatcttttctgagaaccatgaaaagactcagaatttataatacacatcatggccttc
caaagcaactcagcctatgttgaaatgtaattctagtaccatccacttattattggtattcatg
ttactagtatattatcacatacggcataggaaggtgacataatgattaaggaacagctatcaca
tataaaggaagcagaaatcctaaggttgataatgcaataggataataaatgataacatataaaa
tggaagaagaagtaaacattatcataatatgtagatatatcgactctccttttgtgaattcgat
atcaagcttatcgataccgtcgacctcgagggggggcccggtaccagcttttgttccctttagt
gagggttaatttcgagcttggcgtaatcatggtcatagctgtttcctgtgtgaaattgttatcc
gctcacaattccacacaacatacgagccggaagcataaagtgtaaagcctggggtgcctaatga
gtgagctaactcacattaattgcgttgcgctcactgcccgctttccagtcgggaaacctgtcgt
gccagctgcattaatgaatcggccaacgcgcggggagaggcggtttgcgtattgggcgctcttc
cgcttcctcgctcactgactcgctgcgctcggtcgttcggctgcggcgagcggtatcagctcac
tcaaaggcggtaatacggttatccacagaatcaggggataacgcaggaaagaacatgtgagcaa
aaggccagcaaaaggccaggaaccgtaaaaaggccgcgttgctggcgtttttccataggctccg
cccccctgacgagcatcacaaaaatcgacgctcaagtcagaggtggcgaaacccgacaggacta
taaagataccaggcgtttccccctggaagctccctcgtgcgctctcctgttccgaccctgccgc
ttaccggatacctgtccgcctttctcccttcgggaagcgtggcgctttctcatagctcacgctg
taggtatctcagttcggtgtaggtcgttcgctccaagctgggctgtgtgcacgaaccccccgtt
cagcccgaccgctgcgccttatccggtaactatcgtcttgagtccaacccggtaagacacgact
tatcgccactggcagcagccactggtaacaggattagcagagcgaggtatgtaggcggtgctac
agagttcttgaagtggtggcctaactacggctacactagaagaacagtatttggtatctgcgct

ctgctgaagccagttaccttcggaaaaagagttggtagctcttgatccggcaaacaaaccaccg
ctggtagcggtggtttttttgtttgcaagcagcagattacgcgcagaaaaaaaggatctcaa

**pVJ484 (p413-PGPD-CPY-ProtA-K12-13myc)**

ttgagatcctttttttctgcgcgtaatctgctgcttgcaaacaaaaaaaccaccgctaccagcg
gtggtttgtttgccggatcaagagctaccaactctttttccgaaggtaactggcttcagcagag
cgcagataccaaatactgttcttctagtgtagccgtagttaggccaccacttcaagaactctgt
agcaccgcctacatacctcgctctgctaatcctgttaccagtggctgctgccagtggcgataag
tcgtgtcttaccgggttggactcaagacgatagttaccggataaggcgcagcggtcgggctgaa
cggggggttcgtgcacacagcccagcttggagcgaacgacctacaccgaactgagatacctaca
gcgtgagctatgagaaagcgccacgcttcccgaagggagaaaggcggacaggtatccggtaagc
ggcagggtcggaacaggagagcgcacgagggagcttccagggggaaacgcctggtatctttata
gtcctgtcgggtttcgccacctctgacttgagcgtcgatttttgtgatgctcgtcaggggggcg
gagcctatggaaaaacgccagcaacgcggcctttttacggttcctggccttttgctggcctttt
gctcacatgttctttcctgcgttatcccctgattctgtggataaccgtattaccgcctttgagt
gagctgataccgctcgccgcagccgaacgaccgagcgcagcgagtcagtgagcgaggaagcgga
agagcgcccaatacgcaaaccgcctctccccgcgcgttggccgattcattaatgcagctggcac
gacaggtttcccgactggaaagcgggcagtgagcgcaacgcaattaatgtgagttagctcactc
attaggcaccccaggctttacactttatgcttccggctcgtatgttgtgtggaattgtgagcgg
ataacaatttcacacaggaaacagctatgaccatgattacgccaagcgcgcaattaaccctcac
taaagggaacaaaagctggagctcagtttatcattatcaatactcgccatttcaaagaatacgt
aaataattaatagtagtgattttcctaactttatttagtcaaaaaattagccttttaattctgc
tgtaacccgtacatgcccaaaataggggcgggttacacagaatatataacatcgtaggtgtct
gggtgaacagtttattcctggcatccactaaatataatggagcccgctttttaagctggcatcc
agaaaaaaaaagaatcccagcaccaaaatattgttttcttcaccaaccatcagttcataggtcc
attctcttagcgcaactacagagaacaggggcacaaacaggcaaaaaacgggcacaacctcaat
ggagtgatgcaacctgcctggagtaaatgatgacacaaggcaattgacccacgcatgtatctat
ctcattttcttacaccttctattaccttctgctctctctgatttggaaaagctgaaaaaaaag
gttgaaaccagttccctgaaattattcccctacttgactaataagtatataaagacggtaggta
ttgattgtaattctgtaaatctatttcttaaacttcttaaattctacttttatagttagtcttt
tttttagttttaaaacaccagaacttagtttcgacggattctagaactagtATGAAAGCATTCA
CCAGTTTACTATGTGGACTAGGCCTGTCCACTACACTCGCTAAGGCCATCTCATTGCAAAGACC
GTTGGGTCTAGATAAGGACGTTTTGCTGCAAGCTGCGGAAAAATTTGGTTTGGACCTCGACCTG
GATCATCTCTTGAAGGAGTTGGACTCCAATGTATTGGACGCTTGGGCCCAAATAGAGCATTTGT
ACCCAAACCAGGTTATGAGCCTTGAAACTTCCACTAAGCCAAAATTCCCTGAAGCAATCAAAAC
GAAGAAAGACTGGGACTTTGTGGTCAAGAATGACGCAATTGAAAACTATCAGCTTCGTGTCAAC
AAGATTAAGGACCCTAAAATCCTGGGCATTGACCCAAATGTCACACAGTACACGGGTTACTTGG
ATGTGGAAGACGAGGACAAGCATTTCTTCTTTTGGACTTTTGAAAGTAGAAACGATCCTGCAAA
GGATCCGGTCATCCTTTGGTTGAACGGGGGTCCAGGTTGTTCTTCACTAACCGGGCTGTTCTTT
GAATTAGGACCCTCATCCATTGGACCTGATTTGAAACCCATCGGGAACCCTTACTCTTGGAACA
GCAATGCCACCGTGATCTTCCTTGACCAGCCTGTCAACGTTGGGTTCTCGTATTCCGGGTCCTC
AGGTGTTTCCAACACTGTCGCCGCTGGTAAGGATGTCTATAACTTCTTGGAGTTGTTCTTCGAT
CAGTTCCCTGAATACGTCAACAAGGGCCAAGATTTCCACATCGCTGGGGAATCCTACGCCGGCC

ATTACATCCCTGTTTTTGCCTCTGAAATTTTGTCTCACAAGGACAGAAACTTCAACTTAACCTC
CGTCTTGATCGGAAATGGCCTCACTGACCCATTGACTCAGTATAACTATTACGAACCAATGGCC
TGTGGTGAAGGTGGCGAACCATCTGTTTTGCCCTCGGAGGAATGCTCTGCTATGGAAGACTCTT
TGGAACGTTGTTTGGGCTTGATCGAGTCGTGCTATGACTCGCAATCCGTCTGGTCCTGTGTTCC
AGCTACCATTTATTGTAATAACGCCCAATTGGCTCCTTACCAACGTACCGGCAGAAACGTTTAC
GATATCAGGAAGGATTGTGAAGGTGGCAATTTGTGCTACCCAACGTTACAAGATATCGACGACT
ACTTAAACCAGGACTACGTCAAAGAAGCTGTCGGTGCGGAGGTTGACCACTACGAATCCTGTAA
CTTCGATATCAACAGAAATTTCCTGTTTGCGGGTGATTGGATGAAGCCTTACCACACCGCCGTA
ACAGATCTTTTGAATCAAGACCTACCCATTCTGGTATATGCAGGCGATAAAGATTTCATCTGTA
ACTGGTTGGGTAATAAGGCGTGGACGGATGTCTTGCCATGGAAGTACGACGAAGAATTTGCAAG
CCAAAAAGTACGTAACTGGACTGCTTCTATCACCGACGAGGTCGCTGGTGAAGTCAAATCCTAC
AAGCACTTCACCTATTTGAGAGTCTTCAATGGTGGCCACATGGTTCCATTTGACGTCCCTGAAA
ACGCCTTAAGTATGGTTAACGAATGGATCCACGGTGGTTTCTCCTTAACCGGTTCTGGTCTTGC
GCAACACGATGAAGCCGTGGACAACAAATTCAACAAAGAACAACAAAACGCGTTCTATGAGATC
TTACATTTACCTAACTTAAACGAAGAACAACGAAACGCCTTCATCCAAAGTTTAAAAGATGACC
CAAGCCAAAGCGCTAACCTTTTAGCAGAAGCTAAAAAGCTAAATGATGCTCAGGCGCCGAAAGT
AGACAACAAATTCAACAAAGAACAACAAAACGCGTTCTATGAGATCTTACATTTACCTAACTTA
AACGAAGAACAACGAAACGCCTTCATCCAAAGTTTAAAAGATGACCCAAGCCAAAGCGCTAACC
TTTTAGCAGAAGCTAAAAAGCTAAATGATGCTCAGGCGCCGAAAGTAGACGCGCTGggaagcgg
tAAGAAAAAGAAAAAGAAAAAGAAAAAGAAAAAGAAAgGAgccatggGAcggatccccgggtta
attaacgGTGAACAAAAGCTAATCTCCGAGGAAGACTTGAACGGTGAACAAAAATTAATCTCAG
AAGAAGACTTGAACGGACTCGACGGTGAACAAAAGTTGATTTCTGAAGAAGATTTGAACGGTGA
ACAAAAGCTAATCTCCGAGGAAGACTTGAACGGTGAACAAAAATTAATCTCAGAAGAAGACTTG
AACGGACTCGACGGTGAACAAAAGTTGATTTCTGAAGAAGATTTGAACGGTGAACAAAAGCTAA
TCTCCGAGGAAGACTTGAACGGTGAACAAAAATTAATCTCAGAAGAAGACTTGAACGGACTCGA
CGGTGAACAAAAGTTGATTTCTGAAGAAGATTTGAACGGTGAACAAAAGCTAATCTCCGAGGAA
GACTTGAACGGTGAACAAAAATTAATCTCAGAAGAAGACTTGAACGGACTCGACGGTGAACAAA
AGTTGATTTCTGAAGAAGATTTGAACGGTGAACAAAAGCTAATCTCCGAGGAAGACTTGAACGG
TGAACAAAAATTAATCAATCACTAGccgggtagtgacaccgattatttaaagctgcagcatacg
atatatatacatgtgtatatatgtatacctatgaatgtcagtaagtatgtatacgaacagtatg
atactgaagatgacaaggtaatgcatcattctatacgtgtcattctgaacgaggctcgagtcat
gtaattagttatgtcacgcttacattcacgccctccccccacatccgctctaaccgaaaaggaa
ggagttagacaacctgaagtctaggtccctatttatttttttatagttatgttagtattaagaa
cgttatttatatttcaaatttttcttttttttctgtacagacgcgtgtacgcatgtaacattat
actgaaaaccttgcttgagaaggttttgggacgctcgaaggctttaatttgcggccggtaccca
attcgccctatagtgagtcgtattacgcgcgctcactggccgtcgttttacaacgtcgtgactg
ggaaaaccctggcgttacccaacttaatcgccttgcagcacatccccctttcgccagctggcgt
aatagcgaagaggcccgcaccgatcgcccttcccaacagttgcgcagcctgaatggcgaatgga
cgcgccctgtagcggcgcattaagcgcggcgggtgtggtggttacgcgcagcgtgaccgctaca
cttgccagcgccctagcgcccgctcctttcgctttcttcccttcctttctcgccacgttcgccg
gctttccccgtcaagctctaaatcgggggctccctttagggttccgatttagtgctttacggca

cctcgaccccaaaaaacttgattagggtgatggttcacgtagtgggccatcgccctgatagacg
gtttttcgccctttgacgttggagtccacgttctttaatagtggactcttgttccaaactggaa
caacactcaaccctatctcggtctattcttttgatttataagggattttgccgatttcggccta
ttggttaaaaaatgagctgatttaacaaaaatttaacgcgaattttaacaaaatattaacgctt
acaatttcctgatgcggtattttctccttacgcatctgtgcggtatttcacaccgcatagatcc
gtcgagttcaagagaaaaaaaaagaaaaagcaaaaagaaaaaaggaaagcgcgcctcgttcaga
atgacacgtatagaatgatgcattaccttgtcatcttcagtatcatactgttcgtatacatact
tactgacattcataggtatacatatatacacatgtatatatatcgtatgcgctttaataatcg
gtgtcactacataagaacacctttggtggagggaacatcgttggtaccattgggcgaggtggct
tctcttatggcaaccgcaagagccttgaacgcactctcactacggtgatgatcattcttgcctc
gcagacaatcaacgtggagggtaattctgctagcctctgcaaagctttcaagaaaatgcgggat
catctcgcaagagagatctcctactttctccctttgcaaaccaagttcgacaactgcgtacggc
ctgttcgaaagatctaccaccgctctggaaagtgcctcatccaaaggcgcaaatcctgatccaa
accttttactccacgcacggcccctagggcctctttaaaagcttgaccgagagcaatcccgca
gtcttcagtggtgtgatggtcgtctatgtgtaagtcaccaatgcactcaacgattagcgaccag
ccggaatgcttggccagagcatgtatcatatggtccagaaaccctatacctgtgtggacgttaa
tcacttgcgattgtgtggcctgttctgctactgcttctgcctctttttctgggaagatcgagtg
ctctatcgctaggggaccaccctttaaagagatcgcaatctgaatcttggtttcatttgtaata
cgctttactagggctttctgctctgtcatctttgccttcgtttatcttgcctgctcattttta
gtatattcttcgaagaaatcacattactttatataatgtataattcattatgtgataatgccaa
tcgctaagaaaaaaaaagagtcatccgctaggtggaaaaaaaaaaatgaaaatcattaccgagg
cataaaaaaatatagagtgtactagaggaggccaagagtaatagaaaaagaaaattgcgggaaa
ggactgtgttatgacttccctgactaatgccgtgttcaaacgatacctggcagtgactcctagc
gctcaccaagctcttaaaacgggaatttatggtgcactctcagtacaatctgctctgatgccgc
atagttaagccagccccgacacccgccaacacccgctgacgcgccctgacgggcttgtctgctc
ccggcatccgcttacagacaagctgtgaccgtctccgggagctgcatgtgtcagaggttttcac
cgtcatcaccgaaacgcgcgagacgaaagggcctcgtgatacgcctatttttataggttaatgt
catgataataatggtttcttagggtccttttcatcacgtgctataaaaataattataatttaaa
ttttttaatataaatatataaattaaaaatagaaagtaaaaaaagaaattaaagaaaaaatagt
ttttgttttccgaagatgtaaaagactctagggggatcgccaacaaatactaccttttatcttg
ctcttcctgctctcaggtattaatgccgaattgtttcatcttgtctgtgtagaagaccacacac
gaaaatcctgtgattttacattttacttatcgttaatcgaatgtatatctatttaatctgcttt
tcttgtctaataaatatatatgtaaagtacgctttttgttgaaattttttaaacctttgtttat
ttttttttcttcattccgtaactcttctaccttctttatttactttctaaaatccaaatacaaa
acataaaaataaataaacacagagtaaattcccaaattattccatcattaaaagatacgaggcg
cgtgtaagttacaggcaagcgatccgtcaggtggcacttttcggggaaatgtgcgcggaacccc
tatttgtttatttttctaaatacattcaaatatgtatccgctcatgagacaataaccctgataa
atgcttcaataatattgaaaaaggaagagtatgagtattcaacatttccgtgtcgcccttattc
ccttttttgcggcattttgccttcctgtttttgctcacccagaaacgctggtgaaagtaaaaga
tgctgaagatcagttgggtgcacgagtgggttacatcgaactggatctcaacagcggtaagatc
cttgagagttttcgccccgaagaacgttttccaatgatgagcacttttaaagttctgctatgtg

gcgcggtattatcccgtattgacgccgggcaagagcaactcggtcgccgcatacactattctca
gaatgacttggttgagtactcaccagtcacagaaaagcatcttacggatggcatgacagtaaga
gaattatgcagtgctgccataaccatgagtgataacactgcggccaacttacttctgacaacga
tcggaggaccgaaggagctaaccgcttttttgcacaacatgggggatcatgtaactcgccttga
tcgttgggaaccggagctgaatgaagccataccaaacgacgagcgtgacaccacgatgcctgta
gcaatggcaacaacgttgcgcaaactattaactggcgaactacttactctagcttcccggcaac
aattaatagactggatggaggcggataaagttgcaggaccacttctgcgctcggcccttccggc
tggctggtttattgctgataaatctggagccggtgagcgtgggtctcgcggtatcattgcagca
ctggggccagatggtaagccctcccgtatcgtagttatctacacgacggggagtcaggcaacta
tggatgaacgaaatagacagatcgctgagataggtgcctcactgattaagcattggtaactgtc
agaccaagtttactcatatatactttagattgatttaaaacttcatttttaatttaaaaggatc
taggtgaagatcctttttgataatctcatgaccaaaatcccttaacgtgagttttcgttccact
gagcgtcagaccccgtagaaaagatcaaaggatcttc

**pVJ498 (pRS316-hrd1-C399S-3HA)**

gaagatcctttgatcttttctacggggtctgacgctcagtggaacgaaaactcacgttaaggga
ttttggtcatgagattatcaaaaaggatcttcacctagatccttttaaattaaaaatgaagttt
taaatcaatctaaagtatatatgagtaaacttggtctgacagttaccaatgcttaatcagtgag
gcacctatctcagcgatctgtctatttcgttcatccatagttgcctgactccccgtcgtgtaga
taactacgatacgggagggcttaccatctggccccagtgctgcaatgataccgcgagacccacg
ctcaccggctccagatttatcagcaataaaccagccagccggaagggccgagcgcagaagtggt
cctgcaactttatccgcctccatccagtctattaattgttgccgggaagctagagtaagtagtt
cgccagttaatagtttgcgcaacgttgttgccattgctacaggcatcgtggtgtcacgctcgtc
gtttggtatggcttcattcagctccggttcccaacgatcaaggcgagttacatgatcccccatg
ttgtgcaaaaaagcggttagctccttcggtcctccgatcgttgtcagaagtaagttggccgcag
tgttatcactcatggttatggcagcactgcataattctcttactgtcatgccatccgtaagatg
cttttctgtgactggtgagtactcaaccaagtcattctgagaatagtgtatgcggcgaccgagt
tgctcttgcccggcgtcaatacgggataataccgcgccacatagcagaactttaaaagtgctca
tcattggaaaacgttcttcggggcgaaaactctcaaggatcttaccgctgttgagatccagttc
gatgtaacccactcgtgcacccaactgatcttcagcatcttttactttcaccagcgtttctggg
tgagcaaaaacaggaaggcaaaatgccgcaaaaaagggaataagggcgacacggaaatgttgaa
tactcatactcttcctttttcaatattattgaagcatttatcagggttattgtctcatgagcgg
atacatatttgaatgtatttagaaaaataaacaaataggggttccgcgcacatttccccgaaaa
gtgccacctgggtccttttcatcacgtgctataaaaataattataatttaaattttttaatata
aatatataaattaaaaatagaaagtaaaaaaagaaattaaagaaaaaatagtttttgttttccg
aagatgtaaaagactctagggggatcgccaacaaatactaccttttatcttgctcttcctgctc
tcaggtattaatgccgaattgtttcatcttgtctgtgtagaagaccacacacgaaaatcctgtg
attttacattttacttatcgttaatcgaatgtatatctatttaatctgcttttcttgtctaata
aatatatatgtaaagtacgcttttgttgaaattttttaaacctttgtttattttttttttcttc
attccgtaactcttctaccttctttatttactttctaaaatccaaatacaaaacataaaaataa
ataaacacagagtaaattcccaaattattccatcattaaaagatacgaggcgcgtgtaagttac
aggcaagcgatccgtctaagaaaccattattatcatgacattaacctataaaaataggcgtatc
acgaggccctttcgtctcgcgcgtttcggtgatgacggtgaaaacctctgacacatgcagctcc
cggagacggtcacagcttgtctgtaagcggatgccgggagcagacaagcccgtcagggcgcgtc
agcgggtgttggcgggtgtcggggctggcttaactatgcggcatcagagcagattgtactgaga
gtgcaccataccaccttttcaattcatcattttttttttattcttttttttgatttcggtttcc
ttgaaatttttttgattcggtaatctccgaacagaaggaagaacgaaggaaggagcacagactt
agattggtatatatacgcatatgtagtgttgaagaaacatgaaattgcccagtattcttaaccc
aactgcacagaacaaaaacctgcaggaaacgaagataaatcatgtcgaaagctacatataagga
acgtgctgctactcatcctagtcctgttgctgccaagctatttaatatcatgcacgaaaagcaa
acaaacttgtgtgcttcattggatgttcgtaccaccaaggaattactggagttagttgaagcat
taggtcccaaaatttgtttactaaaaacacatgtggatatcttgactgatttttccatggaggg
cacagttaagccgctaaaggcattatccgccaagtacaatttttactcttcgaagacagaaaa
tttgctgacattggtaatacagtcaaattgcagtactctgcgggtgtatacagaatagcagaat

gggcagacattacgaatgcacacggtgtggtgggcccaggtattgttagcggtttgaagcaggc
ggcagaagaagtaacaaaggaacctagaggccttttgatgttagcagaattgtcatgcaagggc
tccctatctactggagaatatactaagggtactgttgacattgcgaagagcgacaaagatttg
ttatcggctttattgctcaaagagacatgggtggaagagatgaaggttacgattggttgattat
gacacccggtgtgggtttagatgacaagggagacgcattgggtcaacagtatagaaccgtggat
gatgtggtctctacaggatctgacattattattgttggaagaggactatttgcaaagggaaggg
atgctaaggtagagggtgaacgttacagaaaagcaggctgggaagcatatttgagaagatgcgg
ccagcaaaactaaaaaactgtattataagtaaatgcatgtatactaaactcacaaattagagct
tcaatttaattatatcagttattaccctatgcggtgtgaaataccgcacagatgcgtaaggaga
aaataccgcatcaggaaattgtaagcgttaatattttgttaaaattcgcgttaaatttttgtta
aatcagctcattttttaaccaataggccgaaatcggcaaaatcccttataaatcaaaagaatag
accgagatagggttgagtgttgttccagtttggaacaagagtccactattaaagaacgtggact
ccaacgtcaaagggcgaaaaaccgtctatcagggcgatggcccactacgtgaaccatcaccct a
atcaagttttttggggtcgaggtgccgtaaagcactaaatcggaaccctaaagggagcccccga
tttagagcttgacggggaaagccggcgaacgtggcgagaaaggaagggaagaaagcgaaaggag
cgggcgctagggcgctggcaagtgtagcggtcacgctgcgcgtaaccaccacacccgccgcgct
taatgcgccgctacagggcgcgtccattcgccattcaggctgcgcaactgttgggaagggcgat
cggtgcgggcctcttcgctattacgccagctggcgaaagggggatgtgctgcaaggcgattaag
ttgggtaacgccagggttttcccagtcacgacgttgtaaaacgacggccagtgaattgtaatac
gactcactatagggcgaattggagctccaccgcggtggcggccgctctagaactagtggatcca
aagctgccatttatttgactgctgtgttggaatatttgactgctgaagtgctggagttggctgg
taatgcagccaaggatcttaaagtcaagaggattaccccaagacatctgcaattggccattaga
ggtgacgatgagttagattctttgatcagggctacaattgcttcaggtggtgttttgcctcata
taaataaagcattattattgaaagtggaaaaaaagggaagtaagaaataagaagaatagttttt
ctttcccatttcccgtaattctccctgctcctgtatcctcattctatccatattaacgtctgta
tcattattataagttgaaatttgtctctccgtacccgagttatatacatacatatatgtaatct
atctctcctttcacgtctaatctcttatgtttttactataatatatagcatgaaaatatactct
cacacaatattcccaacgatagcgcagtagccgttatgtacgcagaaatttggtcagggctttc
tttttgttgaactttgcatacccggaaaaatggaaattaaggacacctacaaaagtacactgta
gaaatcaaatcaaagaaaaggtttaatagacgataaatttccatacgtgccgacaggacaaaaa
aaaaacccccctaccatttctaataacagcttcaccactagttatactgtcgactttctaccga
tttgttggttcctttttataaccagaaaatcatctatcaattgcaatttgtaagagaaggggag
aaagacaaaataataatatggtgccagaaaatagaaggaaacagttggcaatttttgtagttgt
cacatatttgctcacattttattgcgtgtattcagccaccaagacaagcgtttcctttttgcaa
gtaacactgaagctaaatgaaggcttcaatctaatggttttgtcgatattcatcttattaaatt
ctaccttactatggcaactcctaacgaaactattatttggtgaactgaggcttattgagcatga
gcacatttttgaaaggttaccatttaccattataaacaccttgtttatgtcctcactgttccac
gaacggtatttttttcacagtggcatttttttggactattactactctatctgaaagttttccatt
ggattttaaaggataggctggaggccttattacagtcaataaatgattccaccacaatgaaaac
ccttatctttagtagattctcatttaacctcgtactattggcggttgtagactaccagataata
acacgatgcatctcctccatatatacaaaccaaaagagtgatattgaatccacatccctttacc

tgatacaagtaatggagtttaccatgcttttgattgatttgctaaatttattcctacagacttg
tttgaatttctgggaattttatcgctcacaacaaagtctgtctaatgagaacaaccatattgtc
catggcgatcctacagatgaaaacacggttgagtctgatcaatctcagccagtgctgaatgacg
acgacgatgacgacgatgatgatagacaatttaccggcctggagggtaaattcatgtatgaaaa
agcaattgacgtattcacaagattcttaaaaacggcacttcatttgtctatgctaataccattt
aggatgcctatgatgcttttgaaagatgtggtgtgggatatcttggcactatatcaaagtggca
caagtttgtggaaaatctggagaaataacaaacagctcgacgacactcttgtcactgtcaccgt
agaacagctacaaaattctgcaaatgatgacaatatttgtatcatttgtatggatgagttaata
cattctccaaaccagcagacgtggaagaataaaaacaagaaacccaaaaggttaccttgtggcc
acatacttcatttgtcgtgtttaaagaattggatggaacgttctcagacttgtcctatttctag
attgcctgtctttgatgaaaaaggtaatgttgtgcaaacgactttcacttccaatagtgatatc
acgacacagaccaccgtaacagatagcactgggatagcgacagatcaacaaggtttcgcaaacg
aagtagatctacttcccacaagaacaacttcccctgatataaggatagtgcctactcaaaatat
agacacattagcaatgagaacaaggtcaacctctacaccatctcctacgtggtatacgttccca
ttacataaaactggtgataattctgttgggtcaagccgatcagcctacgaattttgatcacaa
attcagatgagaaagaaaatggtattcctgtcaaattaacaatagaaaatcacgaagtaaattc
tctgcatggagacgggggcgagcaaattgccaagaaaattgtcataccagataaatttatccag
catatcggccgcatcttttacccatacgatgttcctgactatgcgggctatccctatgacgtcc
cggactatgcaggatcctatccatatgacgttccagattacgctgctcagtgcggctagtaatg
acgcatgctttgtccccctgttaatcaggaagtcgcccaaagcgagaatcataccactagacca
cacgcccgatcttttctgagaaccatgaaaagactcagaatttataatacacatcatggccttc
caaagcaactcagcctatgttgaaatgtaattctagtaccatccacttattattggtattcatg
ttactagtatattatcacatacggcataggaaggtgacataatgattaaggaacagctatcaca
tataaaggaagcagaaatcctaaggttgataatgcaataggataataaatgataacatataaaa
tggaagaagaagtaaacattatcataatatgtagatatatcgactctccttttgtgaattcgat
atcaagcttatcgataccgtcgacctcgagggggggcccggtaccagcttttgttccctttagt
gagggttaatttcgagcttggcgtaatcatggtcatagctgtttcctgtgtgaaattgttatcc
gctcacaattccacacaacatacgagccggaagcataaagtgtaaagcctggggtgcctaatga
gtgagctaactcacattaattgcgttgcgctcactgcccgctttccagtcgggaaacctgtcgt
gccagctgcattaatgaatcggccaacgcgcggggagaggcggtttgcgtattgggcgctcttc
cgcttcctcgctcactgactcgctgcgctcggtcgttcggctgcggcgagcggtatcagctcac
tcaaaggcggtaatacggttatccacagaatcaggggataacgcaggaaagaacatgtgagcaa
aaggccagcaaaaggccaggaaccgtaaaaaggccgcgttgctggcgtttttccataggctccg
cccccctgacgagcatcacaaaaatcgacgctcaagtcagaggtggcgaaacccgacaggacta
taaagataccaggcgtttccccctggaagctccctcgtgcgctctcctgttccgaccctgccgc
ttaccggatacctgtccgcctttctcccttcgggaagcgtggcgctttctcatagctcacgctg
taggtatctcagttcggtgtaggtcgttcgctccaagctgggctgtgtgcacgaaccccccgtt
cagcccgaccgctgcgccttatccggtaactatcgtcttgagtccaacccggtaagacacgact
tatcgccactggcagcagccactggtaacaggattagcagagcgaggtatgtaggcggtgctac
agagttcttgaagtggtggcctaactacggctacactagaagaacagtatttggtatctgcgct

ctgctgaagccagttaccttcggaaaaagagttggtagctcttgatccggcaaacaaaccaccg
ctggtagcggtggtttttttgtttgcaagcagcagattacgcgcagaaaaaaaggatctcaa

**pVJ556 (p3HA-Ste24)**

ttgagatcctttttttctgcgcgtaatctgctgcttgcaaacaaaaaaaccaccgctaccagcg
gtggtttgtttgccggatcaagagctaccaactctttttccgaaggtaactggcttcagcagag
cgcagataccaaatactgttcttctagtgtagccgtagttaggccaccacttcaagaactctgt
agcaccgcctacatacctcgctctgctaatcctgttaccagtggctgctgccagtggcgataag
tcgtgtcttaccgggttggactcaagacgatagttaccggataaggcgcagcggtcgggctgaa
cggggggttcgtgcacacagcccagcttggagcgaacgacctacaccgaactgagatacctaca
gcgtgagctatgagaaagcgccacgcttcccgaagggagaaaggcggacaggtatccggtaagc
ggcagggtcggaacaggagagcgcacgagggagcttccagggggaaacgcctggtatctttata
gtcctgtcgggtttcgccacctctgacttgagcgtcgatttttgtgatgctcgtcaggggggcg
gagcctatggaaaaacgccagcaacgcggcctttttacggttcctggccttttgctggcctttt
gctcacatgttctttcctgcgttatcccctgattctgtggataaccgtattaccgcctttgagt
gagctgataccgctcgccgcagccgaacgaccgagcgcagcgagtcagtgagcgaggaagcgga
agagcgcccaatacgcaaaccgcctctccccgcgcgttggccgattcattaatgcagctggcac
gacaggtttcccgactggaaagcgggcagtgagcgcaacgcaattaatgtgagttagctcactc
attaggcaccccaggctttacactttatgcttccggctcgtatgttgtgtggaattgtgagcgg
ataacaatttcacacaggaaacagctatgaccatgattacgccaagctcgaaattaaccctcac
taaagggaacaaaagctggtaccgggccccccctcgagttcacgggccacgccattggctccgt
atatgtgatgctgttgtccttaagaccgttctgccaaccctggatcggcaggttccttatctac
gaggcctccactccctttgtgaacatcaattggttcatcatgcaatgtaacgccaagagcaaga
actctatccctctgtggtttaatgtcgtcaatggccttttgctcatgactgtgtttttcgtcgt
caggatttgctggggctccatcgcatccgctctcttattcaggcagatgtggaaggtaagagac
gagttacccaagttttctgctgtcacgatgatgtcgctgaacattttcatgaacctttgaacg
tgctttggttcaagaagatgattagaatcgccaagaaacttgcaaaaccagccccaacgtcgaa
gctagactaaacctacctttttttctatcttcaacaacgaaacgccttacacacacacacacat
acatctacatacatacatacaaatatacatatatgtaaacttgtatattcattcctattaacca
aaaagaggcaattaaactttccctctttttctacgtcatttactcaaaaactctaattccttc
gtctctgttctgccattttctccagaaaaaaatcgacgggaaataaaaaaaaaaagacaacgaa
caagagaaaaagttcgcgaattataaaccacttctataattaacaggaaaaggaaggaaaaaaa
aggaggaaatagaaaactgcaggcctttattcatgagcggccgcatcttttacccatacgatgt
tcctgactatgcgggctatccctatgacgtcccggactatgcaggatcctatccatatgacgtt
ccagattacgctgctcagtgcggccgctttgatcttaagacgattctcgaccatcctaatatcc
cgtggaaattaatcatttctgggttctcgattgcccaattttctttcgaatcttacttgacgta
cagacagtaccagaagctatctgaaacaaagttgccacctgtgctggaagacgaaattgatgat
gaaacttttcataaatcaaggaactactcccgggccaaggccaagttctccattttcggtgacg
tctataacctagcccaaaagctagtttcatcaaatacgacctcttccctaaaatctggcacat
ggccgtttctttattgaatgcagtcctgccagtcagatttcatatggtctccactgtcgcacag
agtttatgcttcttgggtctcttatccagtttgtctaccttggttgatttgccactctcttact
atagccattttgtcctggaagaaaaatttggtttcaataaattgaccgtccaactatggatcac
cgatatgatcaagagtctgactttggcgtatgctattggtggcccaatcctttacctgttcctt

aagatctttgataaattccctactgatttcctttggtacattatggtcttcttgttcgttgtcc
aaatcttagccatgacaatcattccagtcttcatcatgcccatgtttaataagttcactccatt
ggaggacggtgaactgaaaaaatctattgaaagtttggccgatagagttgggttccctctagat
aagatttttgtcattgacggctcaaaaagatcttctcattcaaacgcatatttcacaggtttgc
cattcacctccaagagaattgttttgttcgacactttagtgaacagtaattctactgatgaaat
tacggctgttttggcccatgaaatcggtcactggcaaaaaaaccacatcgttaatatggtcatc
tttagtcaattgcacaccttcctcatttctccctttcaccagcatctacagaaatacatcat
tttacaacaccttcggctttttcttagagaagtccactggcagttttgttgatcccgttatcac
taaggaattccccattatcattggatttatgttatttaacgacttattaactccactcgaatgt
gccatgcaattcgtgatgagtttaatttccagaactcatgaatatcaagctgatgcttatgcta
aaaaattgggctacaagcaaaatctatgtagggctctaattgatctacaaatcaaaaaccttc
caccatgaatgtagatcctctgtattctagctatcattattcccatccaactctagctgaaaga
ttgaccgctctagactatgttagtgaaaagaagaaaaactaatctatagagtacacatattagc
atgtaccgttaaattcagcttcgttatgtctatatctacatacatacacaggtatctactataa
gaataaaggaaagaaaaaataaacgattaaacatttatttttttgcatcgctatcatcattac
ctttctcattgttatcatcattatctttcccatcttcatcgtcattttcttcatcattgattat
ttcaatgatttctgtggtttcagtcttcacggtaaatggttggggttcttcaatatcctcatca
ttaaccatgaattggttcaatcttttgaactcttcatttttctcctccctaatggcattataca
aaagaaaatttatccaaacttctgcaaaacagtaaataaaaacagagttattgtgaacgtaaaa
gtttgactcccaaaggtacgagatacttgtcaaaataaaaaaaattagtaatctaactggcaca
acagattggaagtatttaacattgttttctaacaggggaattagatcatgaacaccgttaagag
taaacagaagagcgaaaagacccaattgaccactataacgtgacagtgttccactagttctaga
gcggccggccgccaccgcggtggagctccaattcgccctatagtgagtcgtattacaattcact
ggccgtcgttttacaacgtcgtgactgggaaaaccctggcgttacccaacttaatcgccttgca
gcacatccccctttcgccagctggcgtaatagcgaagaggcccgcaccgatcgcccttcccaac
agttgcgcagcctgaatggcgaatggacgcgccctgtagcggcgcattaagcgcggcgggtgtg
gtggttacgcgcagcgtgaccgctacacttgccagcgccctagcgcccgctcctttcgctttct
tcccttcctttctcgccacgttcgccggctttccccgtcaagctctaaatcgggggctcccttt
agggttccgatttagtgctttacggcacctcgaccccaaaaaacttgattagggtgatggttca
cgtagtgggccatcgccctgatagacggtttttcgccctttgacgttggagtccacgttcttta
atagtggactcttgttccaaactggaacaacactcaaccctatctcggtctattcttttgattt
ataagggattttgccgatttcggcctattggttaaaaaatgagctgatttaacaaaaatttaac
gcgaattttaacaaaatattaacgcttacaatttcctgatgcggtattttctccttacgcatct
gtgcggtatttcacaccgcatagggtaataactgatataattaaattgaagctctaatttgtga
gtttagtatacatgcatttacttataatacagtttttttagttttgctggccgcatcttctcaaa
tatgcttcccagcctgcttttctgtaacgttcaccctctaccttagcatcccttccctttgcaa
atagtcctcttccaacaataataatgtcagatcctgtagagaccacatcatccacggttctata
ctgttgacccaatgcgtctcccttgtcatctaaacccacaccgggtgtcataatcaaccaatcg
taaccttcatctcttccacccatgtctctttgagcaataaagccgataacaaaatctttgtcgc
tcttcgcaatgtcaacagtacccttagtatattctccagtagatagggagcccttgcatgacaa
ttctgctaacatcaaaaggcctctaggttcctttgttacttcttctgccgcctgcttcaaaccg

ctaacaatacctgggcccaccacaccgtgtgcattcgtaatgtctgcccattctgctattctgt
atacacccgcagagtactgcaatttgactgtattaccaatgtcagcaaattttctgtcttcgaa
gagtaaaaaattgtacttggcggataatgcctttagcggcttaactgtgccctccatggaaaaa
tcagtcaagatatccacatgtgtttttagtaaacaaattttgggacctaatgcttcaactaact
ccagtaattccttggtggtacgaacatccaatgaagcacacaagtttgtttgcttttcgtgcat
gatattaaatagcttggcagcaacaggactaggatgagtagcagcacgttccttatatgtagct
ttcgacatgatttatcttcgtttcctgcaggtttttgttctgtgcagttgggttaagaatactg
ggcaatttcatgtttcttcaacactacatatgcgtatatataccaatctaagtctgtgctcctt
ccttcgttcttccttctgttcggagattaccgaatcaaaaaaatttcaaggaaaccgaaatcaa
aaaaaagaataaaaaaaaaatgatgaattgaaaaggtggtatggtgcactctcagtacaatctg
ctctgatgccgcatagttaagccagccccgacacccgccaacacccgctgacgcgccctgacgg
gcttgtctgctcccggcatccgcttacagacaagctgtgaccgtctccgggagctgcatgtgtc
agaggttttcaccgtcatcaccgaaacgcgcgagacgaaagggcctcgtgatacgcctattttt
ataggttaatgtcatgataataatggtttcttagacggatcgcttgcctgtaacttacacgcgc
ctcgtatcttttaatgatggaataatttgggaatttactctgtgtttatttatttttatgtttt
gtatttggattttagaaagtaaataaagaaggtagaagagttacggaatgaagaaaaaaaaata
aacaaaggtttaaaaaatttcaacaaaaagcgtactttacatatatatttattagacaagaaaa
gcagattaaatagatatacattcgattaacgataagtaaaatgtaaaatcacaggattttcgtg
tgtggtcttctacacagacaagatgaaacaattcggcattaatacctgagagcaggaagagcaa
gataaaaggtagtatttgttggcgatccccctagagtcttttacatcttcggaaaacaaaaact
atttttctttaatttctttttttactttctatttttaatttatatatttatattaaaaaattt
aaattataattatttttatagcacgtgatgaaaaggacccaggtggcacttttcggggaaatgt
gcgcggaacccctatttgtttatttttctaaatacattcaaatatgtatccgctcatgagacaa
taaccctgataaatgcttcaataatattgaaaaaggaagagtatgagtattcaacatttccgtg
tcgcccttattcccttttttgcggcattttgccttcctgtttttgctcacccagaaacgctggt
gaaagtaaaagatgctgaagatcagttgggtgcacgagtgggttacatcgaactggatctcaac
agcggtaagatccttgagagttttcgccccgaagaacgttttccaatgatgagcacttttaaag
ttctgctatgtggcgcggtattatcccgtattgacgccgggcaagagcaactcggtcgccgcat
acactattctcagaatgacttggttgagtactcaccagtcacagaaaagcatcttacggatggc
atgacagtaagagaattatgcagtgctgccataaccatgagtgataacactgcggccaacttac
ttctgacaacgatcggaggaccgaaggagctaaccgcttttttgcacaacatgggggatcatgt
aactcgccttgatcgttgggaaccggagctgaatgaagccataccaaacgacgagcgtgacacc
acgatgcctgtagcaatggcaacaacgttgcgcaaactattaactggcgaactacttactctag
cttcccggcaacaattaatagactggatggaggcggataaagttgcaggaccacttctgcgctc
ggcccttccggctggctggtttattgctgataaatctggagccggtgagcgtgggtctcgcggt
atcattgcagcactggggccagatggtaagccctcccgtatcgtagttatctacacgacgggga
gtcaggcaactatggatgaacgaaatagacagatcgctgagataggtgcctcactgattaagca
ttggtaactgtcagaccaagtttactcatatatactttagattgatttaaaacttcatttttaa
tttaaaaggatctaggtgaagatcctttttgataatctcatgaccaaaatcccttaacgtgagt
tttcgttccactgagcgtcagaccccgtagaaaagatcaaaggatcttc

**pVJ557 (p3HA-ste24-H297A)**

ttgagatcctttttttctgcgcgtaatctgctgcttgcaaacaaaaaaaccaccgctaccagcg
gtggtttgtttgccggatcaagagctaccaactctttttccgaaggtaactggcttcagcagag
cgcagataccaaatactgttcttctagtgtagccgtagttaggccaccacttcaagaactctgt
agcaccgcctacatacctcgctctgctaatcctgttaccagtggctgctgccagtggcgataag
tcgtgtcttaccgggttggactcaagacgatagttaccggataaggcgcagcggtcgggctgaa
cggggggttcgtgcacacagcccagcttggagcgaacgacctacaccgaactgagatacctaca
gcgtgagctatgagaaagcgccacgcttcccgaagggagaaaggcggacaggtatccggtaagc
ggcagggtcggaacaggagagcgcacgagggagcttccagggggaaacgcctggtatctttata
gtcctgtcgggtttcgccacctctgacttgagcgtcgatttttgtgatgctcgtcaggggggcg
gagcctatggaaaaacgccagcaacgcggcctttttacggttcctggccttttgctggcctttt
gctcacatgttctttcctgcgttatcccctgattctgtggataaccgtattaccgcctttgagt
gagctgataccgctcgccgcagccgaacgaccgagcgcagcgagtcagtgagcgaggaagcgga
agagcgcccaatacgcaaaccgcctctccccgcgcgttggccgattcattaatgcagctggcac
gacaggtttcccgactggaaagcgggcagtgagcgcaacgcaattaatgtgagttagctcactc
attaggcaccccaggctttacactttatgcttccggctcgtatgttgtgtggaattgtgagcgg
ataacaatttcacacaggaaacagctatgaccatgattacgccaagctcgaaattaaccctcac
taaagggaacaaaagctggtaccgggccccccctcgagttcacgggccacgccattggctccgt
atatgtgatgctgttgtccttaagaccgttctgccaaccctggatcggcaggttccttatctac
gaggcctccactccctttgtgaacatcaattggttcatcatgcaatgtaacgccaagagcaaga
actctatccctctgtggtttaatgtcgtcaatggccttttgctcatgactgtgtttttcgtcgt
caggatttgctggggctccatcgcatccgctctcttattcaggcagatgtggaaggtaagagac
gagttacccaagttttctgctgtcacgatgatgtcgctgaacattttcatgaacctttttgaacg
tgctttggttcaagaagatgattagaatcgccaagaaacttgcaaaaccagccccaacgtcgaa
gctagactaaacctacctttttttctatcttcaacaacgaaacgccttacacacacacacacat
acatctacatacatacatacaaatatacatatatgtaaacttgtatattcattcctattaacca
aaaagaggcaattaaacttttccctctttttctacgtcatttactcaaaaactctaattccttc
gtctctgttctgccattttctccagaaaaaaatcgacgggaaataaaaaaaaaagacaacgaac
aagagaaaaagttcgcgaattataaaccacttctataattaacaggaaaaggaaggaaaaaaa
ggaggaaatagaaaactgcaggcctttattcatgagcggccgcatcttttacccatacgatgtt
cctgactatgcgggctatccctatgacgtcccggactatgcaggatcctatccatatgacgttc
cagattacgctgctcagtgcggccgctttgatcttaagacgattctcgaccatcctaatatccc
gtggaaattaatcatttctgggttctcgattgcccaattttctttcgaatcttacttgacgtac
agacagtaccagaagctatctgaaacaaagttgccacctgtgctggaagacgaaattgatgatg
aaacttttcataaatcaaggaactactcccgggccaaggccaagttctccattttcggtgacgt
ctataacctagcccaaaagctagttttcatcaaatacgacctcttccctaaaatctggcacatg
gccgtttctttattgaatgcagtcctgccagtcagatttcatatggtctccactgtcgcacaga
gtttatgcttcttgggtctcttatccagtttgtctaccttggttgatttgccactctcttacta
tagccattttgtcctggaagaaaaatttggtttcaataaattgaccgtccaactatggatcacc
gatatgatcaagagtctgactttggcgtatgctattggtggcccaatcctttacctgttcctta

agatctttgataaattccctactgatttcctttggtacattatggtcttcttgttcgttgtcca
aatcttagccatgacaatcattccagtcttcatcatgcccatgtttaataagttcactccattg
gaggacggtgaactgaaaaaatctattgaaagtttggccgatagagttgggttccctctagata
agatttttgtcattgacggctcaaaaagatcttctcattcaaacgcatatttcacaggtttgcc
attcacctccaagagaattgttttgttcgacactttagtgaacagtaattctactgatgaaatt
acggctgttttggcagctgaaatcggtcactggcaaaaaaaccacatcgttaatatggtcatct
ttagtcaattgcacaccttcctcatttctcccttttcaccagcatctacagaaatacatcatt
ttacaacaccttcggctttttcttagagaagtccactggcagttttgttgatcccgttatcact
aaggaattccccattatcattggatttatgttatttaacgacttattaactccactcgaatgtg
ccatgcaattcgtgatgagtttaatttccagaactcatgaatatcaagctgatgcttatgctaa
aaaattgggctacaagcaaaatctatgtagggctctaattgatctacaaatcaaaaacctttcc
accatgaatgtagatcctctgtattctagctatcattattcccatccaactctagctgaaagat
tgaccgctctagactatgttagtgaaaagaagaaaaactaatctatagagtacacatattagca
tgtaccgttaaattcagcttcgttatgtctatatctacatacatacacaggtatctactataag
aataaaggaaagaaaaaataaacgattaaacattttatttttttgcatcgctatcatcattacc
tttctcattgttatcatcattatctttcccatcttcatcgtcattttcttcatcattgattatt
tcaatgatttctgtggtttcagtcttcacggtaaatggttggggttcttcaatatcctcatcat
taaccatgaattggttcaatcttttgaactcttcatttttctcctccctaatggcattatacaa
aagaaaatttatccaaacttctgcaaaacagtaaataaaaacagagttattgtgaacgtaaaag
tttgactcccaaaggtacgagatacttgtcaaaataaaaaaaattagtaatctaactggcacaa
cagattggaagtatttaacattgttttctaacaggggaattagatcatgaacaccgttaagagt
aaacagaagagcgaaaagacccaattgaccactataacgtgacagtgttccactagttctagag
cggccggccgccaccgcggtggagctccaattcgccctatagtgagtcgtattacaattcactg
gccgtcgttttacaacgtcgtgactgggaaaaccctggcgttacccaacttaatcgccttgcag
cacatccccctttcgccagctggcgtaatagcgaagaggcccgcaccgatcgcccttcccaaca
gttgcgcagcctgaatggcgaatggacgcgccctgtagcggcgcattaagcgcggcgggtgtgg
tggttacgcgcagcgtgaccgctacacttgccagcgccctagcgcccgctcctttcgctttctt
cccttcctttctcgccacgttcgccggctttccccgtcaagctctaaatcgggggctcccttta
gggttccgatttagtgctttacggcacctcgaccccaaaaaacttgattagggtgatggttcac
gtagtgggccatcgccctgatagacggtttttcgccctttgacgttggagtccacgttctttaa
tagtggactcttgttccaaactggaacaacactcaaccctatctcggtctattcttttgattta
taagggattttgccgatttcggcctattggttaaaaaatgagctgatttaacaaaaatttaacg
cgaattttaacaaaatattaacgcttacaatttcctgatgcggtattttctccttacgcatctg
tgcggtatttcacaccgcatagggtaataactgatataattaaattgaagctctaatttgtgag
tttagtatacatgcatttacttataatacagtttttagttttgctggccgcatcttctcaaat
atgcttcccagcctgcttttctgtaacgttcaccctctaccttagcatcccttccctttgcaaa
tagtcctcttccaacaataataatgtcagatcctgtagagaccacatcatccacggttctatac
tgttgacccaatgcgtctcccttgtcatctaaacccacaccgggtgtcataatcaaccaatcgt
aaccttcatctcttccacccatgtctctttgagcaataaagccgataacaaaatctttgtcgct
cttcgcaatgtcaacagtacccttagtatattctccagtagatagggagcccttgcatgacaat
tctgctaacatcaaaaggcctctaggttcctttgttacttcttctgccgcctgcttcaaaccgc

taacaatacctgggcccaccacaccgtgtgcattcgtaatgtctgcccattctgctattctgta
tacacccgcagagtactgcaatttgactgtattaccaatgtcagcaaattttctgtcttcgaag
agtaaaaaattgtacttggcggataatgcctttagcggcttaactgtgccctccatggaaaaat
cagtcaagatatccacatgtgtttttagtaaacaaattttggacctaatgcttcaactaactc
cagtaattccttggtggtacgaacatccaatgaagcacacaagtttgtttgcttttcgtgcatg
atattaaatagcttggcagcaacaggactaggatgagtagcagcacgttccttatatgtagctt
tcgacatgatttatcttcgtttcctgcaggtttttgttctgtgcagttgggttaagaatactgg
gcaatttcatgtttcttcaacactacatatgcgtatatataccaatctaagtctgtgctccttc
cttcgttcttccttctgttcggagattaccgaatcaaaaaaatttcaaggaaaccgaaatcaaa
aaaaagaataaaaaaaaaatgatgaattgaaaaggtggtatggtgcactctcagtacaatctgc
tctgatgccgcatagttaagccagccccgacacccgccaacacccgctgacgcgccctgacggg
cttgtctgctcccggcatccgcttacagacaagctgtgaccgtctccgggagctgcatgtgtca
gaggttttcaccgtcatcaccgaaacgcgcgagacgaaagggcctcgtgatacgcctattttta
taggttaatgtcatgataataatggtttcttagacggatcgcttgcctgtaacttacacgcgcc
tcgtatcttttaatgatggaataatttgggaatttactctgtgtttatttatttttatgttttg
tatttggattttagaaagtaaataaagaaggtagaagagttacggaatgaagaaaaaaaataa
acaaaggtttaaaaaatttcaacaaaaagcgtactttacatatatatttattagacaagaaaag
cagattaaatagatatacattcgattaacgataagtaaaatgtaaaatcacaggattttcgtgt
gtggtcttctacacagacaagatgaaacaattcggcattaatacctgagagcaggaagagcaag
ataaaaggtagtatttgttggcgatccccctagagtcttttacatcttcggaaaacaaaaacta
ttttttctttaatttcttttttttactttctatttttaatttatatatttatattaaaaattta
aattataattatttttatagcacgtgatgaaaaggacccaggtggcacttttcggggaaatgtg
cgcggaacccctatttgtttatttttctaaatacattcaaatatgtatccgctcatgagacaat
aaccctgataaatgcttcaataatattgaaaaaggaagagtatgagtattcaacatttccgtgt
cgcccttattcccttttttgcggcattttgccttcctgtttttgctcacccagaaacgctggtg
aaagtaaaagatgctgaagatcagttgggtgcacgagtgggttacatcgaactggatctcaaca
gcggtaagatccttgagagttttcgccccgaagaacgttttccaatgatgagcacttttaaagt
tctgctatgtggcgcggtattatcccgtattgacgccgggcaagagcaactcggtcgccgcata
cactattctcagaatgacttggttgagtactcaccagtcacagaaaagcatcttacggatggca
tgacagtaagagaattatgcagtgctgccataaccatgagtgataacactgcggccaacttact
tctgacaacgatcggaggaccgaaggagctaaccgcttttttgcacaacatgggggatcatgta
actcgccttgatcgttgggaaccggagctgaatgaagccataccaaacgacgagcgtgacacca
cgatgcctgtagcaatggcaacaacgttgcgcaaactattaactggcgaactacttactctagc
ttcccggcaacaattaatagactggatggaggcggataaagttgcaggaccacttctgcgctcg
gcccttccggctggctggtttattgctgataaatctggagccggtgagcgtgggtctcgcggta
tcattgcagcactggggccagatggtaagccctcccgtatcgtagttatctacacgacggggag
tcaggcaactatggatgaacgaaatagacagatcgctgagataggtgcctcactgattaagcat
tggtaactgtcagaccaagtttactcatatatactttagattgatttaaaacttcatttttaat
ttaaaaggatctaggtgaagatcctttttgataatctcatgaccaaaatcccttaacgtgagtt
ttcgttccactgagcgtcagaccccgtagaaaagatcaaaggatcttc

**pVJ636 (p416-PGAL1-Kar2SS-6xIAPP-3HA)**

agatcagttgggtgcacgagtgggttacatcgaactggatctcaacagcggtaagatccttgag
agttttcgccccgaagaacgttttccaatgatgagcacttttaaagttctgctatgtggcgcgg
tattatcccgtattgacgccgggcaagagcaactcggtcgccgcatacactattctcagaatga
cttggttgagtactcaccagtcacagaaaagcatcttacggatggcatgacagtaagagaatta
tgcagtgctgccataaccatgagtgataacactgcggccaacttacttctgacaacgatcggag
gaccgaaggagctaaccgcttttttgcacaacatgggggatcatgtaactcgccttgatcgttg
ggaaccggagctgaatgaagccataccaaacgacgagcgtgacaccacgatgcctgtagcaatg
gcaacaacgttgcgcaaactattaactggcgaactacttactctagcttcccggcaacaattaa
tagactggatggaggcggataaagttgcaggaccacttctgcgctcggcccttccggctggctg
gtttattgctgataaatctggagccggtgagcgtgggtctcgcggtatcattgcagcactgggg
ccagatggtaagccctcccgtatcgtagttatctacacgacggggagtcaggcaactatggatg
aacgaaatagacagatcgctgagataggtgcctcactgattaagcattggtaactgtcagacca
agtttactcatatatactttagattgatttaaaacttcatttttaatttaaaaggatctaggtg
aagatcctttttgataatctcatgaccaaaatcccttaacgtgagttttcgttccactgagcgt
cagaccccgtagaaaagatcaaaggatcttcttgagatcctttttttctgcgcgtaatctgctg
cttgcaaacaaaaaaaccaccgctaccagcggtggtttgtttgccggatcaagagctaccaact
ctttttccgaaggtaactggcttcagcagagcgcagataccaaatactgttcttctagtgtagc
cgtagttaggccaccacttcaagaactctgtagcaccgcctacatacctcgctctgctaatcct
gttaccagtggctgctgccagtggcgataagtcgtgtcttaccgggttggactcaagacgatag
ttaccggataaggcgcagcggtcgggctgaacggggggttcgtgcacacagcccagcttggagc
gaacgacctacaccgaactgagatacctacagcgtgagctatgagaaagcgccacgcttcccga
agggagaaaggcggacaggtatccggtaagcggcagggtcggaacaggagagcgcacgagggag
cttccagggggaaacgcctggtatctttatagtcctgtcgggtttcgccacctctgacttgagc
gtcgatttttgtgatgctcgtcaggggggcggagcctatggaaaaacgccagcaacgcggcctt
tttacggttcctggccttttgctggccttttgctcacatgttctttcctgcgttatcccctgat
tctgtggataaccgtattaccgcctttgagtgagctgataccgctcgccgcagccgaacgaccg
agcgcagcgagtcagtgagcgaggaagcggaagagcgcccaatacgcaaaccgcctctccccgc
gcgttggccgattcattaatgcagctggcacgacaggtttcccgactggaaagcgggcagtgag
cgcaacgcaattaatgtgagttagctcactcattaggcaccccaggctttacactttatgcttc
cggctcgtatgttgtgtggaattgtgagcggataacaatttcacacaggaaacagctatgacca
tgattacgccaagcgcgcaattaaccctcactaaagggaacaaaagctggagctctagtacgga
ttagaagccgccgagcgggtgacagccctccgaaggaagactctcctccgtgcgtcctcgtctt
caccggtcgcgttcctgaaacgcagatgtgcctcgcgccgcactgctccgaacaataaagattc
tacaatactagcttttatggttatgaagaggaaaaattggcagtaacctggccccacaaacctt
caaatgaacgaatcaaattaacaaccataggatgataatgcgattagttttttagccttatttc
tggggtaattaatcagcgaagcgatgatttttgatctattaacagatatataaatgcaaaaact
gcataaccactttaactaatactttcaacattttcggtttgtattacttcttattcaaatgtaa
taaaagtatcaacaaaaaattgttaatatacctctatactttaacgtcaaggagaaaaaacccc
ggattctagaaaaacaactaattcgaaatgtttttcaacagactaagcgctggcaagctgctgg

taccactctccgtggtcctgtacgccctttcgtggtaatattacctttacagaattctttcca
ctcctccaatgttttagttagaggtaaatgcaacactgccacatgtgctactcaaagattggct
aactttttggtccactcctctaacaatttcggtgccattttgtcctctaccaatgttggttcta
acacctacaaaagagaagccgaagctaagtgtaataccgcaacatgtgcaacacaaagattagc
aaatttcttagttcattcttccaacaactttggtgctatcttgtctagtacaaacgtcggttct
aatacttataagagagaagcagacgctgaagctaaatgcaatactgccacatgcgccacccaaa
gattggccaacttcttagtacattccagtaacaacttcggtgcaatattgagttctaccaacgt
cggtagtaacacatacaagagagaagctgacgccgaagcaaagtgtaacacagctacatgcgcc
actcaaagattagcaaactttttggttcacagttcaaacaattttggtgcaatcttatcttcca
ccaatgtcggttccaacacttacaagagagaagccgacgctgaagccaagtgtaatactgcaac
ctgcgctactcaaagattggcaaatttcttggtacacagtagtaacaactttggtgccatttta
tcctcaactaacgtcggttcaaatacctacaaaagagaagctgacgctgaagccaaatgcaaca
cagcaacctgtgccacccaaagattggctaactttttagttcacagtagtaacaatttcggtgc
tatcttgagtagtactaatgttggtagtaatacctacggccgcatcttttacccatacgatgtt
cctgactatgcgggctatccctatgacgtcccggactatgcaggatcctatccatatgacgttc
cagattacgctgctcagtgcggctagaagcttatcgataccgtcgacctcgagtcatgtaatta
gttatgtcacgcttacattcacgccctccccccacatccgctctaaccgaaaaggaaggagtta
gacaacctgaagtctaggtccctatttattttttttatagttatgttagtattaagaacgttatt
tatatttcaaatttttcttttttttctgtacagacgcgtgtacgcatgtaacattatactgaaa
accttgcttgagaaggttttgggacgctcgaaggctttaatttgcggccggtacccaattcgcc
ctatagtgagtcgtattacgcgcgctcactggccgtcgttttacaacgtcgtgactgggaaaac
cctggcgttacccaacttaatcgccttgcagcacatccccctttcgccagctggcgtaatagcg
aagaggcccgcaccgatcgcccttcccaacagttgcgcagcctgaatggcgaatggacgcgccc
tgtagcggcgcattaagcgcggcgggtgtggtggttacgcgcagcgtgaccgctacacttgcca
gcgccctagcgcccgctcctttcgctttcttcccttcctttctcgccacgttcgccggctttcc
ccgtcaagctctaaatcgggggctccctttagggttccgatttagtgctttacggcacctcgac
cccaaaaaacttgattagggtgatggttcacgtagtgggccatcgccctgatagacggtttttc
gccctttgacgttggagtccacgttctttaatagtggactcttgttccaaactggaacaacact
caaccctatctcggtctattcttttgatttataagggattttgccgatttcggcctattggtta
aaaaatgagctgatttaacaaaaatttaacgcgaattttaacaaaatattaacgcttacaattt
cctgatgcggtattttctccttacgcatctgtgcggtatttcacaccgcatagggtaataactg
atataattaaattgaagctctaatttgtgagtttagtatacatgcatttacttataatacagtt
ttttagttttgctggccgcatcttctcaaatatgcttcccagcctgcttttctgtaacgttcac
cctctaccttagcatcccttccctttgcaaatagtcctcttccaacaataataatgtcagatcc
tgtagagaccacatcatccacggttctatactgttgacccaatgcgtctcccttgtcatctaaa
cccacaccgggtgtcataatcaaccaatcgtaaccttcatctcttccacccatgtctctttgag
caataaagccgataacaaaatctttgtcgctcttcgcaatgtcaacagtacccttagtatattc
tccagtagatagggagcccttgcatgacaattctgctaacatcaaaaggcctctaggttccttt
gttacttcttctgccgcctgcttcaaaccgctaacaatacctgggcccaccacaccgtgtgcat
tcgtaatgtctgcccattctgctattctgtatacacccgcagagtactgcaatttgactgtatt
accaatgtcagcaaattttctgtcttcgaagagtaaaaaattgtacttggcggataatgccttt

agcggcttaactgtgccctccatggaaaaatcagtcaagatatccacatgtgtttttagtaaac
aaattttgggacctaatgcttcaactaactccagtaattccttggtggtacgaacatccaatga
agcacacaagtttgtttgcttttcgtgcatgatattaaatagcttggcagcaacaggactagga
tgagtagcagcacgttccttatatgtagctttcgacatgatttatcttcgtttcctgcaggttt
ttgttctgtgcagttgggttaagaatactgggcaatttcatgtttcttcaacactacatatgcg
tatatataccaatctaagtctgtgctccttccttcgttcttccttctgttcggagattaccgaa
tcaaaaaaatttcaaggaaaccgaaatcaaaaaaaagaataaaaaaaaaatgatgaattgaaaa
ggtggtatggtgcactctcagtacaatctgctctgatgccgcatagttaagccagccccgacac
ccgccaacacccgctgacgcgccctgacgggcttgtctgctcccggcatccgcttacagacaag
ctgtgaccgtctccgggagctgcatgtgtcagaggttttcaccgtcatcaccgaaacgcgcgag
acgaaagggcctcgtgatacgcctatttttataggttaatgtcatgataataatggtttcttag
acggatcgcttgcctgtaacttacacgcgcctcgtatcttttaatgatggaataatttgggaat
ttactctgtgtttatttatttttatgttttgtatttggattttagaaagtaaataaagaaggta
gaagagttacggaatgaagaaaaaaaaaataaacaaaggtttaaaaaatttcaacaaaaagcgta
ctttacatatatatttattagacaagaaaagcagattaaatagatatacattcgattaacgata
agtaaaatgtaaaatcacaggattttcgtgtgtggtcttctacacagacaagatgaaacaattc
ggcattaatacctgagagcaggaagagcaagataaaaggtagtatttgttggcgatcccctag
agtcttttacatcttcggaaaacaaaaactattttttctttaatttctttttttactttctatt
tttaatttatatatttatattaaaaaatttaaattataattatttttatagcacgtgatgaaaa
ggacccaggtggcacttttcggggaaatgtgcgcggaacccctatttgtttatttttctaaata
cattcaaatatgtatccgctcatgagacaataaccctgataaatgcttcaataatattgaaaaa
ggaagagtatgagtattcaacatttccgtgtcgcccttattcccttttttgcggcattttgcct
tcctgtttttgctcacccagaaacgctggtgaaagtaaaagatgctga

**pVJ691 (p316-Dfm1-3HA)**

attctttttttttgatttcggtttccttgaaatttttttgattcggtaatctccgaacagaagga
agaacgaaggaaggagcacagacttagattggtatatatacgcatatgtagtgttgaagaaaca
tgaaattgcccagtattcttaacccaactgcacagaacaaaaacctgcaggaaacgaagataaa
tcatgtcgaaagctacatataaggaacgtgctgctactcatcctagtcctgttgctgccaagct
atttaatatcatgcacgaaaagcaaacaaacttgtgtgcttcattggatgttcgtaccaccaag
gaattactggagttagttgaagcattaggtcccaaaatttgtttactaaaaacacatgtggata
tcttgactgatttttccatggagggcacagttaagccgctaaaggcattatccgccaagtacaa
tttttactcttcgaagacagaaaatttgctgacattggtaatacagtcaaattgcagtactct
gcgggtgtatacagaatagcagaatgggcagacattacgaatgcacacggtgtggtgggcccag
gtattgttagcggtttgaagcaggcggcagaagaagtaacaaaggaacctagaggccttttgat
gttagcagaattgtcatgcaagggctccctatctactggagaatatactaagggtactgttgac
attgcgaagagcgacaaagattttgttatcggctttattgctcaaagagacatgggtggaagag
atgaaggttacgattggttgattatgacacccggtgtgggtttagatgacaagggagacgcatt
gggtcaacagtatagaaccgtggatgatgtggtctctacaggatctgacattattattgttgga
agaggactatttgcaaagggaagggatgctaaggtagagggtgaacgttacagaaaagcaggct
gggaagcatatttgagaagatgcggccagcaaaactaaaaaactgtattataagtaaatgcatg
tatactaaactcacaaattagagcttcaatttaattatatcagttattaccctatgcggtgtga
aataccgcacagatgcgtaaggagaaaataccgcatcaggaaattgtaagcgttaatattttgt
taaaattcgcgttaaatttttgttaaatcagctcattttttaaccaataggccgaaatcggcaa
aatcccttataaatcaaaagaatagaccgagatagggttgagtgttgttccagtttggaacaag
agtccactattaaagaacgtggactccaacgtcaaagggcgaaaaaccgtctatcagggcgatg
gcccactacgtgaaccatcaccctaatcaagttttttggggtcgaggtgccgtaaagcactaaa
tcggaaccctaaagggagcccccgatttagagcttgacggggaaagccggcgaacgtggcgaga
aaggaagggaagaaagcgaaaggagcgggcgctagggcgctggcaagtgtagcggtcacgctgc
gcgtaaccaccacacccgccgcgcttaatgcgccgctacagggcgcgtccattcgccattcagg
ctgcgcaactgttgggaagggcgatcggtgcgggcctcttcgctattacgccagctggcgaaag
ggggatgtgctgcaaggcgattaagttgggtaacgccagggttttcccagtcacgacgttgtaa
aacgacggccagtgaattgtaatacgactcactatagggcgaattggagctccaccgcggtggc
ggccgcagctttctcttgtgaaagccagtcagataatccaaacgagaatccttgtcgaacgtca
cctcatcggtaccgaatttctttgattgctttgtagcatagtttttgcctcttgtaaggatttg
cctattggtgtgaacagccatagcaaaaatcgaaaacctttaattcacccctcttgattcgagc
agctaccaggaacttgaatattgcaactctctatttttatctgctattatctgctgctttgca
gaatcgatcctcatcacatttgaaaaatttcaaatgtgaaaagtccctacttacccgttcgcgg
ctcatgaaatcgtgtcctcttcaatataagacaagagtatcaaacaaaggatataaggcaaaat
acaacaaattaccagtgaattccgcatttccaagtcaaatcaaaaactattttcgaggaaatag
gatccatggcaggcccaaggaatgtgcgtacattacatggaaatggaggccgtaataatgatgt
gatgggcccaaaagaattctggttaaatatcccccaataacaagaactttatttactctagcg
atcgttatgacgatcgtcggccggttgaatctaatcaatccgtggtactttatttatgtttgga
atttgacgttcaagaaggttcagatatggagacttcttacttcttgtgtaatgctttcgtctcg

tgccatgcctgcgctaatggaactatatagtatttatgacaggtcctcacagttggagcgaggg
cattttggtcccggtttgtccaatagacgaggaccgatggtaacagtagactatgcctactacc
tgtgcttttgtatactagccatcactacggccactacaatcatttatgggtcctactaccctgt
tgtgctcacctctgggttcatatcatgcattacatatacttggtcgatcgacaatgccaacgtg
cagatcatgttttatggtttgattcctgtttgggggaagtacttccccttaattcaattgttta
tctctttcgtttttaatgagggtgatttcgtaatctcattgatcggttttactacagggtacct
ttatacatgcttagatacccatacactggggccaatatgggggatgatatccagaaaagctgat
ccaacctacggaatctcgcccaatgggaaattctcaacgccatggtggtttactagtctatatg
ctcgcataacgggtgcccacaatgaaactgcaacattcaacaacaactttgccaatgtgccatc
ttcgcaaagagaaacgagaacttttagtggaagaggtcaacgattgggtacggcccctgcaaca
ttgtctcaaaccagtggcacagattcaggcagagcttctggaagtcaattaagaagtggcccat
cgaatttgaaccaatttcaaggccgcggccaacgagtaggacaaacaaacagcccctccgactc
acaaggccgcatcttttacccatacgatgttcctgactatgcgggctatccctatgacgtcccg
gactatgcaggatcctatccatatgacgttccagattacgctgctcagtgcggctagtaatgac
ctcgagggggggcccggtaccagcttttgttccctttagtgagggttaatttcgagcttggcgt
aatcatggtcatagctgtttcctgtgtgaaattgttatccgctcacaattccacacaacatacg
agccggaagcataaagtgtaaagcctggggtgcctaatgagtgagctaactcacattaattgcg
ttgcgctcactgcccgctttccagtcgggaaacctgtcgtgccagctgcattaatgaatcggcc
aacgcgcggggagaggcggtttgcgtattgggcgctcttccgcttcctcgctcactgactcgct
gcgctcggtcgttcggctgcggcgagcggtatcagctcactcaaaggcggtaatacggttatcc
acagaatcaggggataacgcaggaaagaacatgtgagcaaaaggccagcaaaaggccaggaacc
gtaaaaaggccgcgttgctggcgtttttccataggctccgcccccctgacgagcatcacaaaaa
tcgacgctcaagtcagaggtggcgaaacccgacaggactataaagataccaggcgtttccccct
ggaagctccctcgtgcgctctcctgttccgaccctgccgcttaccggatacctgtccgcctttc
tcccttcgggaagcgtggcgctttctcatagctcacgctgtaggtatctcagttcggtgtaggt
cgttcgctccaagctgggctgtgtgcacgaaccccccgttcagcccgaccgctgcgccttatcc
ggtaactatcgtcttgagtccaacccggtaagacacgacttatcgccactggcagcagccactg
gtaacaggattagcagagcgaggtatgtaggcggtgctacagagttcttgaagtggtggcctaa
ctacggctacactagaagaacagtatttggtatctgcgctctgctgaagccagttaccttcgga
aaaagagttggtagctcttgatccggcaaacaaaccaccgctggtagcggtggtttttttgttt
gcaagcagcagattacgcgcagaaaaaaaggatctcaagaagatcctttgatcttttctacggg
gtctgacgctcagtggaacgaaaactcacgttaagggattttggtcatgagattatcaaaaagg
atcttcacctagatccttttaaattaaaaatgaagttttaaatcaatctaaagtatatatgagt
aaacttggtctgacagttaccaatgcttaatcagtgaggcacctatctcagcgatctgtctatt
tcgttcatccatagttgcctgactccccgtcgtgtagataactacgatacgggagggcttacca
tctggccccagtgctgcaatgataccgcgagacccacgctcaccggctccagatttatcagcaa
taaaccagccagccggaagggccgagcgcagaagtggtcctgcaactttatccgcctccatcca
gtctattaattgttgccgggaagctagagtaagtagttcgccagttaatagtttgcgcaacgtt
gttgccattgctacaggcatcgtggtgtcacgctcgtcgtttggtatggcttcattcagctccg
gttcccaacgatcaaggcgagttacatgatcccccatgttgtgcaaaaaagcggttagctcctt
cggtcctccgatcgttgtcagaagtaagttggccgcagtgttatcactcatggttatggcagca

ctgcataattctcttactgtcatgccatccgtaagatgcttttctgtgactggtgagtactcaa
ccaagtcattctgagaatagtgtatgcggcgaccgagttgctcttgcccggcgtcaatacggga
taataccgcgccacatagcagaactttaaaagtgctcatcattggaaaacgttcttcggggcga
aaactctcaaggatcttaccgctgttgagatccagttcgatgtaacccactcgtgcacccaact
gatcttcagcatcttttactttcaccagcgtttctgggtgagcaaaaacaggaaggcaaaatgc
cgcaaaaaagggaataagggcgacacggaaatgttgaatactcatactcttcctttttcaatat
tattgaagcatttatcagggttattgtctcatgagcggatacatatttgaatgtatttagaaaa
ataaacaaataggggttccgcgcacatttccccgaaaagtgccacctgggtccttttcatcacg
tgctataaaaataattataatttaaattttttaatataaatatataaattaaaaatagaaagta
aaaaaagaaattaaagaaaaaatagtttttgttttccgaagatgtaaaagactctagggggatc
gccaacaaatactaccttttatcttgctcttcctgctctcaggtattaatgccgaattgtttca
tcttgtctgtgtagaagaccacacacgaaaatcctgtgattttacattttacttatcgttaatc
gaatgtatatctatttaatctgcttttcttgtctaataaatatatatgtaaagtacgcttttg
ttgaaattttttaaacctttgtttatttttttttcttcattccgtaactcttctaccttcttta
tttactttctaaaatccaaatacaaaacataaaaataaataaacacagagtaaattcccaaatt
attccatcattaaaagatacgaggcgcgtgtaagttacaggcaagcgatccgtctaagaaacca
ttattatcatgacattaacctataaaaataggcgtatcacgaggccctttcgtctcgcgcgttt
cggtgatgacggtgaaaacctctgacacatgcagctcccggagacggtcacagcttgtctgtaa
gcggatgccgggagcagacaagcccgtcagggcgcgtcagcgggtgttggcgggtgtcggggct
ggcttaactatgcggcatcagagcagattgtactgagagtgcaccataccaccttttcaattca
tcattttttttt

**pVJ704 (p316-Dfm1)**

gaagatcctttgatcttttctacggggtctgacgctcagtggaacgaaaactcacgttaaggga
ttttggtcatgagattatcaaaaaggatcttcacctagatccttttaaattaaaaatgaagttt
taaatcaatctaaagtatatatgagtaaacttggtctgacagttaccaatgcttaatcagtgag
gcacctatctcagcgatctgtctatttcgttcatccatagttgcctgactccccgtcgtgtaga
taactacgatacgggagggcttaccatctggccccagtgctgcaatgataccgcgagacccacg
ctcaccggctccagatttatcagcaataaaccagccagccggaagggccgagcgcagaagtggt
cctgcaactttatccgcctccatccagtctattaattgttgccgggaagctagagtaagtagtt
cgccagttaatagtttgcgcaacgttgttgccattgctacaggcatcgtggtgtcacgctcgtc
gtttggtatggcttcattcagctccggttcccaacgatcaaggcgagttacatgatcccccatg
ttgtgcaaaaaagcggttagctccttcggtcctccgatcgttgtcagaagtaagttggccgcag
tgttatcactcatggttatggcagcactgcataattctcttactgtcatgccatccgtaagatg
cttttctgtgactggtgagtactcaaccaagtcattctgagaatagtgtatgcggcgaccgagt
tgctcttgcccggcgtcaatacgggataataccgcgccacatagcagaactttaaaagtgctca
tcattggaaaacgttcttcggggcgaaaactctcaaggatcttaccgctgttgagatccagttc
gatgtaacccactcgtgcacccaactgatcttcagcatcttttactttcaccagcgtttctggg
tgagcaaaaacaggaaggcaaaatgccgcaaaaaagggaataagggcgacacggaaatgttgaa
tactcatactcttcctttttcaatattattgaagcatttatcagggttattgtctcatgagcgg
atacatatttgaatgtatttagaaaaataaacaaataggggttccgcgcacatttccccgaaaa
gtgccacctgggtccttttcatcacgtgctataaaaataattataatttaaattttttaatata
aatatataaattaaaaatagaaagtaaaaaaagaaattaaagaaaaaatagtttttgttttccg
aagatgtaaaagactctagggggatcgccaacaaatactaccttttatcttgctcttcctgctc
tcaggtattaatgccgaattgtttcatcttgtctgtgtagaagaccacacacgaaaatcctgtg
attttacattttacttatcgttaatcgaatgtatatctatttaatctgcttttcttgtctaata
aatatatatgtaaagtacgctttttgttgaaattttttaaacctttgtttatttttttttcttc
attccgtaactcttctaccttctttatttactttctaaaatccaaatacaaaacataaaaataa
ataaacacagagtaaattcccaaattattccatcattaaaagatacgaggcgcgtgtaagttac
aggcaagcgatccgtctaagaaaccattattatcatgacattaacctataaaaataggcgtatc
acgaggccctttcgtctcgcgcgtttcggtgatgacggtgaaaacctctgacacatgcagctcc
cggagacggtcacagcttgtctgtaagcggatgccgggagcagacaagcccgtcagggcgcgtc
agcgggtgttggcgggtgtcggggctggcttaactatgcggcatcagagcagattgtactgaga
gtgcaccataccaccttttcaattcatcatttttttttttattcttttttttgatttcggtttcc
ttgaaatttttttgattcggtaatctccgaacagaaggaagaacgaaggaaggagcacagactt
agattggtatatatacgcatatgtagtgttgaagaaacatgaaattgcccagtattcttaaccc
aactgcacagaacaaaaacctgcaggaaacgaagataaatcatgtcgaaagctacatataagga
acgtgctgctactcatcctagtcctgttgctgccaagctatttaatatcatgcacgaaaagcaa
acaaacttgtgtgcttcattggatgttcgtaccaccaaggaattactggagttagttgaagcat
taggtcccaaaatttgtttactaaaaacacatgtggatatcttgactgatttttccatggaggg
cacagttaagccgctaaaggcattatccgccaagtacaatttttactcttcgaagacagaaaa
tttgctgacattggtaatacagtcaaattgcagtactctgcgggtgtatacagaatagcagaat

gggcagacattacgaatgcacacggtgtggtgggcccaggtattgttagcggtttgaagcaggc
ggcagaagaagtaacaaaggaacctagaggccttttgatgttagcagaattgtcatgcaagggc
tccctatctactggagaatatactaagggtactgttgacattgcgaagagcgacaaagatttg
ttatcggctttattgctcaaagagacatgggtggaagagatgaaggttacgattggttgattat
gacacccggtgtgggtttagatgacaagggagacgcattgggtcaacagtatagaaccgtggat
gatgtggtctctacaggatctgacattattattgttggaagaggactatttgcaaagggaaggg
atgctaaggtagagggtgaacgttacagaaaagcaggctgggaagcatatttgagaagatgcgg
ccagcaaaactaaaaaactgtattataagtaaatgcatgtatactaaactcacaaattagagct
tcaatttaattatatcagttattaccctatgcggtgtgaaataccgcacagatgcgtaaggaga
aaataccgcatcaggaaattgtaagcgttaatattttgttaaaattcgcgttaaatttttgtta
aatcagctcattttttaaccaataggccgaaatcggcaaaatcccttataaatcaaaagaatag
accgagatagggttgagtgttgttccagtttggaacaagagtccactattaaagaacgtggact
ccaacgtcaaagggcgaaaaaccgtctatcagggcgatggcccactacgtgaaccatcaccta
atcaagttttttggggtcgaggtgccgtaaagcactaaatcggaaccctaaagggagcccccga
tttagagcttgacggggaaagccggcgaacgtggcgagaaaggaagggaagaaagcgaaaggag
cgggcgctagggcgctggcaagtgtagcggtcacgctgcgcgtaaccaccacacccgccgcgct
taatgcgccgctacagggcgcgtccattcgccattcaggctgcgcaactgttgggaagggcgat
cggtgcgggcctcttcgctattacgccagctggcgaaagggggatgtgctgcaaggcgattaag
ttgggtaacgccagggttttcccagtcacgacgttgtaaaacgacggccagtgaattgtaatac
gactcactatagggcgaattggagctccaccgcggtggcggccgcagctttctcttgtgaaagc
cagtcagataatccaaacgagaatccttgtcgaacgtcacctcatcggtaccgaatttctttga
ttgctttgtagcatagtttttgcctcttgtaaggatttgcctattggtgtgaacagccatagca
aaaatcgaaaacctttaattcacccctcttgattcgagcagctaccaggaacttgaatattgca
actctctatttttatctgctattatctgctgctttgcagaatcgatcctcatcacatttgaaa
aatttcaaatgtgaaaagtccctacttacccgttcgcggctcatgaaatcgtgtcctcttcaat
ataagacaagagtatcaaacaaaggatataaggcaaaatacaacaaattaccagtgaattccgc
atttccaagtcaaatcaaaaactattttcgaggaaataggatccatggcaggcccaaggaatgt
gcgtacattacatggaaatggaggccgtaataatgatgtgatgggcccaaaagaattctggtta
aatatcccccaataacaagaactttatttactctagcgatcgttatgacgatcgtcggccggt
tgaatctaatcaatccgtggtactttatttatgtttggaatttgacgttcaagaaggttcagat
atggagacttcttacttcttgtgtaatgctttcgtctcgtgccatgcctgcgctaatggaacta
tatagtatttatgacaggtcctcacagttggagcgagggcattttggtcccggtttgtccaata
gacgaggaccgatggtaacagtagactatgcctactacctgtgcttttgtatactagccatcac
tacggccactacaatcatttatgggtcctactaccctgttgtgctcacctctgggttcatatca
tgcattacatatacttggtcgatcgacaatgccaacgtgcagatcatgttttatggtttgattc
ctgtttgggggaagtacttccccttaattcaattgtttatctctttcgtttttaatgagggtga
tttcgtaatctcattgatcggttttactacagggtacctttatacatgcttagatacccataca
ctggggccaatatgggggatgatatccagaaaagctgatccaacctacggaatctcgcccaatg
ggaaattctcaacgccatggtggtttactagtctatatgctcgcataacgggtgcccacaatga
aactgcaacattcaacaacaactttgccaatgtgccatcttcgcaaagagaaacgagaactttt
agtggaagaggtcaacgattgggtacggcccctgcaacattgtctcaaaccagtggcacagatt

caggcagagcttctggaagtcaattaagaagtggcccatcgaatttgaaccaatttcaaggccg
cggccaacgagtaggacaaacaaacagcccctccgactcacaataactcgagggggggcccggt
accagcttttgttccctttagtgagggttaatttcgagcttggcgtaatcatggtcatagctgt
ttcctgtgtgaaattgttatccgctcacaattccacacaacatacgagccggaagcataaagtg
taaagcctggggtgcctaatgagtgagctaactcacattaattgcgttgcgctcactgcccgct
ttccagtcgggaaacctgtcgtgccagctgcattaatgaatcggccaacgcgcggggagaggcg
gtttgcgtattgggcgctcttccgcttcctcgctcactgactcgctgcgctcggtcgttcggct
gcggcgagcggtatcagctcactcaaaggcggtaatacggttatccacagaatcaggggataac
gcaggaaagaacatgtgagcaaaaggccagcaaaaggccaggaaccgtaaaaaggccgcgttgc
tggcgtttttccataggctccgcccccctgacgagcatcacaaaaatcgacgctcaagtcagag
gtggcgaaacccgacaggactataaagataccaggcgtttccccctggaagctccctcgtgcgc
tctcctgttccgaccctgccgcttaccggatacctgtccgcctttctcccttcgggaagcgtgg
cgctttctcatagctcacgctgtaggtatctcagttcggtgtaggtcgttcgctccaagctggg
ctgtgtgcacgaaccccccgttcagcccgaccgctgcgccttatccggtaactatcgtcttgag
tccaacccggtaagacacgacttatcgccactggcagcagccactggtaacaggattagcagag
cgaggtatgtaggcggtgctacagagttcttgaagtggtggcctaactacggctacactagaag
aacagtatttggtatctgcgctctgctgaagccagttaccttcggaaaaagagttggtagctct
tgatccggcaaacaaaccaccgctggtagcggtggtttttttgtttgcaagcagcagattacgc
gcagaaaaaaaggatctcaa
